# HIV-1 neutralizing antibodies in SHIV-infected macaques recapitulate structurally divergent modes of human V2 apex recognition with a single D gene

**DOI:** 10.1101/2024.06.11.598384

**Authors:** Ryan S. Roark, Rumi Habib, Jason Gorman, Hui Li, Andrew Jesse Connell, Mattia Bonsignori, Yicheng Guo, Michael P. Hogarty, Adam S. Olia, Kirsten Sowers, Baoshan Zhang, Frederic Bibollet-Ruche, Sean Callaghan, John W. Carey, Gabriele Cerutti, Darcy R. Harris, Wanting He, Emily Lewis, Tracy Liu, Rosemarie D. Mason, Younghoon Park, Juliette M. Rando, Ajay Singh, Jeremy Wolff, Q. Paula Lei, Mark K. Louder, Nicole A. Doria-Rose, Raiees Andrabi, Kevin O. Saunders, Michael S. Seaman, Barton F. Haynes, Daniel W. Kulp, John R. Mascola, Mario Roederer, Zizhang Sheng, Beatrice H. Hahn, George M. Shaw, Peter D. Kwong, Lawrence Shapiro

## Abstract

Broadly neutralizing antibodies targeting the V2 apex of the HIV-1 envelope trimer are among the most common specificities elicited in HIV-1-infected humans and simian-human immunodeficiency virus (SHIV)-infected macaques. To gain insight into the prevalent induction of these antibodies, we isolated and characterized 11 V2 apex-directed neutralizing antibody lineages from SHIV-infected rhesus macaques. Remarkably, all SHIV-induced V2 apex lineages were derived from reading frame two of the rhesus DH3-15*01 gene. Cryo-EM structures of envelope trimers in complex with antibodies from nine rhesus lineages revealed modes of recognition that mimicked three canonical human V2 apex-recognition modes. Notably, amino acids encoded by DH3-15*01 played divergent structural roles, inserting into a hole at the trimer apex, H-bonding to an exposed strand, or forming part of a loop scaffold. Overall, we identify a DH3-15*01-signature for rhesus V2 apex broadly neutralizing antibodies and show that highly selected genetic elements can play multiple roles in antigen recognition.

**Highlights:**

- Isolated 11 V2 apex-targeted HIV-neutralizing lineages from 10 SHIV-infected Indian-origin rhesus macaques
- Cryo-EM structures of Fab-Env complexes for nine rhesus lineages reveal modes of recognition that mimic three modes of human V2 apex antibody recognition
- All SHIV-elicited V2 apex lineages, including two others previously published, derive from the same DH3-15*01 gene utilizing reading frame two
- The DH3-15*01 gene in reading frame two provides a necessary, but not sufficient, signature for V2 apex-directed broadly neutralizing antibodies
- Structural roles played by DH3-15*01-encoded amino acids differed substantially in different lineages, even for those with the same recognition mode
- Propose that the anionic, aromatic, and extended character of DH3-15*01 in reading frame two provides a selective advantage for V2 apex recognition compared to B cells derived from other D genes in the naïve rhesus repertoire
- Demonstrate that highly selected genetic elements can play multiple roles in antigen recognition, providing a structural means to enhance recognition diversity

## Introduction

Antibodies directed to the V2 apex of the HIV-1 envelope (Env) trimer represent one of the most prevalent categories of broadly neutralizing antibodies elicited by natural infection^1–5^. These antibodies are characterized by heavy chain third complementarity determining regions (HCDR3s) that are exceptionally long and anionic, and often tyrosine sulfated^6,7^. V2 apex-directed antibodies generally have less affinity maturation than other categories of HIV-1 broadly neutralizing antibodies, with longitudinal analyses indicating that cross-clade neutralization can be achieved rapidly, in some cases within a few weeks or months after initial B cell activation^8,9^.

The prevalence of appropriate HCDR3s within the naïve B cell population that might serve as precursors to V2 apex-directed broadly neutralizing antibodies has been investigated in humans. Two factors appear critical: (i) generation of long, negatively-charged HCDR3s by recombination, and (ii) placement of appropriate HCDR3s into a context that enables V2 apex recognition^10–12^. For preclinical vaccine development, however, such HCDR3s need to be generated in standard vaccine-test species. Long HCDR3s are extremely rare in mice, rats, and guinea pigs, which are often used to assess vaccine immunogens, whereas non-human primates can generate long HCDR3s^13–16^ and may represent a more appropriate vaccine-test species.

Previously, we observed V2 apex-directed antibodies to be the most common broadly neutralizing antibody specificity elicited in chimeric simian-human immunodeficiency virus (SHIV)-infected rhesus macaques^17^ (HL and GMS, personal communication). In that study, we demonstrated the induction of a V2 apex-directed lineage, RHA1, that exhibited ∼50% neutralization breadth and a PGT145-like mode of epitope recognition^9,18^. Here, we conducted a systematic analysis of the immunogenetics and structures of a repertoire of rhesus V2 apex-targeted neutralizing antibodies isolated from ten SHIV-infected rhesus macaques to gain insight into their prevalence and to assess the relevance of the rhesus model for V2 apex-targeted HIV-1 vaccine design and testing. We used antigen-specific single-cell sorting to isolate 11 new rhesus V2 apex-directed antibody lineages and characterized these lineages for their immunogenetics, structures, modes of epitope recognition, and neutralization phenotype. For structural characterization, we used single particle cryo-EM analysis of complexes between the antigen-binding fragments of antibodies and Env trimers, stabilized in the prefusion-closed conformation. Remarkably, all rhesus V2 apex-directed antibodies utilized the DH3-15*01 gene in reading frame two, which unexpectedly adopted divergent structural motifs and diverse modes of paratope-epitope binding. Overall, our results demonstrate how structural diversity and unique biochemical properties of the DH3-15*01 gene can yield highly advantageous rhesus-specific paratopes for V2 apex recognition that recapitulate mode-defining features of human broadly neutralizing antibodies.

## Results

### Identification of diverse V2 apex-targeted neutralizing lineages from ten SHIV-infected rhesus macaques

To characterize the molecular repertoire of rhesus V2 apex recognition, we used antigen-specific single-cell sorting to isolate monoclonal antibodies from ten SHIV-infected rhesus macaques with polyclonal V2 apex-directed broadly neutralizing responses (**Figure 1A, S1**). These animals were infected by SHIVs expressing one of five HIV-1 subtype A or C Envs, which elicited measurable heterologous tier-2 neutralizing antibody responses as early as 4 weeks and as late as 88 weeks post-infection (**Table S1**). Memory B cells were sorted from peripheral blood mononuclear cells of each rhesus macaque from a single timepoint using heterologous or epitope-specific Env SOSIP-probe pairs (**Table S2**). Pairings comprised (i) a heterologous Env trimer with different fluorophores, (ii) two different heterologous Env trimers, or (iii) an Env trimer paired with a V2 apex epitope mutant trimer. By using PCR to amplify the paired immunoglobin variable genes from single-cell transcripts^19,20^, we recovered monoclonal antibodies that belonged to individual expanded lineages from each rhesus macaque, with two distinct lineages from RM42056, each bearing atypically long ≥23 residue HCDR3s (IMGT numbering) with an overall electronegative charge due to an enrichment of anionic residues (Glu [E] and Asp [D]) (antibodies named to indicate rhesus ID-lineage.clone with a representative monoclonal antibody from each lineage included in Figure 1) (**Figure 1B, Table S2**). The HCDR3 features of the rhesus antibodies shown in **Figure 1B** are characteristic of human and rhesus broadly neutralizing antibodies that target the HIV-1 V2 apex epitope and must penetrate apical glycans to reach the shielded cationic C-strand underneath^4,6,7,21^.

**Figure 1.**
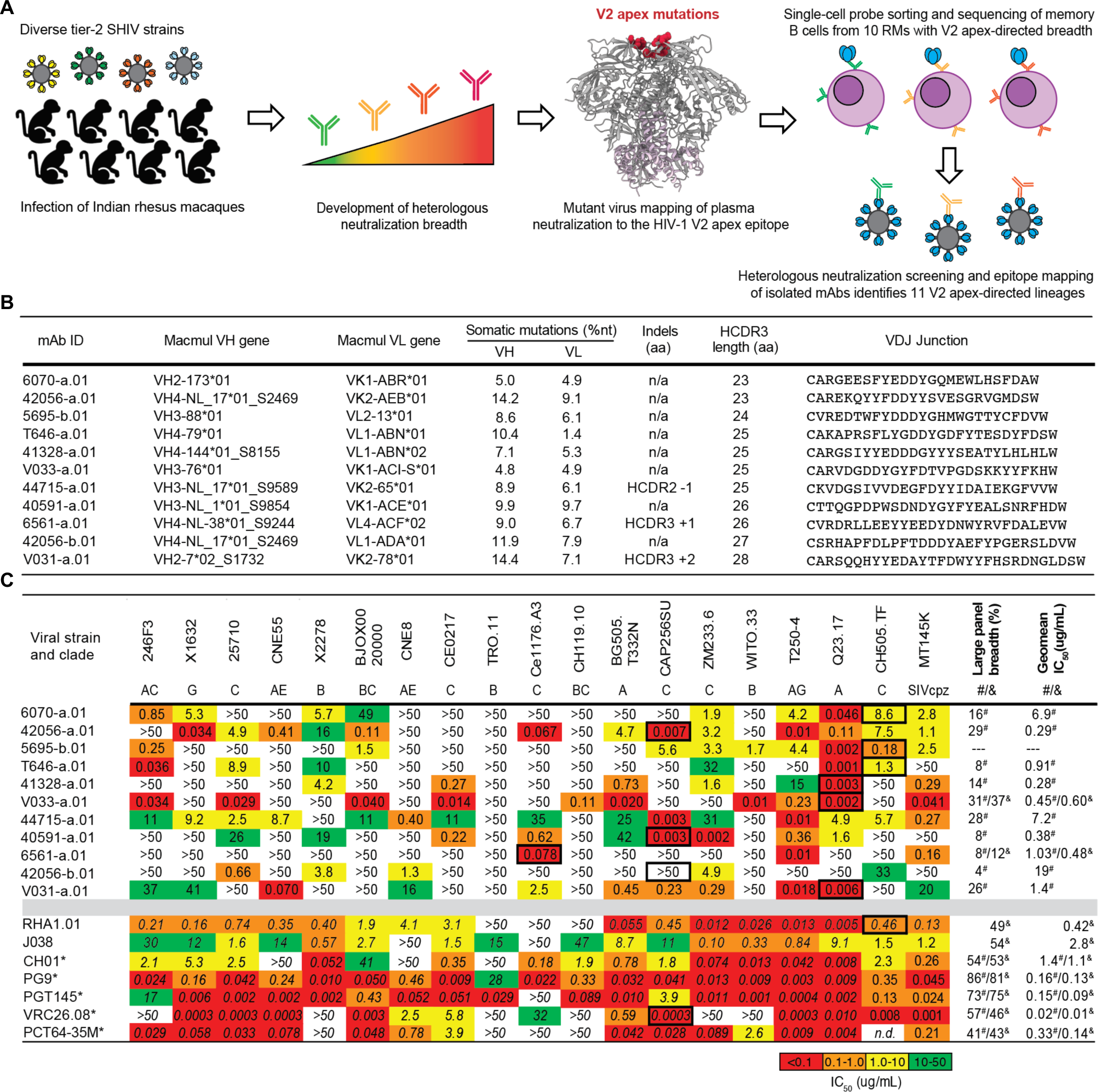
Single-cell sorting identifies broadly neutralizing lineages from 10 rhesus macaques with HIV-1 V2 apex-directed heterologous neutralization breadth. (A) Schematic for the present study. (B) Immunogenetics of a representative monoclonal antibody from each of the 11 rhesus lineages that are reported in this study. (C) Neutralization breadth and potency of representative lineage antibodies reported here. Left, neutralization activity against a 19-member panel of cross-clade tier-2 HIV-1 strains and a simian immunodeficiency virus infecting chimpanzees (SIVcpz), MT145K, which has been shown to bear the conserved HIV-1 V2 apex epitope^60,73^. Data is reported as IC50 titer (ug/mL) and colored according to the legend. Bold boxes demarcate activity against autologous virus; for example, 6070-a.01 was isolated from an animal infected with a SHIV bearing the HIV-1 CH505.TF Env (SHIV.CH505). Right, neutralization breadth and geomean IC50 against one or two large cross-clade panels of HIV-1 strains. # denotes activity against 119 viruses (Seaman panel); & denotes activity against 208 viruses (VRC panel). Bottom, previously published rhesus and human V2 apex broadly neutralizing antibodies are included below the gray row for comparison; human antibodies are denoted with *. IC50 data for these antibodies are shown in italics when obtained from their respective publications. 119 virus panel data (#) for CH01, VRC26.08, and PCT64-35M were derived from CATNAP^74^ (https://www.hiv.lanl.gov/components/sequence/HIV/neutralization/).

To accurately assign the germline genes and levels of somatic hypermutation (SHM) for each lineage, we performed next-generation sequencing of naïve B cell transcripts of each rhesus macaque and analyzed these sequences with IgDiscover software^22^ to curate personalized immunoglobulin gene libraries (**Table S3**). The heavy and light chain variable regions of each rhesus lineage were derived from unique recombined genetic origins, although the 42056-a and 42056-b lineages utilized the same germline VH4-NL_17*01_S2469 gene and the T646-a and 41328-a lineages utilized similar alleles of the VL1-ABN gene (**Figure 1B**, **Table S3**). SHM within all lineages was modest, with nucleotide divergence from each respective germline V gene ranging from 2-16% in heavy chain and 3-10% in light chain across all recovered antibodies. Only 3 of the 11 lineages contained insertions or deletions (indels) compared with germline: a single residue deletion within HCDR2 of the 44715-a lineage, a two-residue insertion in the HCDR3 of the V031-a lineage, and a single residue insertion within HCDR3 of the 6561-a.01 lineage. These SHM features are consistent with previously described human and rhesus V2 apex-directed broadly neutralizing antibodies, which typically require less affinity maturation and fewer or no indels to achieve breadth^8,9,23–25^ compared with human broadly neutralizing antibodies targeting other sites of vulnerability^26–29^.

We then synthesized and expressed antibodies from each lineage and tested them for neutralization against a panel of 19 tier-2 neutralization-resistant viruses (**Figure 1C**, **Table S4**). The lineages exhibited a range of activity, with the broadest members of each lineage neutralizing 11% to 78% of heterologous viruses with a median IC50 ranging from 0.04 to 8.7 ug/mL (**Figure 1C**). Most of these lineages were subsequently tested for neutralization against large 119 or 208 member panels of diverse HIV-1 strains and found to exhibit breadth of 4-37% with geomean IC50 of 0.28-19.8 ug/mL (**Figure 1C, Table SX**). The V033-a lineage was notable because it was isolated just 24 weeks post-SHIV infection, exhibited particularly low levels of SHM (2.0 – 6.8% VH nucleotide), and neutralized 31 and 37% of the 119 and 208 member cross-clade panels with geomean IC50sof 0.45 and 0.60 ug/mL, respectively; this highlights the rapidity by which potent V2 apex-directed broadly neutralizing antibodies can develop once an appropriate naïve B cell precursor is primed and activated.

A single monoclonal antibody from 8 of the 11 lineages was able to recapitulate most of the heterologous neutralization in each respective macaque plasma, suggesting that a single lineage was responsible for neutralization breadth in a majority of animals (**Figure S2A**). There were three exceptions: First was 5695-b isolated from SHIV.CH505-infected RM5695, which was more narrow in breadth, while the RHA1 broadly neutralizing lineage that was also isolated from RM5695^17^ largely recapitulated the the animal’s plasma breadth. Second was animal RM6561 from whom we isolated the 6561-a lineage, which targeted the V2 apex; a second broadly neutralizing lineage targeting the fusion peptide was also isolated from this animal (GMS, personal communication). Lastly, RM44715, from whom we isolated the V2 apex-targeted lineage 44715-a, also harbored a second neutralizing antibody lineage that targeted the V3-glycan supersite (GMS, personal communication). The lineages targeting two distinct epitopes in RMs 6561 and 44715 were together responsible for plasma breadth.

To map phenotypically the epitope specificity of the newly identified lineages, we tested representative lineage members for neutralization against heterologous viruses bearing mutations at canonical V2 apex epitope residues 160, 166, 169 and 171 (**Figure S2B**). Removal of the N160 glycan or substitutions of positively-charged residues abrogated or substantially reduced neutralization of these mutant heterologous viruses. Based on patterns of neutralization loss against viruses containing different canonical V2 apex mutations, we could divide the rhesus antibody lineages into five distinct phenotypic groups (**Figure S2B, S2D**): Three of these groups shared similar patterns with the three prototypic modes of HCDR3-dominated human V2 apex broadly neutralizing antibodies^6,8^, while a fourth and fifth group exhibited novel phenotypes. The fourth group, comprising lineages 6070-a and T646-a, was distinguished by dramatic enhancement in neutralization potency (100 to 10,000-fold) against heterologous viruses lacking N160 glycan; this stands in contrast to previously reported V2 apex broadly neutralizing antibodies which generally are N160 dependent. The fifth group included the V033-a lineage, which exhibited variable strain-specific dependence on N160 glycan for neutralization (**Figure S2C**), like the human antibody VRC26.25. However, unlike VRC26.25, the V033-a lineage was not affected by mutations at residue 166 within the apex hole ^30^. Altogether, characterization of the 11 new V2 apex-targeted broadly neutralizing lineages from SHIV-infected rhesus macaques revealed antibodies that shared many immunogenetic and phenotypic features with previously described human V2 apex broadly neutralizing antibodies, though segregating into five distinct phenotypic groups – two newly identified – based on their sensitivity to specific V2 apex substitutions in sites of paratope-epitope interaction (**Figure S2D**).

### Rhesus V2 apex-targeted lineages all utilize the same DH3-15*01 gene

Our initial immunogenetic analysis revealed that each of the 11 newly identified rhesus lineages were derived from unique heavy and light chain V gene and J gene pairs, but all utilized the same DH3-15*01 gene [alternate designation: DH3-9*01^31,32^] (**Figure 2A, Table S2**). Moreover, previously reported rhesus V2 apex broadly neutralizing antibodies RHA1 and J038 also utilize DH3-15*01^17,33^. We confirmed the presence of the exact germline DH3-15*01 sequence in each of ten rhesus macaques for which we had naïve B cell transcript sequences (**Table S3**). To gain further insight into the utilization of this D gene, we performed VDJ junctional analysis (residues C92H to W103H; Kabat numbering) for all 11 rhesus V2 apex lineages along with the two previously reported lineages. To facilitate and visualize comparisons, we aligned the 13 sets of germline and VDJ junction sequences against DH3-15*01 (**Figure 2B**).

**Figure 2.**
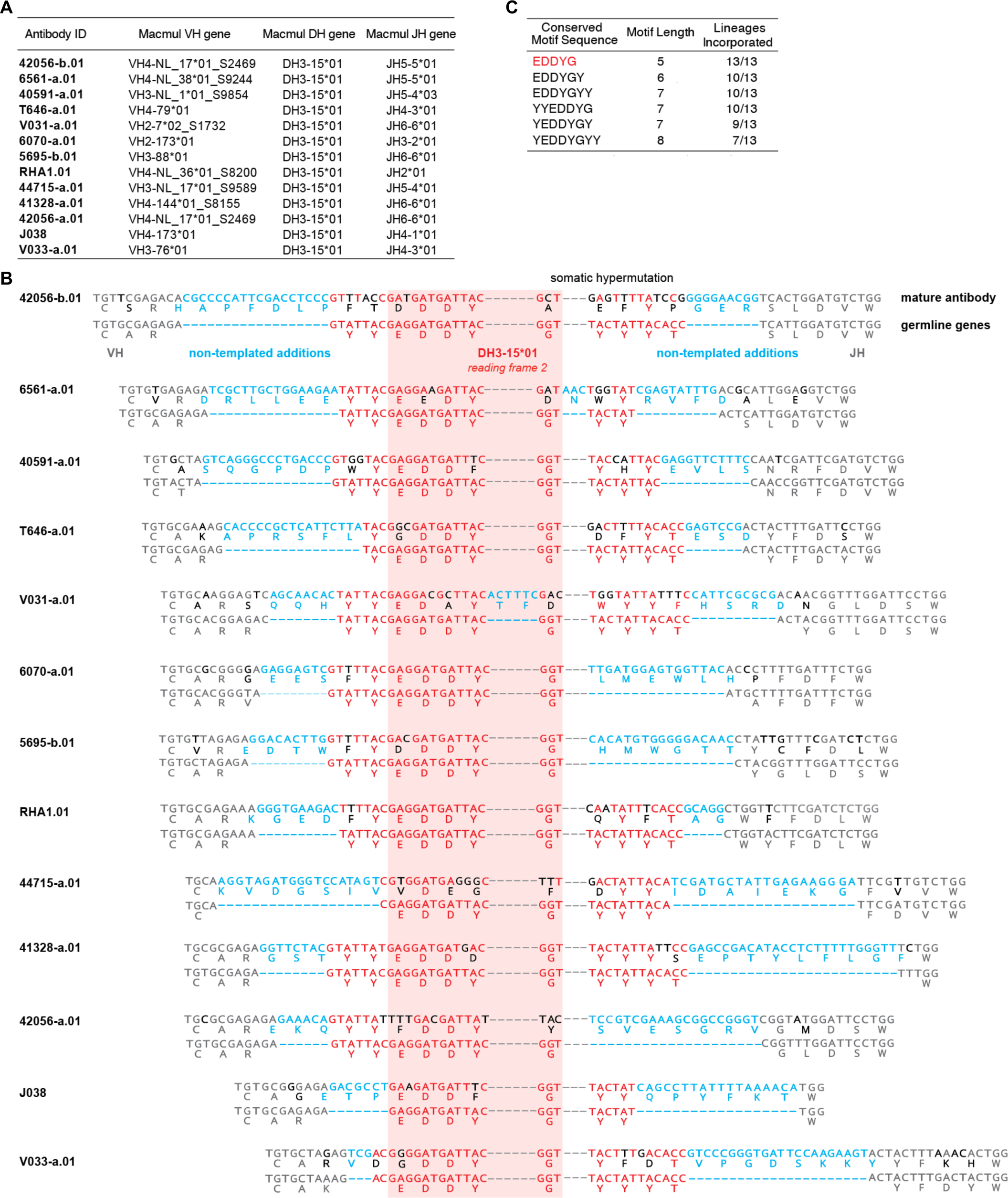
All SHIV-induced V2 apex-directed lineages are derived from the same rhesus DH3-15*01 gene in reading frame two and invariantly acquire a minimal five residue motif. (A) Germline VH, DH and JH genes for all potently neutralizing rhesus V2 apex-directed lineages described here and previously (13 total). (B) VDJ junction analysis of a representative antibody from each rhesus lineage. The respective germline VH, DH, and JH gene sequences are truncated and aligned to each VDJ junction, and each VDJ junction is aligned with respect to the DH gene. VH and JH nucleotides and residues are colored grey, non-templated nucleotides and residues (insertions and N/P additions) are colored blue, DH-gene nucleotides and residues are colored red, and somatic hypermutation is colored black; we do not interpret somatic hypermutation within non-templated regions. The reading frame in which each DH gene has been incorporated is denoted. The five residue DH3-15*01 motif (EDDYG) acquired by all 13 lineages during VDJ recombination is highlighted with transparent red shading. (C) List of conserved DH3-15*01 residue motifs of varying length that are acquired by at least half of the rhesus lineages during VDJ recombination. The five-residue EDDYG motif described in panel (B) is written in red.

Strikingly, DH3-15*01 was invariantly incorporated in reading frame two by each lineage. In addition to being rich in anionic residues, rhesus and human V2 apex broadly neutralizing antibody HCDR3s also contain many aromatic residues (most commonly Tyr [Y]) (**Figure 1B, Table S2**)^8,17,23–25^. A majority of these characteristic residues in rhesus lineages were contributed by DH3-15*01, which could only be achieved by translation in the second reading frame. The DH3-15*01 start positions (the first residue fully coded by D gene nucleotides) for all rhesus lineages spanned just five residues from 97H to 100bH, with the most common positions 98H and 99H shared by three lineages each. DH3-15*01 added significantly to the atypical length of each HCDR3 by contributing 21 to 33 D gene nucleotides, resulting in a minimal germline 15 nucleotide sequence that was incorporated into all 13 rhesus lineages (highlighted by red shading in the alignment) (**Figure 2B**). This conserved D gene sequence yielded a five-residue EDDYG motif that was shared by each rhesus lineage inferred unmutated common ancestor following VDJ recombination (**Figure 2C**). This motif was not commonly subjected to somatic hypermutation, as most lineages (8 of 13) had zero to one mutated motif residue in mature clonal sequences. Even longer seven-residue motifs, YYEDDYG or EDDYGYY, achieved by including two consecutive germline-encoded Tyr residues at either the N- or C-termini, were observed in 9 of 13 lineages (**Figure 2B, 2C**). Identification of 6561-a and V031-a lineage sequences from timepoints preceding monoclonal antibody isolation confirmed an insertion of one and two residues, respectively, within the original nine-residue D gene fragment during lineage development (**Figure S3**).

Overall, this analysis indicates DH3-15*01 gene usage in reading frame two to be a signature feature of HCDR3 ontogenies in rhesus V2 apex broadly neutralizing lineages. The invariant incorporation of the EDDYG motif during each VDJ recombination event in otherwise genetically diverse templated and non-templated backgrounds suggests this sequence to be important for rhesus antibody recognition of the HIV-1 V2 apex.

### Cryo-EM structures of rhesus antibody lineages identify three reproducible mechanisms of V2 apex recognition

To provide molecular characterization of rhesus V2 apex recognition and the specific role of the DH3-15*01 gene, we determined the structures of antigen-binding fragments (Fabs) from nine of the new rhesus lineages in complex with HIV-1 Env SOSIP trimers using single-particle cryo-EM (**Dataset S1, Table S6**). Structural analysis revealed that these rhesus lineages mimicked defining features of all three modes of human HCDR3-dominated V2 apex broadly neutralizing antibodies^6,7^.

#### PGT145-like recognition

The cryo-EM structures of 6070-a.01, T646-a.01, 42056-a.01, and 44715-a.01 (solved with 3D molecular reconstructions at 3.6 Å, 3.5 Å, 4.2 Å, and 3.9 Å resolution, respectively) revealed modes of recognition similar to antibodies PGT145 and PCT64, which represent an extended human broadly neutralizing antibody class that utilizes a needle-like hole-insertion mechanism to recognize the V2 apex (**Figure 3E**) ^9,18,25,34^. A single Fab of each of these antibodies bound at the Env C3 symmetry axis with extended HCDR3s inserted directly into the cationic trimer hole (**Figure 3A-D**). Each lineage recognized two or more apical glycans from multiple protomers, resulting in 41% to 55% of their respective interactive surface areas being contributed by glycan interfaces (**Table S5**). This was comparable to the glycan fraction of interactive surfaces for PGT145 (45%) and PCT64-35s (56%). 6070-a.01 and T646-a.01 each recognized N160 glycan from all three protomers despite comprising the phenotypic neutralization group enhanced by N160 glycan removal (**Figure S2D**). However, both lineages reoriented one of these glycans outward from the trimer C3 axis (6070-a) or into a horizonal conformation parallel with the Fab combining surface (T646-a) (denoted with * in Figure 3A and 3B); this is in contrast to other human and rhesus PGT145-like antibodies that accommodate N160 glycans in a more vertical conformation similar to their conformation on the unliganded trimer. The induced glycan reorientation to accomodate 6070-a.01 and T646-a.01 is likely a barrier to binding, resulting in the enhanced potency of these antibodies once N160 glycan has been removed.

**Figure 3.**
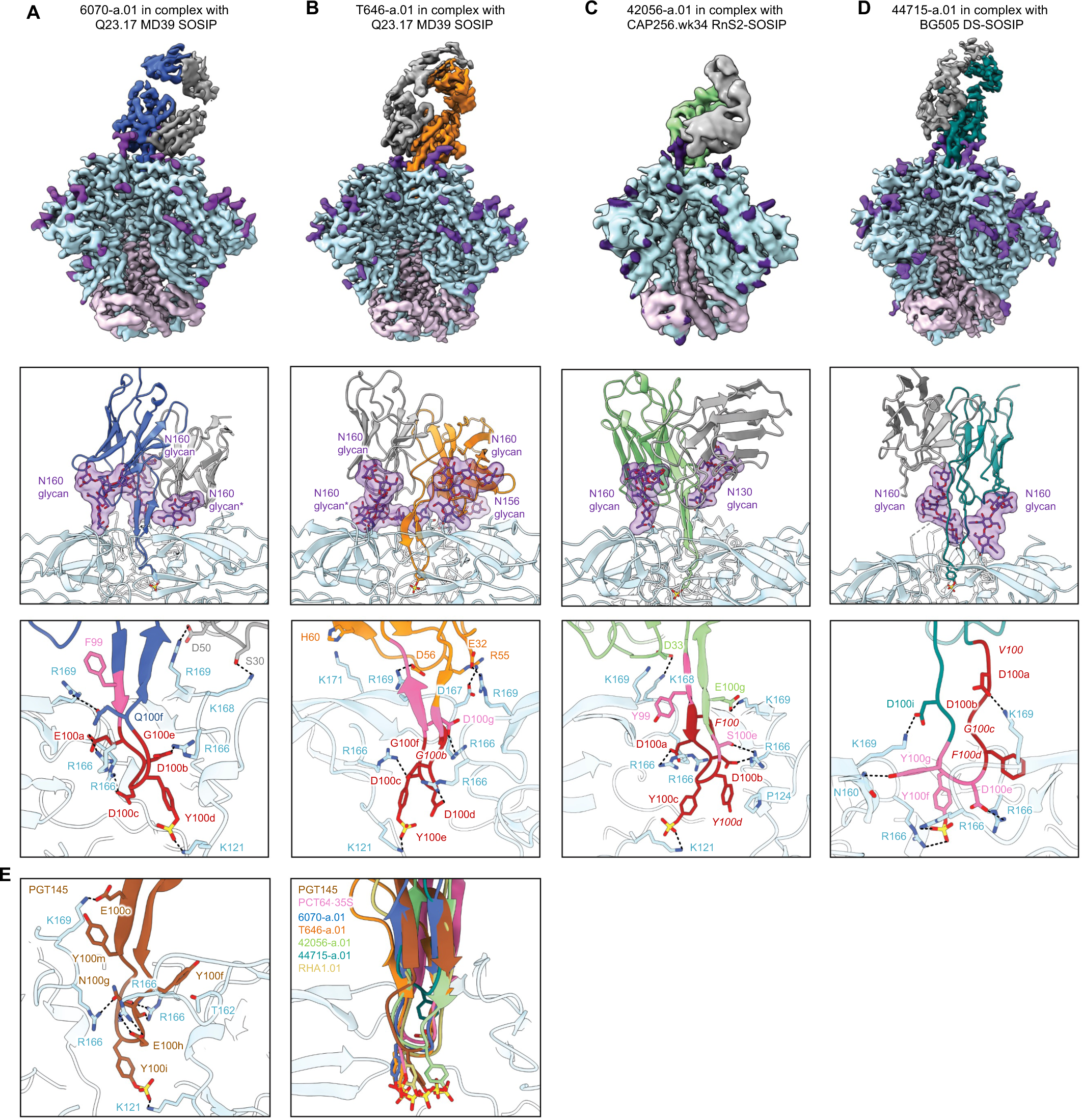
Cryo-EM structures reveal PGT145-like modes of V2 apex recognition to be a reproducible antibody extended-class in rhesus macaques. (A) Top; cryo-EM reconstruction of 6070-a.01 in complex with Q23.17 MD39 SOSIP at 3.6 Å resolution. The 6070-a.01 heavy and light chains are colored blue and gray, respectively. Envelope gp120, gp41, and *N*-linked glycans are colored turquoise, pink, and purple, respectively. Middle; expanded interface view of 6070-a.01 from Top panel to highlight binding position and interactions with apical envelope glycans. Glycans bound by 6070-a.01 are shown in stick representation with transparent surfaces. The N160 glycan reoriented outward and away from the 3-fold trimer axis is denoted with *. Sulfated tyrosine residues are shown in stick representation to highlight their position within the trimer. Bottom; further expanded interface view of 6070-a.01 to highlight interactions with apical envelope residues. Interacting residues are depicted in stick representation. Residues at positions corresponding to the conserved five-residue DH3-15*01 gene motif are colored dark red, while the remaining D gene residues are colored pink. Conserved motif position labels are italicized when subjected to somatic hypermutation. Nitrogen atoms are colored blue, oxygen atoms colored bright red, and sulfur atoms colored yellow. Hydrogen bonds and salt bridges (distance < 3.3 Å) are depicted with dashed lines. (B) Top; cryoEM reconstruction of T646-a.01 in complex with Q23.17 MD39 SOSIP at 3.5 Å resolution. The T646-a.01 heavy chain is colored orange and the remainder of the complex is colored similarly to panel (A). Middle; expanded interface view of T646-a.01 from Top panel to highlight binding position and apical glycan interactions is shown similarly to panel (A), including the N160 glycan reoriented into a horizonal conformation denoted with *. Bottom; further expanded interface view of T646-a.01 to highlight apical residue interactions is shown similarly to panel (A). (C) Top; cryo-EM reconstruction of 42056-a.01 in complex with CAP256.wk34.c80 RnS2 SOSIP determined at 4.1 Å resolution. The 42056-a.01 heavy chain is colored light green and the remainder of the complex is colored similarly to panel (A). Middle; expanded interface view of 42056-a.01 from Top panel to highlight binding position and apical glycan interactions is shown similarly to panel (A). Bottom; further expanded interface view of 42056-a.01 highlight apical residue interactions is shown similarly to panel (A). (D) Top; cryo-EM reconstruction of 44715-a.01 in complex with BG505 DS-SOSIP at 3.9 Å resolution. The 44715-a.01 heavy chain is colored teal and the remainder of the complex is colored similarly to panel (A). Middle; expanded interface view of 44715-a.01 from Top panel to highlight binding position and apical glycan interactions is shown similarly to panel (A). Bottom; further expanded interface view of 44715-a.01 to highlight apical residue interactions is shown similarly to panel (A). (E) Right; Expanded HCDR3 interface view of PGT145 (PDB-5V8L) to highlight apical residue interactions is shown similarly to panel (A). The tyrosine sulfation posttranslational modification of Y100i was not included in this structure and therefore modeled here. Left; expanded HCDR3 interface side view of the alignment of envelope complex structures of 6070-a.01, T646-a.01, 42056-a.01, and 44715-a.01 determined here to envelope complexes with human Fabs PCT64-35S (PDB-7T74) and PGT145 (PDB-5V8L) and rhesus Fab RHA1.V2.01 (PDB-6XRT). Alignments were made with gp120 from each complex. Only gp120 of the 6070-a.01complex is shown for clarity. Sulfated tyrosine residues are shown to highlight their positioning within the trimer.

The four new lineages each interacted with conserved cationic amino acids from all three protomers lining the trimer apex hole, most commonly through electrostatic interactions with Env residues 166 and 169. The 6070-a.01 complex revealed a three-residue anionic motif (E100aHCDR3, D100bHCDR3, D100cHCDR3) to form salt bridges with all three R166 residues, while F100 and Q100fHCDR3 together interacted with R169 from a single protomer and light chain residues S30LCDR1 and D50LCDR2 engaged K168 and R169 from a second protomer (**Figure 3A, bottom**). T646-a.01 similarly formed salt bridges with R166 from all three protomers mediated by HCDR3 residues D100c, D100d, and D100g, while R169 from two different protomers were recognized by E32HCDR1 and D56HCDR2 (**Figure 3B, bottom**). For the 42056-a.01 complex, consecutive anionic residues D100aHCDR3 and D100bHCDR3 formed salt bridges with R166 from two protomers while the third R166 residue was engaged by cation-π interactions with F100HCDR3 (**Figure 3C, bottom**). In addition, K168 and K169 from two different protomers were recognized through salt bridges formed with D33LCDR1 and E100gHCDR3, respectively. The 44715-a.01 paratope recognizing Env was entirely comprised of HCDR3, which included F100dHCDR3 and Y100gHCDR3 stabilizing the elongated aliphatic chains of K169 on two protomers and salt bridges formed between cationic residues on all three protomers: D100aHCDR3 interacted with K169, D100iHCDR3 with another K169, and D100eHCDR3 with R166 (**Figure 3D, bottom**).

In addition, like PGT145, 6070-a.01, T646-a.01, and 42056-a.01 each contained tyrosine-sulfated HCDR3 tips that penetrated deep enough into the trimer to form salt bridges with conserved residue K121 (**Figure 3A-C, S4**). 44715-a.01 also had a tyrosine-sulfated HCDR3 tip, but it did not insert as deeply into the trimer; instead, this sulfated tyrosine formed salt bridges with R166 from two different protomers (**Figure 3D, S4**). An overlay of these four structures with the Env complexes of PGT145, PCT64-35S, and RHA1 revealed an alignment of HCDR3 loops extending along the C3 axis into the trimer despite a constellation of unique Fab orientations (**Figure 3E, Figure S5A**). Apart from 44715-a.01, whose HCDR3 penetrated ∼10 Å shallower than the other lineages, the sulfated HCDR3 tips of the other six lineages aligned within the middle of the trimer. In particular, the sulfated-tyrosine residues in five of these structures were positioned at precisely the same location, while this sulfated-tyrosine of the sixth structure (42056-a.01) was just one residue position downstream.

Despite originating from a common D gene and sharing the same overall mode of recognition, the structural role played by D gene-encoded amino acids diverged in these four antibodies. In three of the lineages, the Tyr of the D-gene-encoded EDDYG motif was tyrosine sulfated and inserted into the V2-apex hole; in the 44715-a lineage, the EDDYG motif formed part of the supporting stem scaffold, not the tip, and was substantially affinity matured to *V*D*EGF* (**Figures 2A-D**). Regardless of the different structural role played by the D gene, these data nonetheless demonstrate chemical and structural mimicry of the human PGT145 lineage to be a reproducible mode of broadly neutralizing V2 apex recognition in rhesus macaques.

#### PG9-like recognition

Cryo-EM structures of 41328-a.01 and V033-a.01 in complex with BG505 DS-SOSIP revealed modes of V2 apex recognition similar to human broadly neutralizing antibodies PG9 and CH03, which utilize β-strand pairing of the V2 C-strand as a focus of trimer apex recognition (**Figure 4C**) ^4,7,23,24^. While 41328-a.01 solely bound Env with a 1:1 stoichiometry that yielded a single 3D reconstruction of 2.9 Å resolution, we obtained reconstructions for 1, 2 and 3 V033-a.01 Fab-bound Env complexes (**Figure S5C**). To facilitate comparison with other rhesus and human lineages, we solved the atomic structure of V033-a.01 using the 3D reconstruction of the single Fab-bound complex, which extended to 3.1 Å resolution. 41328-a.01 and V033-a.01 Fabs each exhibited asymmetric recognition of the trimer apex by penetrating between the N156 and N160 glycans of one protomer and binding a second N160 glycan from an adjacent protomer (**Figure 4A, 4B**). These apical glycan interactions were substantial, contributing 52% and 68% of the total interactive surface areas for 41321-a.01 and V033-a.01, respectively (**Table S5**). The structures also revealed both lineage HCDR3s to contain an axe-like subdomain that recognized the C-strand from a single protomer through parallel (41328-a.01) or antiparallel (V033-a.01) β-strand interactions. Two parallel hydrogen bonds formed between the mainchains of 41328-a.01 residue Y100fHCDR3 and Env residues 167 and 168, and four antiparallel hydrogen bonds formed between the mainchains of V033-a.01 residues F100cHCDR3 and G100aHCDR3 and Env residues 169 and 171. For 41328-a.01, a third mainchain hydrogen bond was formed between the backbone amide of Env residue 171 and the side chain of E100iHCDR3.

**Figure 4.**
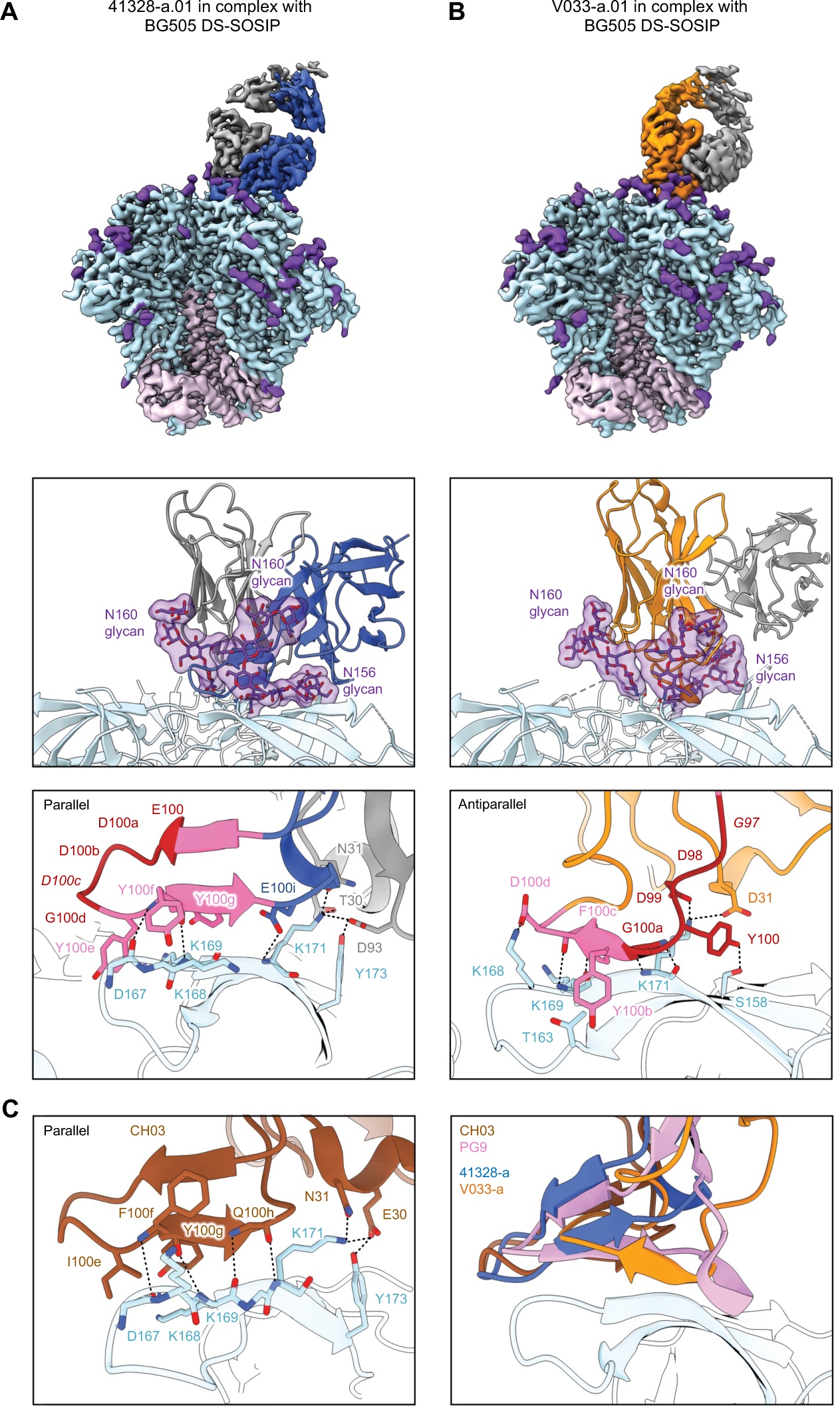
Cryo-EM structures reveal PG9-like modes of V2 apex recognition to be a reproducible antibody extended-class in rhesus macaques. (A) Top; cryo-EM reconstruction of 41328-a.01 in complex with BG505 DS-SOSIP at 2.9 Å resolution. The 41328-a.01 heavy and light chains are colored blue and gray, respectively. Envelope gp120, gp41, and *N*-linked glycans are colored turquoise, pink, and purple, respectively. Middle; expanded interface view of 41328-a.01 from Top panel to highlight binding position and interactions with apical envelope glycans. Glycans bound by 41328-a.01 are shown in stick representation with transparent surfaces. Bottom; further expanded interface view of 41328-a.01 to highlight interactions with apical envelope residues. Interacting residues are depicted in stick representation. Residues at positions corresponding to the conserved five-residue DH3-15*01 gene motif are colored dark red, while the remaining D gene residues are colored pink. Conserved motif position labels are italicized when subjected to somatic hypermutation. Nitrogen atoms are colored blue and oxygen atoms colored bright red. Hydrogen bonds and salt bridges (distance < 3.3 Å) are depicted with dashed lines. The orientations of the C-strand and HCDR3 β-strand mainchain interactions are labeled in the top left corner. (B) Top; cryo-EM reconstruction of V033-a.01 in complex with BG505 DS-SOSIP at 3.1 Å resolution. The V033-a.01 heavy chain is colored orange and the remainder of the complex is colored similarly to panel (A). Middle; expanded interface view of V033-a.01 from Top panel to highlight binding position and apical glycan interactions is shown similarly to panel (A). Bottom; further expanded interface view of V033-a.01 to highlight apical residue interactions is shown similarly to panel (A). (C) Right; Expanded HCDR3 interface view of CH03 (PDB-5ESV) to highlight apical residue interactions is shown similarly to panel (A). Left; expanded HCDR3 interface side view of the alignment of SOSIP complex structures of 41328-a.01 and V033-a.01 determined here to an envelope complex with human Fab PG9 (PDB-8FL1) and V1V2-scaffold complex with human Fab CH03 (PDB-5ESV). Alignments were made with the V1V2 region from each complex. Only gp120 of the 41328-a.01 complex is shown for clarity.

We also observed a number of interactions with Env residue side chains for both structures that closely resembled those of CH03 (**Figure 4C, left**). In the 41328-a.01 complex, a string of three aromatic residues (Y100eHCDR3, Y100fHCDR3, Y100gHCDR3) stabilized the extended aliphatic chains of Env C-strand residues K168 and K169 in an identical manner to the string of three hydrophobic residues (I100eHCDR3, F100fHCDR3, Y100gHCDR3) utilized by CH03. V033-a.01 similarly utilized two consecutive aromatic residues (Y100bHCDR3 and F100cHCDR3) to stabilize the aliphatic chains of K168 and K169, while also engaging K168 through a salt bridge mediated by D100dHCDR3. 41328-a.01 further engaged the C-strand through light chain residues T30LCDR1 and N31LCDR1 forming hydrogen bonds with K171 and D93 LCDR3 forming a salt bridge with K171 and hydrogen bond with Y173; these interactions with K171 and Y173 were strikingly similar to those of CH03 mediated by heavy chain residues E30HCDR1 and N31HCDR1. V033-a.01 could instead engage K171 with two potential salt bridges through D31HCDR1 and D99HCDR3.

An overlay of these two structures with the Env complex of PG9 and the V1V2 scaffold complex of CH03 revealed an approximate alignment of their respective HCDR3 subdomains positioned parallel with the trimer apex plane (**Figure 4C, right**). The four structures demonstrated Fabs to engage Env with one of two different heavy and light chain orientations that were rotated by ∼90° and did not segregate by species (**Figure S5B**). There was exceptional overlap between the HCDR3s of 41328-a.01 and CH03, while the longer HCDR3 of PG9 extended further back along the C-strand before a helical turn redirected and closed the subdomain. Although the HCDR3 length of V033-a.01 was the same as 41328-a.01 and just one residue shorter than CH03, its unique antiparallel subdomain was significantly more compact and did not extend beyond the C-strand toward the trimer C3 axis like the other PG9-like lineages. This smaller footprint provides a structural explanation for the ability of V033-a.01 to also exhibit 1:2 and 1:3 binding stoichiometries with the prefusion-closed conformation of Env (**Figure S5C**).

Differences between 41328-a.01 and V033-a.01 were also apparent in the different structural role played by D gene-encoded amino acids. In 41328-a.01, the D gene-encoded EDDYG did not contact C-strand-interacting residues, while in V033-a.01, Asp and Tyr sidechains did contact the C-strand and the terminal glycine participated in antiparallel mainchain hydrogen bonding. Collectively, these structures demonstrate chemical and structural mimicry of the human PG9 and CH01 lineages, and show β-strand pairing with the C-strand at the trimer apex to be a reproducible mode of broadly neutralizing V2 apex recognition in rhesus macaques.

#### VRC26-like recognition

The cryo-EM structures of V031-a.01, 6561-a.01, and 40591-a.01 (determined with molecular reconstructions at 3.1 Å, 4.1 Å, and 4.2 Å resolution, respectively) revealed modes of V2 apex recognition similar to human broadly neutralizing antibody VRC26.25 ^8,30^. This antibody uses a combined-mode to engage simultaneously both the C-strand and trimer apex hole (**Figure 5G**) ^35^. A single Fab of all three rhesus antibodies bound asymmetrically to the trimer C3 axis with an extended HCDR3 that penetrated between the N160 glycans of two adjacent protomers (**Figure 5A,C,E**). Each lineage also recognized N156 glycan from the right-adjacent protomer (perspective from the Fab body towards Env), but to different extents; whereas 6561-DH1020 buried 354 Å^2^ of N156 glycan surface area, V031-a.01 and 40591-a.01 buried only 34 and 89 Å^2^, respectively. The latter two rhesus lineages were similar to VRC26.25, which buried just 58 Å^2^ of N156 glycan surface area (**Table S5**).

**Figure 5.**
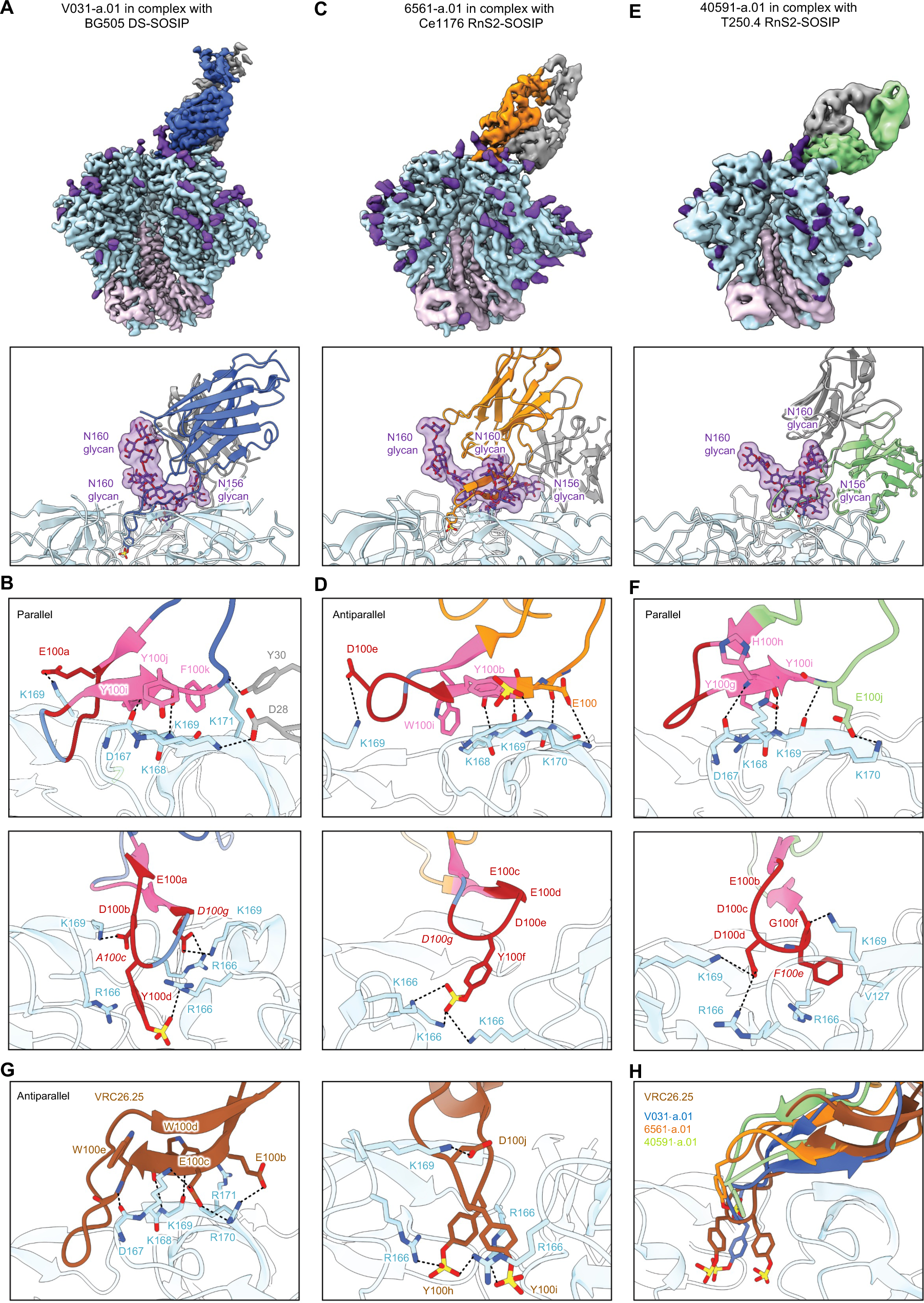
Cryo-EM structures reveal VRC26-like modes of V2 apex recognition to be a reproducible antibody extended-class in rhesus macaques. (A) Top; cryo-EM reconstruction of V031-a.01 in complex with BG505 DS-SOSIP at 3.1 Å resolution. The V031-a.01 heavy and light chains are colored blue and gray, respectively. Env gp120, gp41, and *N*-linked glycans are colored turquoise, pink, and purple, respectively. Bottom; expanded interface view of V031-a.01 from Top panel to highlight binding position and interactions with apical envelope glycans. Glycans bound by V031-a.01 are shown in stick representation with transparent surfaces. Sulfated tyrosine residues are shown in stick representation to highlight their position within the trimer. (B) Further expanded interface views of V031-a.01 to highlight interactions with apical envelope residues. Interacting residues are depicted in stick representation. Residues at positions corresponding to the conserved five-residue DH3-15*01 gene motif are colored dark red, while the remaining D gene residues are colored pink. Conserved motif position labels are italicized when subjected to somatic hypermutation. Nitrogen atoms are colored blue, oxygen atoms colored bright red, and sulfur atoms colored yellow. Hydrogen bonds and salt bridges (distance < 3.3 Å) are depicted with dashed lines. Top; interactions made with primary recognized C-strand. The orientations of the C-strand and HCDR3 β-strand mainchain interactions are labeled in the top left corner. Bottom; interactions mediated by HCDR3 residues inserted into the trimer hole. (C) Top; cryo-EM reconstruction of 6561-a.01 in complex with Ce1176 RnS2 SOSIP at 4.1 Å resolution. The 6561-a.01 heavy chain is colored orange and the remainder of the complex is depicted similarly to panel (A). Bottom; expanded interface view of 6561-a.01 from Top panel to highlight binding position and interactions with apical glycans is shown similarly to panel (A). (D) Further expanded interface views of 6561-a.01 to highlight interactions with apical residues are shown similarly to panel (B). (E) Top; cryo-EM reconstruction of 40591-a.01 in complex with T250.4 RnS2 SOSIP at 4.2 Å resolution. The 40591-a.01 heavy chain is colored orange and the remainder of the complex is color similarly to panel (A). Bottom; expanded interface view of 40591-a.01 from Top panel to highlight binding position and interactions with apical Env glycans is shown similarly to panel (A). (F) Further expanded interface views of 40591-a.01 to highlight interactions with apical Env residues are shown similarly to panel (B). (G) Expanded HCDR3 interface views of VRC26.25 (PDB-6VTT) to highlight apical envelope residue interactions is shown similarly to panel (B). Left; interactions mediated by HCDR3 residues inserted into the trimer hole. Right; interactions mediated by all other Fab residues. (H) Expanded HCDR3 interface side view of the alignment of evelope complex structures of V031-a.01, 6561-a.01, and 40591-a.01 determined here to the envelope complex of VRC26.25 (PDB-6VTT). Alignments were made with gp120 from each complex, while only gp120 of the VRC26.25 complex is shown for clarity. Sulfated tyrosine residues are shown to highlight their positioning within the trimer. Residue F100e of 40591-a.01 is also shown since it is similarly inserted into the trimer hole but cannot be modified posttranslationally.

The HCDR3s from all three lineages recognized the C-strand through a combination of sidechain and mainchain interactions (**Figure 5B,D,F, top**). The V031-a.01 complex revealed the mainchain of Y100jHCDR3 to make two parallel strand hydrogen bonds with the mainchain carbonyl and amide of Env residues 167 and 168, respectively, while a string of aromatic residue sidechains (Y100iHCDR3, Y100jHCDR3, F100kHCDR3) stabilized the extended aliphatic chains of C-strand residues K168, K169 and K171 (**Figure 5B, top**). Further, heavy chain residue E100aHCDR3 was positioned such that it could form a salt bridge with K169 from the C-strand on the right-adjacent protomer in a manner similar to VRC26.25. Additional C-strand interactions with the primary protomer were made by the V031-a.01 light chain: Y30LCDR1 hydrogen-bonded with K171 and D28LCDR1 formed a salt bridge with K168. For 6561-a.01, the mainchains of E100HCDR3, Y100aHCDR3 and Y100bHCDR3 made three antiparallel strand hydrogen bonds with Env residues 169 and 170 (**Figure 5D, top**). Sidechain interactions included the two salt bridges formed between K170 and sulfated Y100bHCDR3 and between K171 and E100HCDR3, while W100iHCDR3 stabilized the extended aliphatic chain of K168. Like V031-a.01, the 6561-a.01 heavy chain residue D100eHCDR3 could establish an additional salt bridge with K169 from the C-strand on the right-adjacent protomer. Lastly, the 40591-a.01 complex demonstrated that the mainchains of Y100gHCDR3, H100hHCDR3 and Y100iHCDR3 to form three parallel strand hydrogen bonds with Env residues 167 and 169 (**Figure 5F, top**). 40591-a.01 further recognized the C-strand through a salt bridge formed between K171 and E100jHCDR3; cation-π interactions formed by K168 sandwiched between two His residues (H30HCDR1 and H100hHCDR3); and stabilization of the K168 aliphatic chain through Y100gHCDR3.

Like VRC26.25, the HCDR3 tips of V031-a.01, 6561-a.01, and 40591-a.01 extended beyond the C-strand and dipped into the middle of the trimer hole nearly along the C3 symmetry axis (**Figure 5B,D,F, bottom)**. However, unlike VRC26.25 which inserted two sulfated Tyr residues, the rhesus lineages utilized either one (V031-a.01 and 6561-a.01) or none (40591-a.01). Notably, 40591-a.01 lacked a Tyr at the tip of its HCDR3 due to a Y100eF somatic mutation, thereby precluding this posttranslational modification (**Figure 2B**). 6561-a.01 made apex hole interactions most similar to VRC26.25: sulfated residue Y100fHCDR3 was positioned to form salt bridges with Env residue 166 from all three protomers (**Figure 5D, bottom**). The V031-a.01 sulfated Y100dHCDR3 residue was positioned such that a salt bridge could be formed with a single R166 while mediating cation-π interactions with R166 from a second protomer (**Figure 5B, bottom**). V031-a.01 also inserted two additional anionic residues (D100bHCDR3 and D100gHCDR3) that formed salt bridges with K169 from two separate protomers; D100gHCDR3 could also interact with the third R166 residue. Despite lacking a sulfated tyrosine, 40591-a.01 still engaged apex hole residues from all three protomers (**Figure 5F, bottom**). F100eHCDR3 interfaced with V127 and could form cation-π interactions with R166, while D100dHCDR3 formed salt bridges with a second R166 residue and K169 from a separate protomer in a manner similar to V031-a.01.

An overlay of these three structures with the Env complex of VRC26.25 revealed an approximate alignment of their respective HCDR3s at the simultaneously engaged C-strand and apex hole epitopes, but not the Fab bodies themselves (**Figure 5H, Figure S5D**). The center of the VRC26.25 Fab was positioned ∼10 Å further from the Env surface due to its exceptionally long HCDR3 that was 10 to 12 residues longer than each of the rhesus lineage HCDR3s (**Figure 1B**). The relative orientation of the 6561-a.01 heavy and light chains overlapped with VRC26.25, while V031-a.01 and 40591-a.01 were rotated ∼45° and ∼180°, respectively (**Figure S5D**). The structure of the V031-a.01 HCDR3 tip was most similar to VRC26.25, whereas the inserted aromatic residues of 6561-a.01 and 40591-a.01 were ∼7 and ∼9 Å shallower than VRC26.25, respectively. These data indicate that the defining VRC26 lineage HCDR3 topology does not require exceptionally residue length; instead, the resulting distal positioning of the VRC26.25 Fab body likely enables the lack of critical interactions with N160 glycan shared by rhesus lineages (**Figure S2B**). Overall, we found chemical and structural mimicry of the human VRC26 lineage to be a reproducible mode of broadly neutralizing V2 apex recognition in rhesus macaques.

### Rhesus DH3-15*01 residues adopt distinct structures to achieve V2 apex-recognizing HCDR3 topologies

To provide a comprehensive view of how DH3-15*01 is utilized to recapitulate the three HCDR3-dominated mechanisms of human V2 apex broadly neutralizing antibodies, we evaluated and compared the conformations and interactions of D gene-derived residues of the rhesus lineages that engage prefusion-closed Env (**Figure 6A-C**).

**Figure 6.**
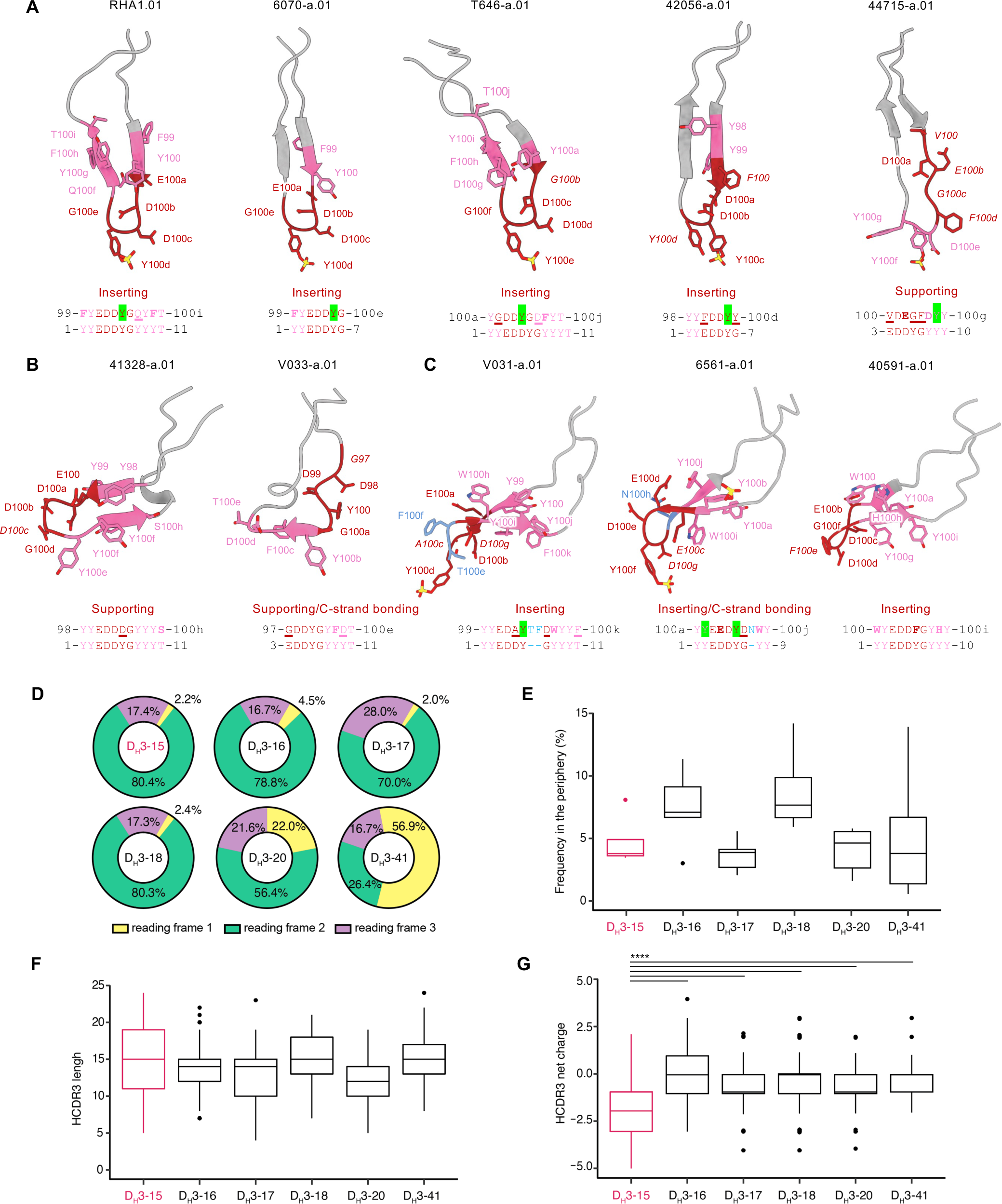
The rhesus DH3-15*01 exhibits structural plasticity and encodes a rhesus-specific anionic motif that is advantageous for V2 apex epitope recognition. (A) HCDR3 structures from rhesus PGT145-like Fabs in complex with envelope trimers. DH3-15*01 conserved five-residue motif (EDDYG) positions are colored and labeled in dark red and the remaining D gene positions are colored and labeled in pink. Conserved motif position labels are italicized when subjected to somatic hypermutation. D gene position side chains are shown in ball and stick representation with nitrogen atoms colored blue, oxygen atoms colored red, and sulfur atoms colored yellow (Kabat numbering). The remaining HCDR3 residues are colored gray with side chains hidden. Below each structure is an alignment of the germline DH3-15*01 coding fragment that was acquired during VDJ recombination (bottom) with the sequence at these positions in the mature antibody (top; Kabat numbering). Somatic mutations that conserve side chain aromaticity or anionic charge are depicted in bold while discordant somatic mutations are underlined. Sites of tyrosine sulfation are highlighted in green. The functional role of the conserved five-residue motif segment is written above the alignment. (B) HCDR3 structures from rhesus PG9-like Fabs in complex with envelope trimers. Structures are depicted similarly to panel (A). (C) HCDR3 structures from rhesus VRC26-like Fabs in complex with SOSIP trimers. Structures are depicted similarly to panel (A). Residue insertions in these structures and sequence alignments are colored blue. (D) Proportion of reading frame usage among naïve rhesus B cells derived from DH3 genes in the peripheral Indian rhesus macaque repertoire. DH3-15*01 is highlighted in red here and in the remaining panels. B cells derived from rhesus DH3-19*01 are exceedingly rare and therefore excluded from analysis. (E) Frequency of DH3 family usage in reading frame two among all naïve B cells in the peripheral rhesus repertoire. (F) HCDR3 length distributions of naïve rhesus B cells in the peripheral rhesus repertoire derived from DH3 genes in reading frame two. (G) HCDR3 net charge distributions of naïve rhesus B cells in the peripheral repertoire derived from DH3 genes in reading frame two. The net charge of DH3-15*01-derived HCDR3s is more anionic (****, p<0.00001 Student’s t-test) than all other groups. Charge calculations only consider amino acid residues and not predicted sites of tyrosine sulfation.

The β-turn and one descending or ascending β-strand of all rhesus PGT145-like HCDR3s contained D gene-derived residue positions, with the minimal conserved five-residue motif encompassing the HCDR3 tip in four of five lineages (**Figure 6A**). HCDR3 residues at these positions engaged conserved Env residues 121, 166 and 169 from one or more protomers, most commonly through electrostatic interactions. When these lineages incorporated any of the first two (1YY2) or last three (9YYT11) D gene residues, they were left unchanged or somatically mutated to residues that conserved sidechain aromaticity (**Figure 2B,6A**). The four rhesus lineages which inserted their HCDR3s as deeply as human antibody PGT145 preserved the germline coded 4DDY6 motif, whereas the fifth lineage (44715-a) retained anionic residues at these positions and acquired a similar 8DY9 motif further upstream with a somatic mutation at position eight. Notably, the germline coded Tyr residue present in either of these motifs was the site of posttranslational sulfation for each lineage (**Figures 3, 5, S4**). The Tyr at position eight was excluded or mutated in all lineages, suggesting this residue to be unfavorable for PGT145-like V2 apex recognition.

The β-strand of the PG9-like HCDR3s that engages the Env C-strand is composed of D gene derived residue positions, while the minimal conserved five-residue motif is positioned in two significantly different conformations (**Figure 6B**). HCDR3 residues at these positions made contacts with Env and glycan residues that are critical for neutralization, but there was not a common pattern of interactions as observed for the lineages with needle-like and combined-mode HCDR3s. Nevertheless, these two lineages each acquired a nine-residue D gene fragment (3EDDYGYYT11) during VDJ recombination that underwent minimal somatic mutation (**Figure 2B, 6B**). As a result, the mature 41328-a.01 and V033-a.01 antibodies each preserved germline coded 4DD5 and 7GYY9 motifs. Both 41328-a.01 and V033-a.01 lacked tyrosine sulfation (**Figure S4**) like human antibody CH03, whose mode of recognition these two rhesus lineages most closely recapitulate.

The inserted HCDR3 tip and one to two β-strands of all combined-mode HCDR3s are composed of D gene-derived residue positions, with the position of the minimal conserved five-residue motif encompassing most or all of the HCDR3 tip for each lineage (**Figure 6C**). HCDR3 residues at these positions comprised a majority of the respective paratopes recognizing both C-strand and trimer hole epitopes though interactions with Env residues 166 and 169 on multiple protomers. These three rhesus lineages had each acquired an eight-residue D gene fragment (1YYEDDYGY8) that underwent minimal somatic mutation, particularly at the N-terminal end (**Figure 2B, 6C**). When any of the first two (1YY2) or last three (8YYY10) D gene Tyr residues were incorporated, they were left unchanged or somatically mutated to residues that conserved sidechain aromaticity. In addition, all three lineages retained a double anionic 3ED4 or 3EE4 motif. For both V031-a.01 and 6561-a.01, the germline coded Tyr at position six was retained and bore the posttranslational sulfation. As mentioned above, this Tyr residue was somatically mutated to Phe (Y100eFHCDR3) in 40591-a*01, the only VRC26-like antibody that did not exhibit tyrosine sulfation (**Figure 5F**, **S4**). Interestingly, antibody 40591-a.05 was part of a distinct phylogenetic clade (40591-a.05-08) that retains the germline D gene position six Tyr (Y100eHCDR3) (**Table S2**) and was sulfated as determined by mass spectroscopy.

Thus, residues coded by the DH3-15*01 gene exhibit structural plasticity that enable their incorporation into three reproducible HCDR3 topologies in otherwise diverse immunogenetic backgrounds to mediate contacts with mode-specific components of the V2 apex epitope.

### The rhesus-specific anionic motif encoded by DH3-15*01 is advantageous for V2 apex epitope recognition

Having elucidated atomic-level interactions between diverse rhesus broadly neutralizing antibodies and the HIV-1 V2 apex, we sought to investigate the immunogenetic basis for the invariant usage of the DH3-15*01 gene by 13 of 13 rhesus antibody lineages (**Figure 2**). We first interrogated the rhesus D gene repertoire ^32^ for homologs of the three different D genes from which HCDR3-dominated human V2 apex broadly neutralizing lineages are derived (**Figure S6A**). The rhesus DH3-41*01 gene is a homolog of the human DH3-3*01 gene utilized by the PG9, VRC26, and PCT64 lineages. Alleles of DH3-41*01 bear one to two amino acid substitutions from the ten-residue long DH3-3*01 gene, with an Asn substitution shared by all alleles resulting in a YY<u>N</u> motif instead of the YY<u>D</u> coded by DH3-3*01; the latter YYD motif is retained in the PG9 and VRC26 lineages and the terminal Asp residue is retained by PCT64. However, three rhesus non-DH3-3*01 homolog D genes code the same YYD sequence, thus enabling rhesus lineages to acquire this three-residue motif through VDJ recombination. The rhesus DH3-18*01 gene is a homolog of the human DH3-10*01 gene utilized by the CH01 lineage and has substitutions at the C-terminal end. However, since the C-terminal end of DH3-10*01 was excluded from the CH01 lineage VDJ recombination event, rhesus lineages can acquire the same D gene sequence as CH01. Lastly, the rhesus DH4-25*01 gene is a homolog of the human DH4-17*01 gene utilized by the PGT145 lineage. This rhesus D gene bears a single substitution and codes a five-residue DYG<u>N</u>Y fragment instead of DYG<u>D</u>Y coded by DH4-17*01. PGT145 retains an anionic residue at this position (E100hHCD3) that forms a salt bridge with Env residue R166, suggesting this to be an important component of the HCDR3 paratope that rhesus lineages would not acquire through VDJ recombination with this D gene (**Figure 3E**). Notably, all substitutions in rhesus D gene homologs are the result of single nucleotide polymorphisms and may be readily mutated to match the human germline sequence during affinity maturation. In summary, a lack of homology with human D genes effectively utilized by human V2 apex broadly neutralizing antibodies is likely not responsible for restricting rhesus lineages to DH3-15*01.

We next interrogated the rhesus D gene repertoire for sequence characteristics shared by V2 apex-targeted rhesus and human lineage HCDR3s: length, aromatic residues, and anionic residues (**Figure 7A**). Similar to the human D gene repertoire, the rhesus DH2 and DH3 families are comprised of genes with some of the longest nucleotide lengths, a majority of which could contribute up to 10 residues in at least one reading frame. The longest D gene fragments in these families were 11 residues, which was expressed by IGHD3-15*01 in reading frame two and IGHD3-20*01 in reading frame three. The DH3 family was also comprised of the genes coding for the greatest number of aromatic residues in the D gene repertoire, which ranged from four to six residues in reading frame two. Finally, we found that DH3-15*01 alone expresses three anionic residues in reading frame two, which is the greatest number in the D gene repertoire. The next highest number was two anionic residues expressed in reading frame two by DH1-7*01, DH3-17*01, DH3-19*02_S4720, and DHD3-20*01. Overall, we found that the rhesus DH3 family contains multiple genes that combine two or more sequence features favorable for V2 apex recognition despite the exclusive use of DH3-15*01 by rhesus lineages. To investigate any functional VDJ recombination biases that could explain this phenomenon, we analyzed the features of DH3-derived HCDR3s in the peripheral rhesus B cell repertoire. DH3 family genes must be expressed in reading frame two to code for their characteristic anionic and aromatic residues and be devoid of stop codons. ∼80% of naïve B cells derived from DH3-15*01 incorporated this gene in reading frame two, which was a similar frequency to those derived from DH3-16*01, DH3-17*01, and DH3-18*01 (**Figure 6D**). Naive B cells derived from DH3-15*01 in reading frame two were not overrepresented in the periphery nor did they have substantially longer HCDR3s than B cells expressing other DH3 family genes (**Figure 6E, 6F**). However, the net HCDR3 charge distribution of DH3-15*01-derived B cells was significantly more anionic (p<0.00001) (**Figure 6G**).

**Figure 7.**
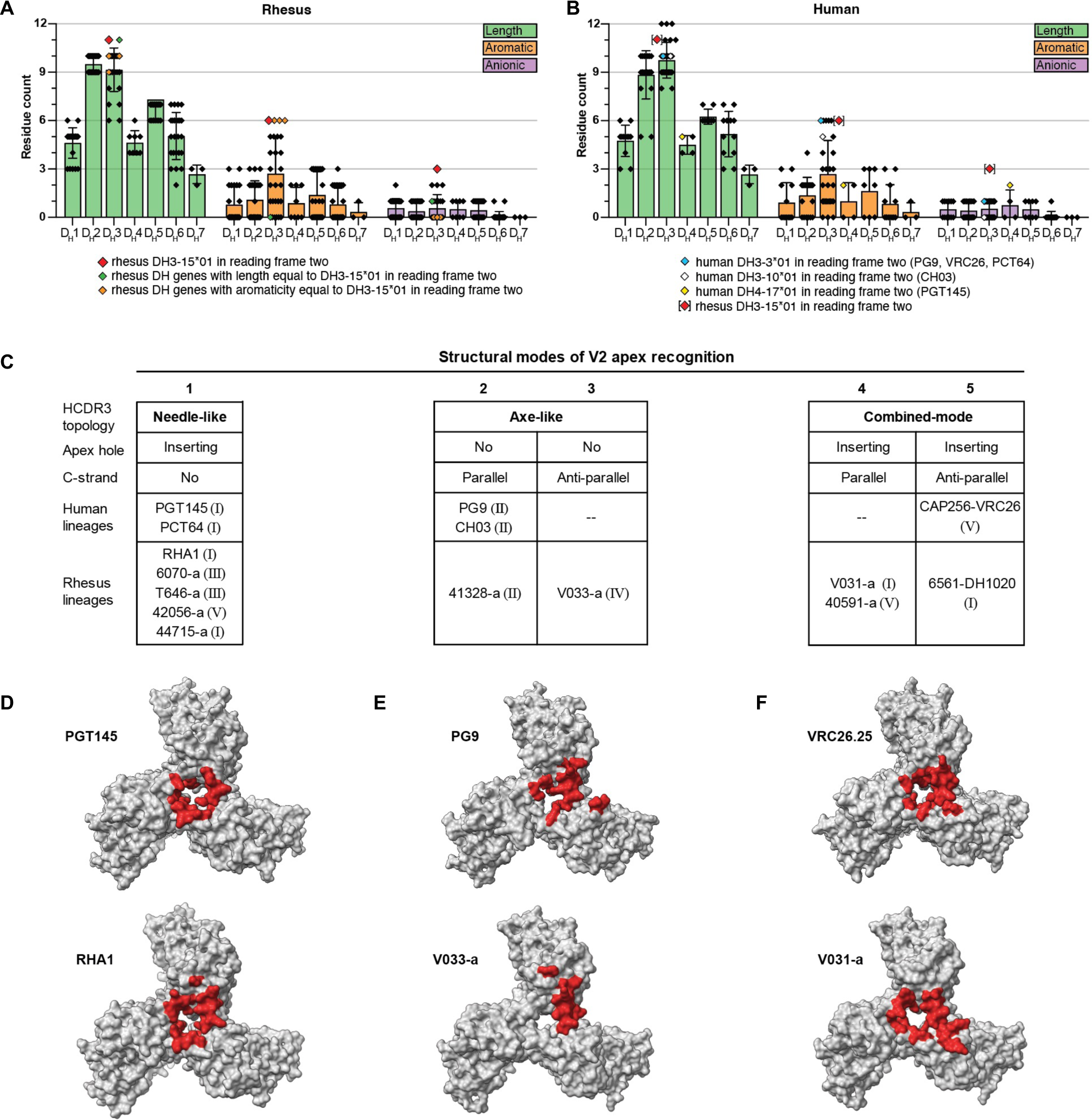
Cross-species recognition of the HIV-1 V2 apex site of vulnerability. (A,B) Features of the Indian rhesus macaque and human DH coding sequence repertoire partitioned by D gene family. Each dot represents a germline D gene expressed in one of the three forward reading frames with a value corresponding to the total number of residues (Length), the number of Tyr, Trp, and Phe residues (Aromatic), and the number of Glu and Asp residues (Anionic). Boxes extend to the respective D gene family mean value and whiskers show the standard deviation. Sequences were excluded if stop codons are found in the middle of the coding sequence in any particular reading frame. In panel A, the rhesus DH3-15*01 gene expressed in reading frame two is marked with a red symbol and rhesus D genes with length and aromatic features equal to to DH3-15*01 in reading frame two are marked with green and orange symbols, respectively. In panel B, D genes utilized by HCDR3-dominated human V2 apex lineages are marked according to the legend and the rhesus DH3-15*01 gene expressed in reading frame two is included as a reference to facilitate comparison. (C) Summary of rhesus and human structural modes of HCDR3-dominated V2 apex recognition. Roman numerals denote the neutralization mapping group (groups I-V) for each lineage as detailed in Figure S2D. (D-F) The 5 Å Env footprints of the most broadly neutralizing rhesus (top) and human (bottom) lineages with PGT145-like (B), PG9-like (C), and VRC26-like (D) modes of recognition are mapped in red onto each respective trimer complex. The remaining gp120 surface is shown in gray and all other components of the structure are omitted for clarity. Top view of trimer.

While single component analysis showed the unique three-residue anionic motif expressed by DH3-15*01 in reading frame two to be the largest statistical outsider for the predilection of rhesus V2 apex-targeted lineages to utilize this D gene, it may be the combination of length, aromaticity, and negative charge, which coalesce to make this D gene so highly over-represented. DH3-15*01 in its second reading frame was an outlier compared to nearly all other D genes in length, aromaticity, and net negative charge (**Figure 7A**). We note in this context that these characteristics are all favorable for tyrosine sulfation ^36–38^, which is utilized in many of the inserting HCDR3 motifs observed in rhesus (**Figures 3, 5, 6, S4**) and human V2 apex-targeted broadly neutralizing antibodies ^4,18,34,35,39,40^.

### Comparison of rhesus and human neutralizing antibody recognition of the HIV-1 V2 apex site of vulnerability

To further define modes of V2 apex recognition shared by rhesus and human broadly neutralizing antibodies, we analyzed features of both their structural architecture and their interactive surfaces. For evaluating structural interactions, we used apex hole and C-strand recognition to categorize human and rhesus antibodies into five recognition modes, which showed general concordance with neutralization mapping groups (**Figures 7C, S2D**). There was significant overlap of the Env surfaces recognized by the most broadly neutralizing lineages across all classes with modest variation between species (**Figure 7D-F**). The PG9-like lineages had the smallest footprint on Env, while PGT145-like and VRC26-like lineages appeared indistinguishable except for the deeper PGT145-epitope surface obstructed within the trimer. The average total interactive surface area (protein and *N*-linked glycan epitopes) of PGT145-like lineages was greater than both PG9-like and VRC26-like, which were nearly identical to each other (**Figure S6B**). *N*-linked glycans comprised the greatest fraction of the total interface surface for PG9-like antibodies (50% or greater for all lineages) and the least for VRC26-like antibodies. Rhesus and human antibodies utilized mode-specific angles of approach to recognize these surfaces: PGT145-like lineages consistently recognized the apex surface with a much steeper approach than PG9-like and VRC26-like antibodies. The similar angles of approach for the latter two recognition modes suggest a restricted spatial freedom to penetrate the apical glycan shield and form main-chain β- strand interactions with the Env C-strand. The heavy chain comprised a majority of the paratope for all lineages, a substantial fraction of which was specifically mediated by HCDR3. Human lineages tended to have more HCDR3-dominated recognition, but this difference was not statistically significant. Overall, this analysis suggested features of V2 apex recognition to be recognition mode-specific, not species-specific.

To assess biochemical features favorable for V2 apex-recognition expressed by rhesus DH3-15*01 versus human D genes, we analyzed the rhesus DH3-15*01 gene relative to translated human D genes. We observed the rhesus DH3-15*01 in reading frame two to be at the extreme for aromaticity and negative charge compared with human D genes (**Figure 7B**). With length, however, at 11 residues, it was shorter than three human D genes of up to 12 residues. Of note, the one human D gene sequence with two anionic residues is DH4-17*01 in reading frame two, which is the same gene and reading frame used by the PGT145 lineage; this gene, however, is only five residues long (DYGDY).

Lastly, we evaluated the relative propensity for tyrosine sulfation of rhesus DH3-15*01 in reading frame two versus that in humans. We inserted the coding segment of each D gene into a 15 amino acid loop, with each Tyr iteratively placed at the loop center and assessed by GPS-TSP 41 for its sulfation propensity. With the default medium cutoff of 1.139, the rhesus DH3-15*01 gene contained four predicted sites of tyrosine sulfation, more than any other D gene (**Figure S6E**). With a more stringent cutoff of 2.276, only one site in DH3-15*01 was predicted for tyrosine sulfation, and indeed this tyrosine was observed to be sulfated in 6 of the 13 characterized rhesus lineages (**Figures 6A-C, S4**). Relative to human D genes, the rhesus DH3-15*01 was not a sulfation outlier, with human DH3-22*01 in reading frame two also showing four predicted sites of sulfation using a 1.139 cutoff, and human DH5-18*01 in reading frame three showing two predicted sites at the higher 2.276 cutoff – twice as many as predicted for rhesus DH3-15*01 (**Figure S6F**). Thus, while structural features of V2 apex recognition appeared to be mode-specific and conserved across species, the unique and favorable biochemical properties of DH3-15*01 in reading frame two likely led to it being highly selected in rhesus macaques irrespective of the mode of antibody recognition.

## Discussion

Highly selected genetic elements are often required for antibody recognition, with antibody “classes” defined by antibodies that utilize the same genetic elements and same recognition mode to bind their target antigens ^29,42^. Certain antibody classes, however, can be found in multiple individuals, indicating that they can arise reproducibly, and we have described such antibody classes as being “multidonor.” Examples of multidonor classes include the potent and broadly neutralizing VRC01-class of antibodies, which are found in individuals with especially broad and potent HIV-1 neutralization ^43–45^. These antibodies derive from a VH1-2*02 heavy chain and a light chain with a LCDR3 of five amino acids. The vaccine implications of this reproducibility are being explored by germline-targeting approaches in human VH1-2 knock-in mouse models ^46–50^ and human clinical trials ^51^. Other multidonor classes of antibodies include those that use the VH1-69 gene, such as the stem-directed broadly neutralizing antibodies against influenza A virus ^52–54^ or the CD4-induced class of antibodies that recognize the co-receptor binding site on HIV-1 ^55^. The CD4-induced antibodies not only use VH1-69, but – like the V2 apex antibodies explored here – are often tyrosine sulfated ^55–57^. In prior cases of highly selected genetic elements, epitope recognition is mediated by similar structural motifs, in the case of VRC01 this being the CD4 mimetic HCDR2 ^44,58^. This contrasts with the results of the current study in which the highly selected rhesus DH3-15*01 gene, a clear signature for V2 apex-directed broadly neutralizing antibodies, encodes amino acids that play diverse structural roles: sometimes as an inserting loop, sometimes as a contacting β-strand, and sometimes as part of the supporting non-contacting HCDR3 loop. In these diverse structural roles, multiple modes of V2 apex recognition are used where each mode forms an extended-class, which we previously defined as antibodies that do not necessarily share genetic commonalities but nonetheless display a characteristic mode of antigen interaction ^7^. Remarkably, DH3-15*01 plays diverse structural roles not only between modes, but also within the same extended-class.

This observed structural diversity of the rhesus DH3-15*01 gene – along with its length, aromaticity, and charge – provide a likely mechanistic explanation for its invariant ‘signature’ usage in rhesus V2 apex broadly neutralizing lineages and the high prevalence of V2 apex-directed responses in SHIV-infected rhesus macaques ^17,59^. The ability of highly-selected genetic elements to assume divergent functional roles for antigen recognition expands our understanding of antibody diversity and mulitdonor reproducibility (**Figure S7**).

The striking frequency of V2 apex broadly neutralizing antibodies using the DH3-15*01 gene, with its favorable germline-encoded EDDYG motif, in SHIV-infected rhesus macaques has important implications for the preclinical testing of V2 apex-targeted vaccines: it suggests that the elicitation of V2 apex broadly neutralizing antibodies may be easier to achieve in non-human primates than in humans, which do not have a homologous EDDYG-encoding D gene. That said, this study has shown in a systematic way that the structural and immunogenetic features of rhesus and human V2 apex-targeted neutralizing antibodies exhibit striking similaries and that the rhesus macaque is a relevant model system for HIV-1 vaccine development. The repeated elicitation of V2 apex broadly neutralizing antibodies in rhesus macaques by SHIV infection provides a model system to elucidate essential elements of broadly neutralizing antibody induction by candidate vaccines, including the critical priming, boosting, affinity maturation and “polishing” steps. It will be fascinating to see how much the elicitation probability of V2 apex broadly neutralizing antibodies can be improved by combining SHIV infection with rational vaccine approaches, such as (i) the use of “special” envelope trimers with affinity for germline V2 apex antibodies ^6,7,60,61^; (ii) the use of V2 epitope signatures as immunogens ^62^; (iii) or the use of engineered immunogens with enhanced recognition of germline V2 apex antibodies ^12,63–67^. It will be equally important to decipher key elements in Env-Ab coevolution in SHIV-infected macaques that are responsible for guiding affinity maturation to engender neutralization breadth and potency, and how these natural events can be augmented by rational vaccine design ^68–72^. Such analyses may ultimately enable V2 apex-directed broadly neutralizing antibodies to be elicited by vaccination alone.

## Acknowledgements

We thank the NIH Vaccine Production Program Laboratory (VPPL) for providing the BG505 DS-SOSIP used for cryo-EM studies. We thank the flow cytometry cores at the NIH Vaccine Research Center (VRC) and University of Pennsylvania Perelman School of Medicine for their assistance with antigen-specific single-cell sorting. We thank members of the Columbia University Cryo-EM Center for assistance with the collection of six cryo-EM datasets used in this study. We thank members of the Simons Electron Microscopy Center (SEMC) at the New York Structural Biology Center (NYSBC) for assistance with the collection of three cryo-EM datasets used in this study.

This work was supported by grants from the Bill & Melinda Gates Foundation (INV-007939, INV-041767), the NIH (R61 AI 161818 and AI 167716 [R.A., G.M.S.], R01 AI160607, R01 AI165080, R61 AI161818, UM1 AI144371, R37 AI150590, P30 AI045008 [G.M.S.], R01 AI 050529 and R37 AI 150590 [B.H.H.], R61 AI 176583 [Z.S.]), by the Vaccine Research Center, an intramural Division of the National Institutute of Allergy and Infectious Diseases (NIAID), NIH, and by the Consortia for HIV Vaccine Development (CHAVD) (UM1 AI 144371 [B.F.H]). R.H was supported by a Training Grant in HIV Pathogenesis (T32-AI007632). M.B. was supported by the Division of Intramural Research, NIAID, NIH; part of this work was performed when M.B. was employed by the Duke Human Vaccine Institute in the laboratory of B.F.H.

## Author Contributions

R.S.R., R.H., H.L., M.B., M.H., and R.D.M performed antibody isolation and characterization. R.S.R. and J.G. performed cryo-EM structure determination and analysis, while G.C. provided initial cryo-EM training. R.S.R., R.H., A.J.C., F. B-R., and J.W.C. performed NGS B cell sequencing and analysis. Y.G. performed rhesus naïve B cell receptor repertoire analysis. J.W. and Q.P.L. performed mass spectrometry studies for tyrosine sulfation analysis. H.L., K.J.S., E.L., J.M.R., and A.S. performed rhesus macaque sample processing and plasma neutralization studies. Y.P. performed antibody expression and purification. M.K.L., N.A.D-R., and M.S.S. provided large virus panel neutralization data. R.H., A.S.O., B.Z., D.R.H., T.L., K.O.S., S.C., and W.H. contributed reagents for antibody isolation and structural studies. R.A., K.O.S., B.F.H., D.W.K., M.R., Z.S., B.H.H., G.M.S., P.D.K, and L.S provided experimental oversight and funding. R.S.R., G.M.S., P.D.K., and L.S. wrote the manuscript, while other authors provided comments and edits.

## EXPERIMENTAL MODEL DETAILS

### Nonhuman Primates

All Indian Rhesus macaques used in this study were housed at Bioqual, Inc., Rockville, MD, according to guidelines of the Association for Assessment and Accreditation of Laboratory Animal Care. Experiments were approved by the University of Pennsylvania (IACUC protocol 806719) and Bioqual (IACUC protocol 21-139) Institutional Animal Care and Use Committees. Macaques were sedated for blood draws, anti-CD8 mAb infusions, and SHIV inoculations and cared for according to AAALAC guidelines and best practice standards. The 10 rhesus macaques described in this study were as follows: 5695 (M, 6 yo), 6561 (M, 7 yo), 6070 (M, 4 yo), 40591 (M, 6 yo), 42056 (M, 5 yo), T646 (F, 5 yo), V031 (F, 4 yo), V033 (F, 3 yo), 41328 (M, 6 yo), and 44715 (F, 4 yo). Rhesus macaques 42056, V033, V031, 44715, and 6070 were transiently depleted of CD8 T cells prior to SHIV inoculation with a 25-50mg/kg subcutaneous or intravenous infusion of anti-CD8α (MT807R1) or anti-CD8β (CD8beta255R1) monoclonal antibody (NIH Nonhuman Primate Reagent Resource [NHPRR], https://www.nhpreagents.org). Rhesus macaques were inoculated intravenously with 293T transfection supernatants containing 50 or 500 ng p27 Ag of molecularly cloned SHIV challenge stock, except for RM 5695, which was inoculated with acute phase SHIV-infected rhesus plasma as previously described^17^. SHIVs used to infect each animal are listed in Figure S1 and Table S2. Design features of SHIVs and methods for preparing 293T-derived virus challenge stocks were previously described^75^. Infection of rhesus macaques 5695, 6070, 40591, and 42056 was previously reported in Roark et al^17^. Infection of rhesus macaque T646 was previously reported in and Bauer et al^76^. Rhesus macaques 5695, V033 and V031 were repurposed from previous HIV-1 envelope DNA or SOSIP immunization studies (animal IDs 5695, 177715 and 178951, respectively)^77,78^. All repurposed animals lacked detectable tier-2 neutralizing responses in plasma prior to infection. Rhesus macaque 44715 was infected with SHIV CH848.wk36con, a consensus sequence of plasma viruses at 36 weeks post infection from a SHIV CH848-infected rhesus macaque. SHIV plasmids used in these experiments have been deposited in the American Type Culture Collection (ATCC) under contract with the National Institute of Allergy and Infectious Diseases (NIAID/NIH).

## METHOD DETAILS

### Blood processing

Blood samples were collected in sterile vacutainers containing acid citrate dextrose formula A (ACD-A) anticoagulant. ACD-A blood (40 ml) was centrifuged (1000*g* for 10 min at 20°C) in sterile 50ml conical tubes, and the plasma collected without disturbing the buffy coat white blood cell (WBC) layer and red cell pellet. Plasma was centrifuged again (1500*g* for 15 min at 20°C) to remove platelets, aliquoted into 1-ml cryovials, and stored at −80°C. The cell pellet was resuspended in an equal volume of Hanks balanced salt solution (HBSS) (without Ca/Mg) containing 2 mM EDTA and divided into four 50 ml conical tubes. Additional HBSS-EDTA buffer was added to each tube to bring the total volume to 30 mL. The cell suspension was underlayered with 14 ml of 96% Ficoll-Paque and centrifuged (725*g* for 20 min at 20°C) with slow acceleration and braking. Mononuclear cells at the Ficoll interface were collected and transferred to a new 50-ml centrifuge tube containing HBSS-EDTA and centrifuged (200*g* for 15 min at 20°C). The supernatant was removed, and the cell pellet was resuspended in 40ml HBSS (with Ca/Mg) + 1% FBS. The suspension was centrifuged again (200g for 15 min at 20°C) and the supernatant discarded. Centrifugation at 200g pellets WBCs but allows most platelets to remain in suspension. The mononuclear cell pellet was tap-resuspended in the residual media and then HBSS (with Ca/Mg) + 1% FBS was added to a volume of 10 ml. Cells were counted and viability assessed by trypan blue staining. Cells were centrifuged again (300g for 10 min at 20°C), the supernatant discarded, and the cells resuspended at a concentration of 5-10 × 10^6^ cells/ml in CryoStor cryopreservation media (Sigma cat. no. C2999) and aliquoted into 1ml cryovials (CryoClear cryovials; Globe Scientific Inc., cat. no. 3010). Cells were stored in a Corning CoolCell LX cell freezing container at −80°C overnight and then transferred to vapor phase liquid nitrogen for long-term storage.

### Site-directed mutagenesis

Site-directed mutants in this study were generated using a Q5 site directed mutagenesis kit according to the manufacturer’s instructions (NEB). Briefly, mutagenesis primers were designed using the NEBaseChanger (https://nebasechanger.neb.com/) to contain a mismatch at the residue of interest. The template virus was then PCR amplified with the mutagenesis primers, and the resulting PCR product treated with KLD enzyme mix to remove template DNA and ligate blunt PCR ends. Plasmids were then transformed into *E. coli* DH5α cells (for envelope clones) or MAX Efficiency Stbl2 cells (for full-length SHIV clones) and grown overnight at 30°C. Plasmid preparations were sequence verified to confirm the presence of the desired mutation.

### Virus stocks

SHIV and pseudovirus stocks were prepared by transfection of HEK 293T/17 cells (ATCC, CRL-11268). 4-5 × 10^6^ cells were plated in 100mm dishes in DMEM containing 10% FBS and 1% Penicillin/Streptomycin and incubated overnight at 37°C. 6 µg of plasmid DNA was then transfected using FuGENE 6 transfection reagent (Promega) according to the manufacturer’s instructions. Transfected cells were then incubated for 48 hours at 37°C. Supernatants were centrifuged (2000g for 8 min at 4°C) to remove cell debris, aliquoted, and stored at -80°C. Viruses were titered on TZM-bl cells. Seven 5-fold serial dilutions of virus stocks were made in DMEM with 6% FBS and 40µg/ml DEAE-dextran, added in quadruplicate to adherent TZM-bl cells, and incubated at 37°C for 48h. Following incubation, cells were fixed for 10 minutes at room temperature in PBS containing 0.8% glutaraldehyde and 2.2% formaldehyde. Cells were then washed 3 times with PBS, and stained with PBS containing 4µM Magnesium Chloride, 4 µM Potassium Ferricyanide, 4 µM Potassium Ferrocyanide, and 400 µg/ml X-Gal for 3 hours at 37°C. Stained cells were then washed three times with PBS and imaged on a CTL Immunospot analyzer.

### Neutralization assays

Neutralization assays were performed on TZM-bl indicator cells as previously described^17^. Briefly, TZM-bl cells were plated in 96 well plates in DMEM containing 10% FBS and 1% Penicillin/Streptomycin and grown overnight at 37°C. Serial plasma or monoclonal antibody dilutions were incubated with virus at 37°C for 1 hour. Plasma was diluted in cell culture media containing 5% normal human or rhesus heat-inactivated serum so as to hold the concentration of test plasma/serum constant across all wells. Following incubation, the virus/antibody mixture was added to TZM-bl cells, and the cells incubated for 48h at 37°C. Cells were then lysed with PBS containing 0.5% Triton X-100 at room temperature for 1 hour, and luciferase levels were quantified using the Promega Luciferase Assay (Promega) on a BioTek Synergy Neo2 microplate reader.

### Rhesus monoclonal antibody isolation

Broadly neutralizing antibody lineages were isolated from memory B cells baited with antigen-specific probes through single-cell sorting into PCR plates. PBMCs were thawed, washed in 10 mL RPMI+10% FBS + 2 µl RNase-free DNase I (NEB), and stained with LIVE/DEAD Aqua for 15 minutes at room temperature. PBMCs were then washed twice with PBS and stained with a cocktail of CD3-PerCP-Cy55, CD4-BV785, CD8-BV711, CD14-PE-Cy7, CD20-BV605, IgD-FITC, IgG-AF680, and IgM-BV650 for 15 minutes at room temperature. PBMCs were again washed twice with PBS and then stained with fluorophore-conjugated SOSIP probes (detailed in Table S2) for 15 minutes at room temperature. Memory B cells (CD3-, CD4-, CD8-, CD14-, CD20+, IgG+, IgD-, IgM-) positive for wild type heterologous SOSIP probes and negative for V2 apex epitope mutant probes were sorted at 1 cell per well into 96-well plates on a BD FACSAria II machine using BD FACSDiva software (BD Biosciences). RNA extraction, cDNA synthesis, and VDJ gene amplification were performed as previously described^19^. Wells with successful DNA PCR amplification were Sanger sequenced (Azenta Biosciences) and initially analyzed with IgBLAST^79^ or IMGT V-QUEST^80^ to identify expanded lineages and inspect HCDR3 sequences.

### Bone marrow negative selection

For isolation of antibody 5696-b.01, RM5695 necropsy (week 65) bone marrow aspirate mononuclear cells were thawed, washed in 10 mL RPMI+10% FBS + 2 µl RNase-free DNase I (NEB), and placed in single-cell suspension in 1 mL RPMI+10% FBS. First, 25 µl of Human TrueStain TcX (BioLegend, cat # 422301) was added to the cell suspension and incubated for 10 min at room temperature. To enrich for plasma cells, the total mononuclear cells were then incubated with anti-human CD3-PerCP-Cy55, CD4-BV785, CD8-BV711, CD14-PE-Cy7, CD20-BV605, CD16, and CD36 for 20 min at 4°C under gentle agitation. Following two wash steps in RPMI + 10% FBS, cells were resuspended in 100 µL of autoMACS Running Buffer (Miltenyi Biotec) and 25 µL of anti-mouse IgG microbeads (Miltenyi Biotec) were added, followed by a 15 min incubation at 4°C. After two wash steps with autoMACS Running Buffer, cells were resuspended in 500µl of autoMACS Running Buffer and loaded on an LS Column (Miltenyi Biotec) for magnetic separation using a QuadroMACS Separator (Miltenyi Biotec), following the manufacturer’s protocol. Flowthrough cells were pelleted and resuspended in 250µl of RPMI +10% FBS. Cell viability was analyzed using AO/Pi staining using a Cellometer Auto 2000 Cell Viability Counter (Nexcelom Bioscience) and determined to be 83.7%.

### Single-cell B cell receptor sequencing

Negatively selected bone marrow cell suspension was loaded on a Chromium X instrument (10X Genomics) to generate single-cell beads emulsion, at a loading concentration for a targeted recovery of 10,000 cells per reaction. Single-cell RNA-seq libraries were then prepared using Chromium Next GEM Single Cell 5’ Kit v2 bead and library construction kit (10X Genomics). B cell receptor (BCR) libraries were constructed using Chromium Single Cell Human BCR Amplification Kit (10X Genomics) and Rhesus macaque-specific primers targeting the constant regions of heavy chain IgM, IgG and IgA gene isotypes and light chain IgK and IgL genes, as described previously^78^. BCR libraries were indexed using Dual Index Kit TT Set A kit (10X Genomics) and sequenced on an Illumina NextSeq 2000 instrument at a minimum read depth of 5000 reads/cell. Illumina BCLconvert 3.10.12 was used for demultiplexing and fastq files were analyzed with Cell Ranger 7.2.0’s VDJ pipeline (10X Genomics) using a custom rhesus macaque VDJ germline reference. The 5695-b.01 mAb was identified based on manual analysis of HCDR3 length, sequence homology, and heavv/light chain germline gene usage.

### Monoclonal antibody cloning and synthesis

Antibodies with features characteristic of V2 apex broadly neutralizing antibodies (expanded lineages with long CDRH3s enriched for negative and aromatic residues) were selected for further analysis. Antibody heavy chain VDJ and light chain VJ gene cassettes were synthesized (Genscript) and cloned into rhesus IgG1 (RhCMV-H), IgK (RhCMV-K), or IgL (RhCMV-L) expression vectors upstream of their constant regions using AgeI/NheI, AgeI/BsiWI, and AgeI/ScaI restriction sites, respectively^19^. Plasmids encoding antibody heavy and light chain genes were used to transfect suspension Expi293F cells in a 1:2 heavy to light chain ratio using ExpiFectamine 293 transfection reagents (Gibco) according to the manufacturer’s instructions. Antibodies were purified using a rProtein A/Protein G Gravitrap kit (Cytiva), buffer-exchanged into PBS, and stored at 4°C.

### B cell next generation sequencing

PBMCs were stained with LIVE/DEAD Aqua, CD3-PerCP-Cy55, CD4-BV785, CD8-BV711, CD14-PE-Cy7, CD20-BV605, IgD-FITC, IgG-AF680, and IgM-BV650. Naïve B cells (CD3-, CD4-, CD8-, CD14-, CD20+, IgG−, IgD+, IgM+) and memory B cells (CD3-, CD4-, CD8-, CD14-, CD20+, IgG+, IgD-, IgM-) were bulk sorted into RPMI with 10% FBS and 1% Pen-Strep using a BD FACSAria II sorter. Total RNA was extracted using RNAzol RT per the manufacturer’s guidelines (Molecular Research Center, Inc). Reverse transcription of mRNA transcripts, IgM and IgL variable region library preparation, and next-generation sequencing were performed as previously described^17,81^. Briefly, cDNA was synthesized using a 5’RACE approach, with SMARTer cDNA template switching and Superscript II RT, following which cDNA was purified using AMPure XP beads (Beckman Coulter). cDNA was then PCR amplified using KAPA HotStart ReadyMix (Roche) with IgG, IgM, IgK, or IgL constant region-specific primers to amplify B cell heavy chain VDJ regions and light chain VJ regions. Finally, the libraries were PCR amplified with primers that added on barcodes and Illumina P5 and P7 sequencing adaptors. Both heavy and lambda immunoglobulin libraries were sequenced on an Illumina Miseq sequencer with 2 × 300 bp runs using the MiSeq Reagent V3 kit (600-cycle). Demultiplexed reads were adapter trimmed using Cutadapt v3.5, and paired forward/reverse reads were merged using PEAR v0.9.6 with a quality cutoff of 15 and a minimum length of 50.

### VDJ repertoire analysis

Filtered, quality-controlled IgM and IgL sequences were analyzed using IgDiscover v0.15.1^22^ to curate a personalized immunoglobulin repertoire library for each rhesus macaque (Table S3). Merged reads were used as input, with the KIMDB 1.1 database^32^ (http://kimdb.gkhlab.se/) used as the heavy chain reference and the Ramesh et al.^31^ allele database used as the light chain reference. The IgDiscover J output for both heavy and light chains was additionally filtered with the “discoverjd” feature (using a J coverage setting of 100 and allele ratio of 0.33) to exclude low-confidence novel alleles. This allowed the curation of a repertoire of both known and novel V and J gene alleles for each rhesus macaque, confirmed the presence of known D gene alleles, and allowed the accurate assignment of V, D, and J alleles for each broadly neutralizing antibody lineage. Data from IgG sequencing datasets was analyzed using SONAR v4.2^82^. The manually guided identity/divergence feature was used to identify ancestral clonal sequences. As IgDiscover does not identify novel D gene alleles, per-animal D gene repertoires were further investigated with MINING-D^83^. Deduplicated HCDR3 sequences of SONAR-clustered reads were used as input, allowing the detection of more complete D segments. These were then compared with published references^22,31,32^ to further verify the presence of known D gene alleles in each animal.

### Prefusion-stabilized envelope trimer production

All recombinant HIV-1 and SIVcpz envelope SOSIP trimers used for antigen-specific single-cell sorting and cryo-EM structure analysis were designed, expressed, and purified using previously described methods^30,60,65,84–88^. Plasmids encoding the SOSIP trimer construct and human Furin were mixed at a 4:1 ratio and complexed with 293fectin (Thermo) to transfect 293F cells with ∼0.6 mg DNA/1 L of suspension cell culture. Six days post-transfection, cultures were clarified by centrifugation and native, prefusion-closed trimers were purified from culture supernatant using affinity chromatography with immobilized PGT145. Gel filtration on a Superdex 200 16/600 column (GE Healthcare) was used to further purify and buffer exchange each trimer into PBS. Trimers to be used as probes for single-cell sorting were appended with a C-terminal Avi-tag (GGLNDIFEAQKIEWHED) to enable biotinylation using BirA biotin-protein ligase (Avidity) once purified. Streptavidin-linked fluorophores (SA-PE, SA-BV421, or SA-APC) were added to biotinylated trimers for 75 minutes at 4°C to prepare them as multimerized memory B cell probes.

### Antibody Fab production

Fabs were expressed and purified with one of two approaches. One approach utilized was to insert the HRV3C protease cleavage site (LEVLFQGP) into the hinge region of the plasmid encoding the full-length rhesus IgG1 heavy chain. Antibodies bearing the HRV3C cleave site were expressed and purified as described above by swapping in the modified heavy chain plasmid to be paired with the plasmid encoding the natural light chain. 1 mg of antibody was digested with HRV3C protease (Thermo) for ∼16 hours at 4C, from which liberated Fab fragments were purified using negative selection on a Protein A column. The second approach utilized was to insert an 8x Histidine tag and early stop codon into the hinge region of the plasmid encoding the full-length rhesus IgG1 heavy chain. When co-transfected with the plasmid encoding the natural light chain mixed at a 1:1 ratio, Fab was purified from clarified culture supernatant six days post-transfection using Ni-NTA affinity chromatography with IMAC Sepharose 6 Fast Flow resin (GE Healthcare). Gel filtration on a Superdex 75 10/300 column (GE Healthcare) was used to further purify and buffer exchange each Fab into PBS.

### Cryo-EM sample preparation

Fabs and HIV-1 envelope SOSIP trimer complexes were prepared for cryo-EM data collection as previously described^17,18,35^. Briefly, envelope trimers were concentrated to ∼4 mg/mL and mixed with Fab at a 1:3 molar ratio for a final trimer concentration of ∼2-3 mg/mL. The mixture was incubated on ice for 20 minutes to allow complexes to form before the addition of the detergent n-Dodecyl β-D-maltoside (DDM) at a final concentration of 0.005% (w/v). Copper C-flat Holey carbon-coated grids (CF-1.2/1.3 300 mesh; EMS) were glow discharged using a PELCO easiGlow before the addition of 3 uL of Fab-trimer complexes. Samples were vitrified in liquid ethane using a Vitrobot Mark IV with a wait time of 30 seconds and a blot time of 3 seconds at room temperature with 100% humidity.

### Cryo-EM data collection and processing

Cryo-EM data were collected on a FEI Titan Krios electron microscope operating at 300 kV, equipped with a Gatan K3 direct detector operating in counting mode. Data were acquired using Leginon^89^ at 105,000x magnification with a 0.83 Å pixel size and defocus range of 0.8-1.8 µm. A total exposure dose of 58 e^-^/Å^2^ was fractionated over 50 raw frames. All data processing – including motion correction, CTF estimation, particle extraction, 2D classification, *ab initio* model generation, and 3D refinements – were performed using cryoSPARC^90^. All homogenous and non-uniform 3D refinements were performed with C1 symmetry. With the exception of V033-a.01, all other rhesus lineages exhibited 1:1 stoichiometry of trimer-Fab binding. V033-a.01 was observed to bind with 1:1, 1:2, and 1:3 stoichiometries in 2D classes, and we obtained 3D reconstructions ranging from 3.1-3.6 Å resolution for each. The varied Fab binding stoichiometries are likely a result of the limiting 1:3 molar ratio of trimer to Fab used to prepare complexes for data collection, and not inherent to the biology of V033-a.01.

### Atomic model building

The initial models for all Fab variable regions (Fv) were obtained using the AbodyBuilder2 application of the SAbPred Antibody Prediction Toolbox^91^. The initial models for all HIV-1 envelope trimers were obtained from the Protein Data Bank and are specified in Table S6. For each sample, the respective Fv and envelope trimer models were fit into the cryo-EM 3D reconstruction density using UCSF Chimera^92^ or ChimeraX^93^ to provide an initial model of the Fab-trimer complex. Each structure was solved using iterative rounds of manual model rebuilding in Coot^94^ and automated real-space refinement of the model in Phenix^95^. Overall structure quality was periodically determined using MolProbity^96^ and EMRinger^97^ until satisfactory validation of the model was achieved. Fab interactive surfaces were analyzed using PDBePISA^98^. Summaries of cryo-EM data collection, 3D reconstruction, and model refinement statistics for each structure are provided in Table S6 and Dataset S1.

### Antibody angles of approach and binding orientation

The latitudinal angles of antibody approach to the V2 apex of the HIV-1 envelope trimer were calculated with UCSF ChimeraX^93^. The latitudinal access of a V2 apex-targeted antibody is the freedom between the 3-fold trimer axis and the V2 apex plane, the latter of which we define as the horizonal plane passing through the Cα atoms of C-strand Lys168 residues on all three protomers. Since the rhesus and human antibodies analyzed in this study all utilize HCDR3-dominated modes of recognition, and these HCDR3s all adopt pronounced conformations extending from the Fab combining surface, we next defined the axis of antibody as the long axis of the HCDR3. The HCDR3 axis is the vector connecting the centers (centroids) of the Fv itself and the HCDR3, which are defined as the averaged coordinates of Ca-atoms of the two pairs of Fv conserved cystines and the averaged coordinates of heavy chain residues 93 and 102 (Kabat numbering), respectively. The angle of intersection between the V2 apex plane and the antibody axis were calculated with the built-in function of ChimeraX. To compare the relative orientations of rhesus and human antibody heavy and light chains, alignments were made with gp120 from each trimer complex (residues 128-192 for the CH03-V1V2 scaffold complex [PDB-5ESV]) utilizing the same mode of V2 apex recognition using MatchMaker in ChimeraX.

### Analysis of Indian rhesus macaque peripheral DH3 gene-derived naïve B cells

The B cell repertoires for Indian rhesus macaque DH3 family gene frequency and charge were download from NCBI under accession numbers ERR4250665 to ERR4250672. Germline V, D, and J genes were obtained from the KIMDB 1.1 database^32^. We processed the B cell receptor transcripts using the SONAR version 2.0 bioinformatics pipeline^82^, which includes steps for quality control and annotation. V(D)J gene assignment for each transcript was performed using BLASTn^79^ (https://www.ncbi.nlm.nih.gov/igblast/index.cgi), with customized parameters against the germline gene database from KIMDB. CDR3 identification was based on BLASTn alignments of the V and J regions, utilizing the conserved second cysteine in the V region and the WGXG (heavy chain) or FGXG (light chain) motifs in the J region, where ’X’ represents any amino acid. For heavy chain transcripts, isotype was assigned using the constant domain 1 (CH1) sequences through BLASTn against a database of rhesus CH1 genes from IMGT, with a BLAST E-value threshold of 10^(-6) to determine significant isotype assignments. The CH1 allele with the lowest E-value was selected. Non-V(D)J sequences were removed, and transcripts with incomplete or frameshifted V(D)J regions were excluded. Remaining transcripts were aligned to their assigned germline V and D genes using Clustal Omega^99^. Frequencies, length, and net charges of DH3-derived transcripts were then calculated using custom Python scripts.

### Tyrosine sulfation prediction for rhesus and human DH genes

Human DH genes were downloaded from the IMGT database^80^ (https://www.imgt.org/). Both human and rhesus macaque DH genes were translated into amino acid sequences using reading frames 1, 2, and 3. Subsequently, the GPS-TSP version 1.0 tool^41^ (https://tsp.biocuckoo.org/) was utilized to predict tyrosine sulfation sites within these sequences. The algorithm specifically evaluates each tyrosine by centering it within a 15 amino acid segment, designed to mimic a HCDR3 region of equivalent length. The threshold scores were set at 1.139 for predicted tyrosine sulfation sites and 2.278 for sites with a high likelihood of sulfation.

### Mass spectrometry to quantify tyrosine sulfation

Each full-length IgG1 antibody was diluted to approximately 3 mg/mL using 50 mM ammonium bicarbonate (Thermo). For IdeS digestion and deglycosylation: 8 uL of the antibody solution was placed in a microcentrifuge tube. 1 uL of IdeS (40 units/uL, Promega) was added to the antibody solution and incubated at 37°C for 1 hour. 2 uL of Rapid PNGase F Buffer (NEB) was added and incubated at 80°C for 3 minutes. The solution was cooled, and 1 uL of Rapid PNGase F was added and incubated at 50°C for 30 minutes. After deglycosylation, 40 uL of 50 mM ammonium bicarbonate was added prior to LC-MS analysis. For LC-MS analysis, 3 – 10 uL of the IdeS digested and deglycosylated sample was injected onto a Waters H-class UPLC (Milford, MA), separated with a C4 column (Acquity UPLC Protein BEH C4 column, 300Å, 17 um, 2.1 mm × 100 mm, Waters, Milford, MA) set to 80°C and with a flow rate of 0.3 mL/min. Mobile phase A was water with 0.1% formic acid (Thermo) and mobile phase B was acetonitrile with 0.1% formic acid (Thermo). The gradient was as follows: 0 min, 10%B; 3 min, 10%B; 3.1 min, 25%B; 33 min, 45%B; 33.1 min, 95%B; 36 min, 95%B; 36.1 min, 10%B; 40 min, 10%B. The column eluate was analyzed by mass spectrometry (Xevo TQ-S, Waters, Milford, MA or Q Exactive HF, Thermo). Mass spectra were processed with Mass Lynx (Waters Xevo data) or BioPharma Finder (Thermo QE-HF data). Three antibodies (42056-a.01, 44715-a.01, and 6561-a.01) had as many as four sulfation proteoforms detected per digested F(ab’)^2^ molecule, indicating two sites of tyrosine sulfation on each Fab. For 6561-a.01, we modeled both sulfated tyrosines at positions Y100bHCDR3 and Y100fHCDR3. For 44715-a.01, we modeled just one sulfated tyrosine at position Y100fHCDR3, but suspect the second sulfation modification to occur at adjacent residue position Y100gHCDR3. For 42056-a.01, we modeled just one sulfated tyrosine at position Y100cHCDR3, but suspect the second sulfation modification to occur at adjacent residue position Y100dHCDR3. We unexpectedly found many *O*-linked glycoforms for each rhesus antibody irrespective of the presence or absence of tyrosine sulfation, but not in for the human antibody controls PGDM1400 and ACS202. There was a trend for tyrosine sulfation to be increased (both the number of sulfation groups and relative intensity of mass peaks) in the presence of *O*-linked glycosylation for the rhesus antibodies.

### Quantification and statistical analysis

The difference in mean values for distributions of frequency, HCDR3 length, and HCDR3 net charge for DH3 family-derived naïve rhesus B cells was calculated with Student’s t-test. The difference in mean values for Fab-V2 apex interactive surface features grouped by species and mode of recognition was calculated using Unpaired t-test. These statsitical analyses were performed using GraphPad Prism 10.

## Summary of Supplementary Information

### Figures

**Figure S1.**
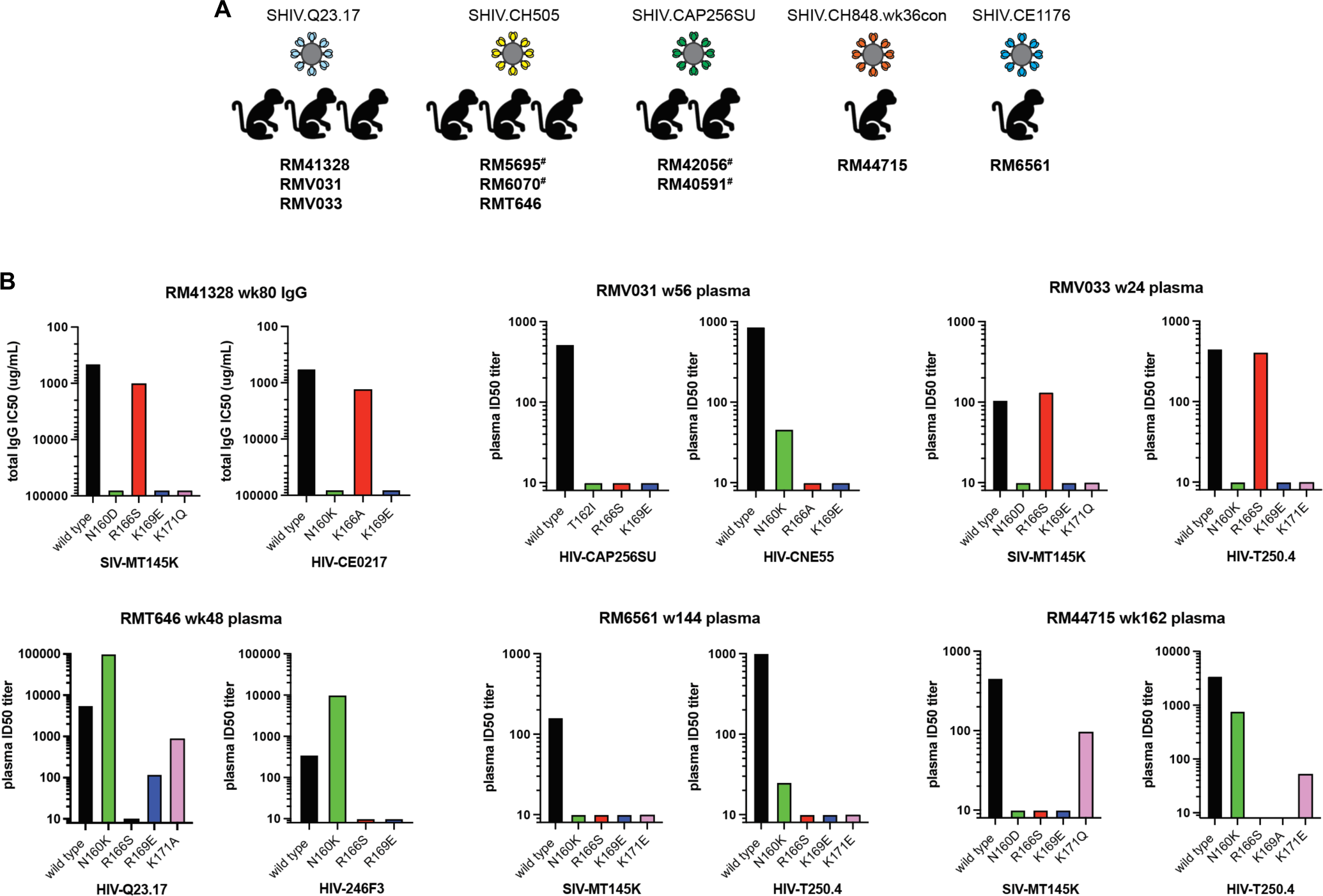
Identification of ten SHIV-infected rhesus macaques with HIV-1 V2 apex-targeted heterologous neutralization breadth. (A) Rhesus macaques from which V2 apex lineages have been isolated in this study are grouped by their respective infecting SHIV strain. Animals previously described in Roark et al17 for polyclonal V2 apex mutational mapping are denoted with #. (B) Neutralization of two sets of heterologous wild type and V2 apex epitope mutant viruses by rhesus macaque plasma or purified polyclonal IgG (RM41328) for SHIV-infected rhesus macaques reported in this study.

**Figure S2.**
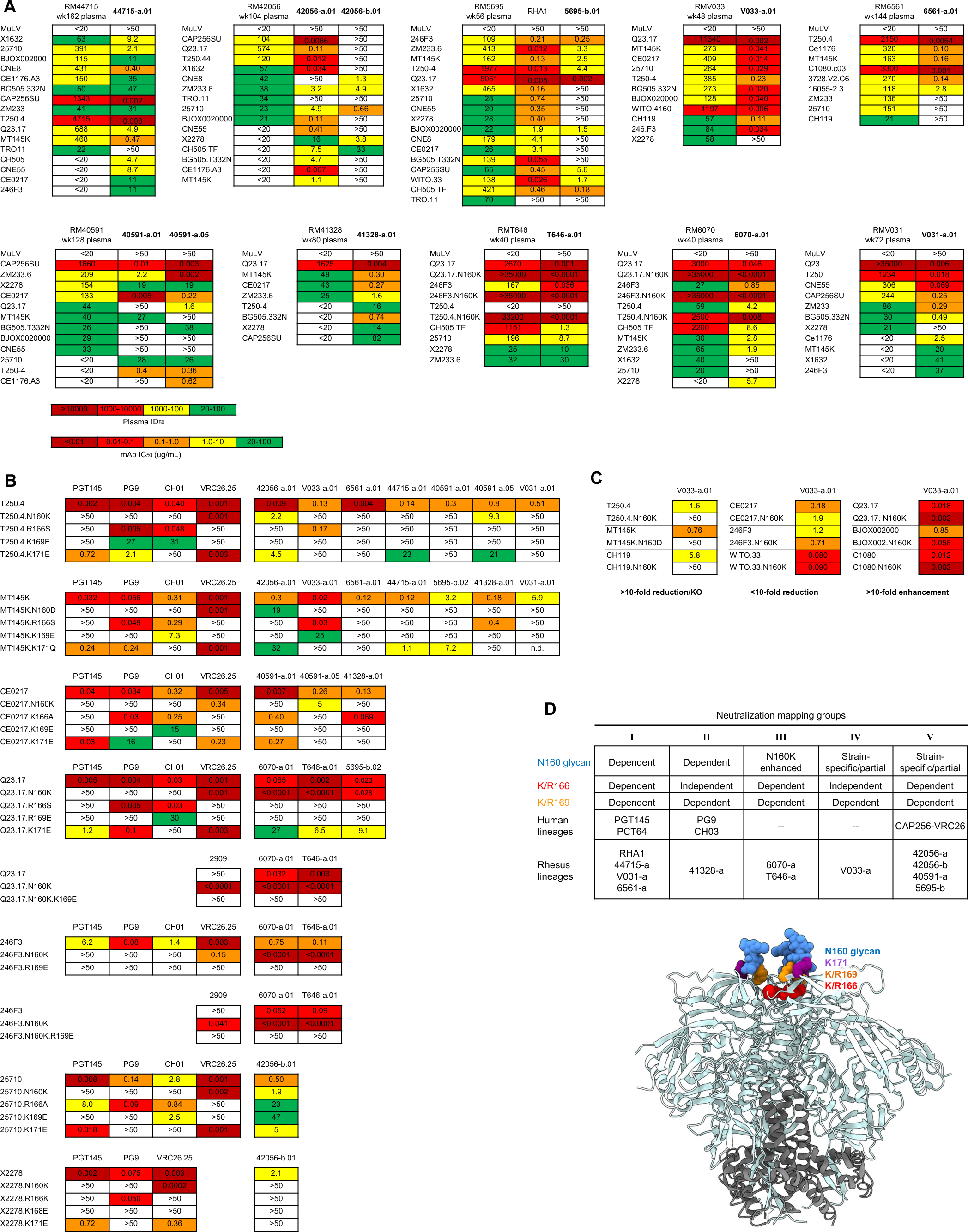
Isolated lineages with long, anionic HCDR3s recapitulate donor plasma heterologous neutralization breadth by targeting the HIV-1 V2 apex. (A) Neutralization of diverse tier-2 viruses by paired donor rhesus macaque plasma and representative lineage member mAbs. MuLV is murine leukemia virus, a negative control for neutralization. Data is reported as reciprocal ID50 titer for plasma and IC50 (ug/mL) titer for mAbs. The color legend for IC50 values also applies panels B and C below. (B) Neutralization of heterologous wild type and V2 apex epitope mutant viruses by representative rhesus and human lineage mAbs. For each set of viruses, human control mAbs are grouped on the left and rhesus mAbs are grouped on the right. Data is reported as IC50 (ug/mL) titer. Not determined = n.d. (C) Neutralization of heterologous wild type and N160 glycan deficient (N160K or N160D) mutant viruses by V033-a.01. Virus pairs are grouped by the effect of N160 glycan removal on neutralization sensitivity compared to wild type. Data is reported as IC50 (ug/mL) titer. (D) Potent rhesus and human V2 apex-targeted lineages can be divided into five neutralization groups (I-V) based on mutant virus epitope mapping. Groups III and IV have not been previously described. The envelope trimer (PDB-4ZMJ) highlights the location of *N*-linked glycan and protein residue substitutions used for V2 apex mapping.

**Figure S3.**
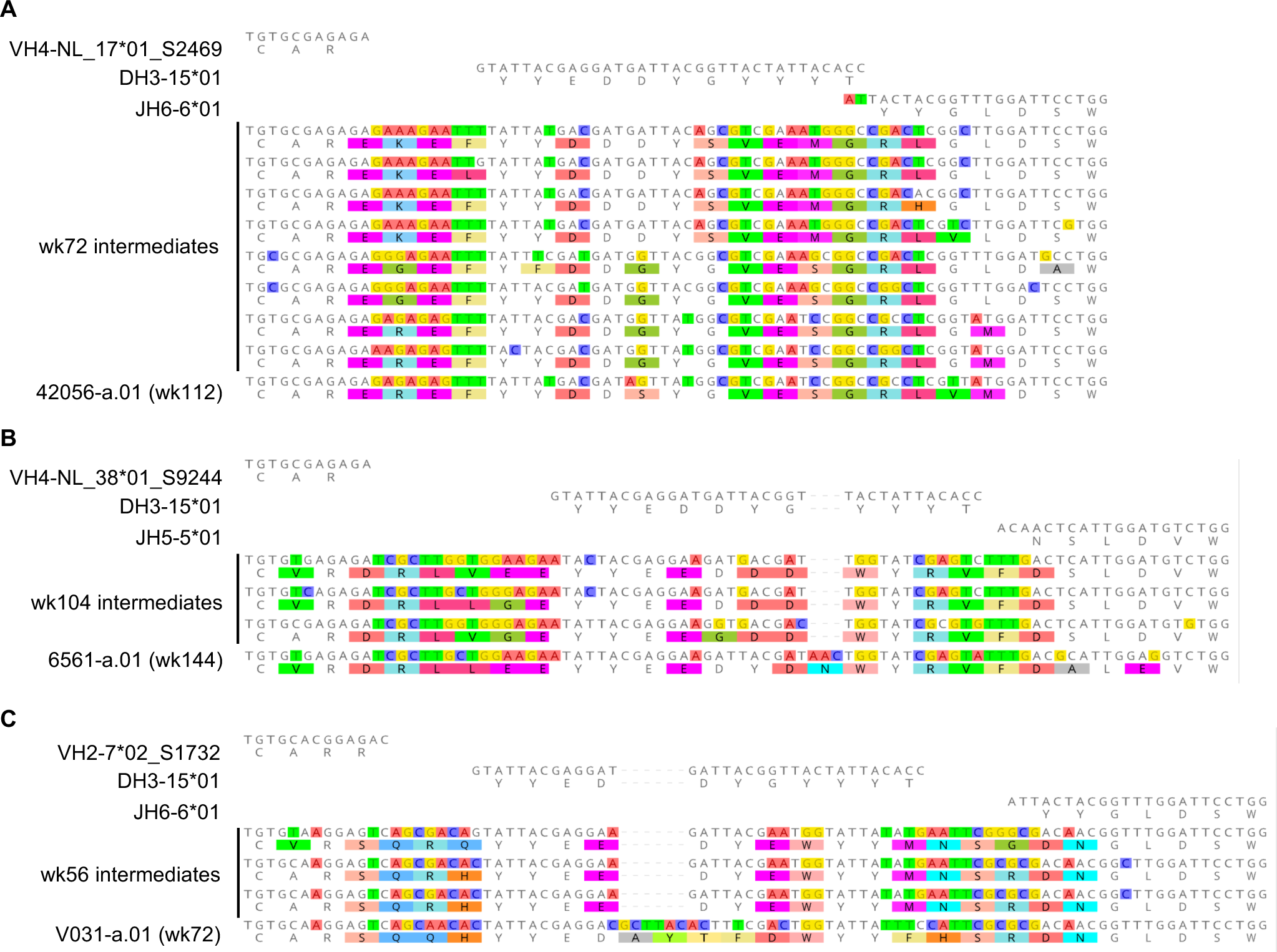
Ancestral clonal sequences identify HCDR3 indels and VDJ gene contributions and confirm acquisition of fiveresidue EDDYG motif for rhesus lineages with significant somatic mutation within D gene segments. HCDR3 alignments of the rhesus antibodies (A) 42056-a.01, (B) 6561-a.01, and (C) V031-a.01 to their respective VDJ germline genes and ancestral lineage intermediate sequences identified from peripheral memory B cells at the indicated time points. Mismatches to the germline V, D, and J genes are highlighted.

**Figure S4.**
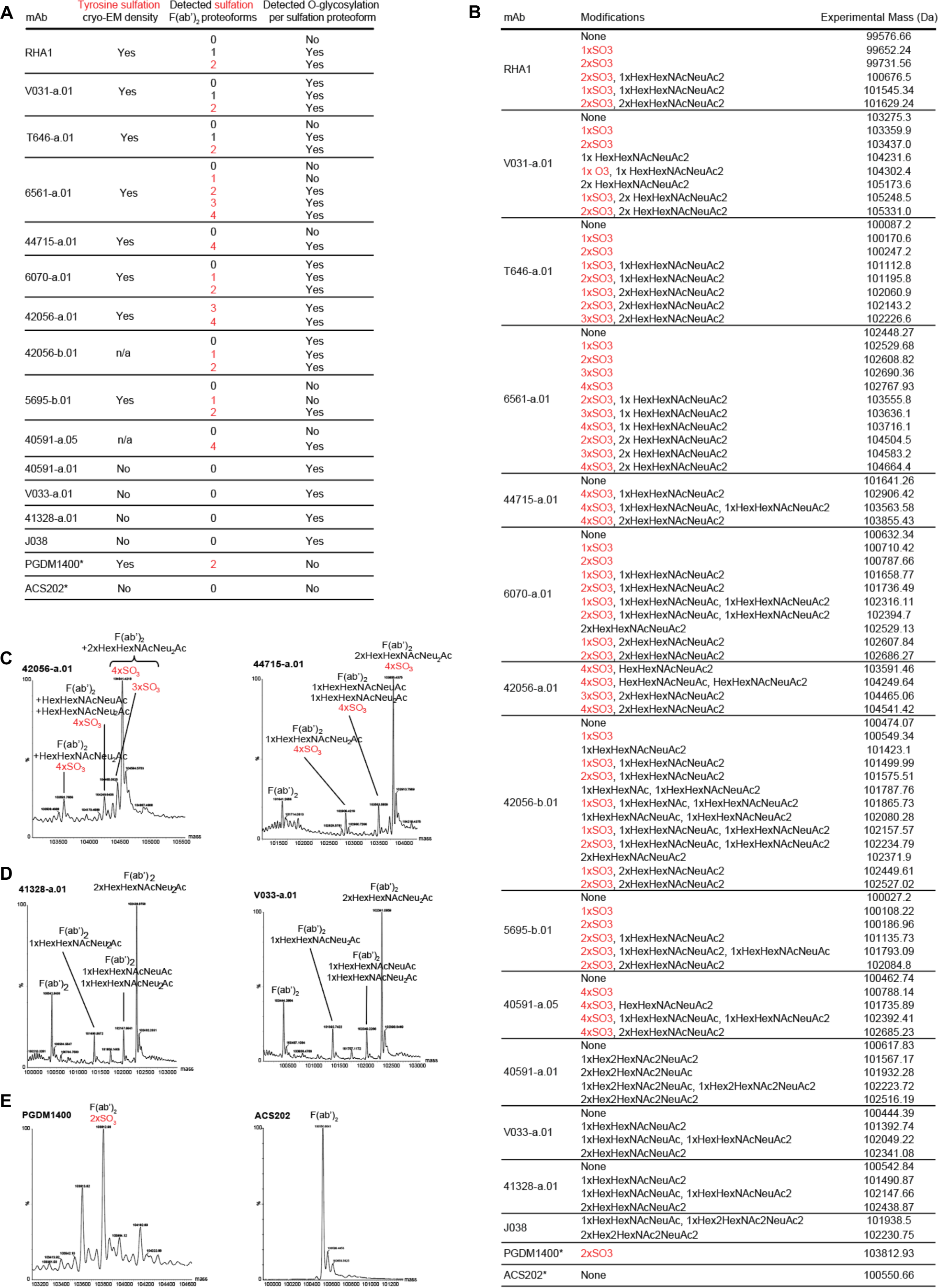
Rhesus V2 apex lineages bear tyrosine sulfation and *O*-linked glycosylation posttranslational modifications. (A) Summary of posttranslational modifications detected on F(ab’)2-digested rhesus and human antibodies by mass spectroscopy. Human antibodies are denoted with *. The number of sulfation groups detected per proteoform are written in red. All rhesus antibodies contained various types of *O*-linked glycosylation, while human antibody controls did not. Antibodies without structural data are denoted with n/a (not applicable). (B) Full list of individual detected experimental masses and corresponding deconvoluted posttranslational modifications that were summarized in panel A. The number of sulfation groups detected per modification are written in red (#xSO3). (C) Representative deconvoluted mass spectra for rhesus lineage F(ab’)2 subunits with tyrosine sulfation peaks. Y axis, relative intensity. X axis, mass (Da). (D) Representative deconvoluted mass spectra for rhesus lineage F(ab’)2 subunits without tyrosine sulfation peaks. Y axis, relative intensity. X axis, mass (Da). (E) Deconvoluted mass spectra for human F(ab’)2-digested antibodies PGDM1400 and ACS202, which serve as positive and negative controls for tyrosine sulfation, respectively. Y axis, relative intensity. X axis, mass (Da).

**Figure S5.**
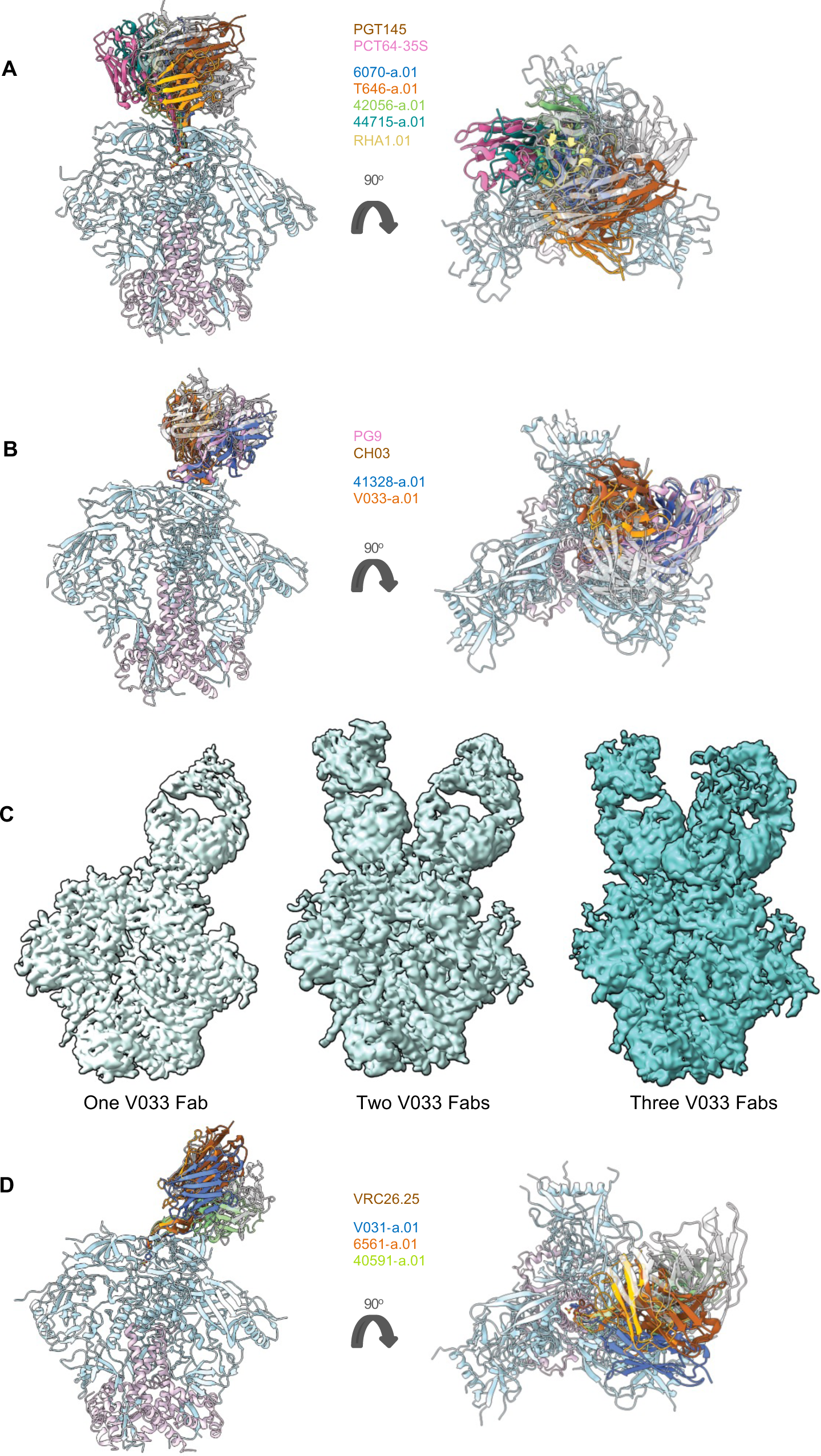
Rhesus and human lineages bind the V2 apex with a constellation of Fab orientations that do not segregate by species. Structures of Fab complexes with (A) PGT145-like, (B) PG9-like, and (D) VRC26-like modes of V2 apex recognition are aligned by gp120. Heavy chains are colored according to each respective legend and light chains are shown in transparent light gray. (C) Cryo-EM 3D reconstructions

**Figure S6.**
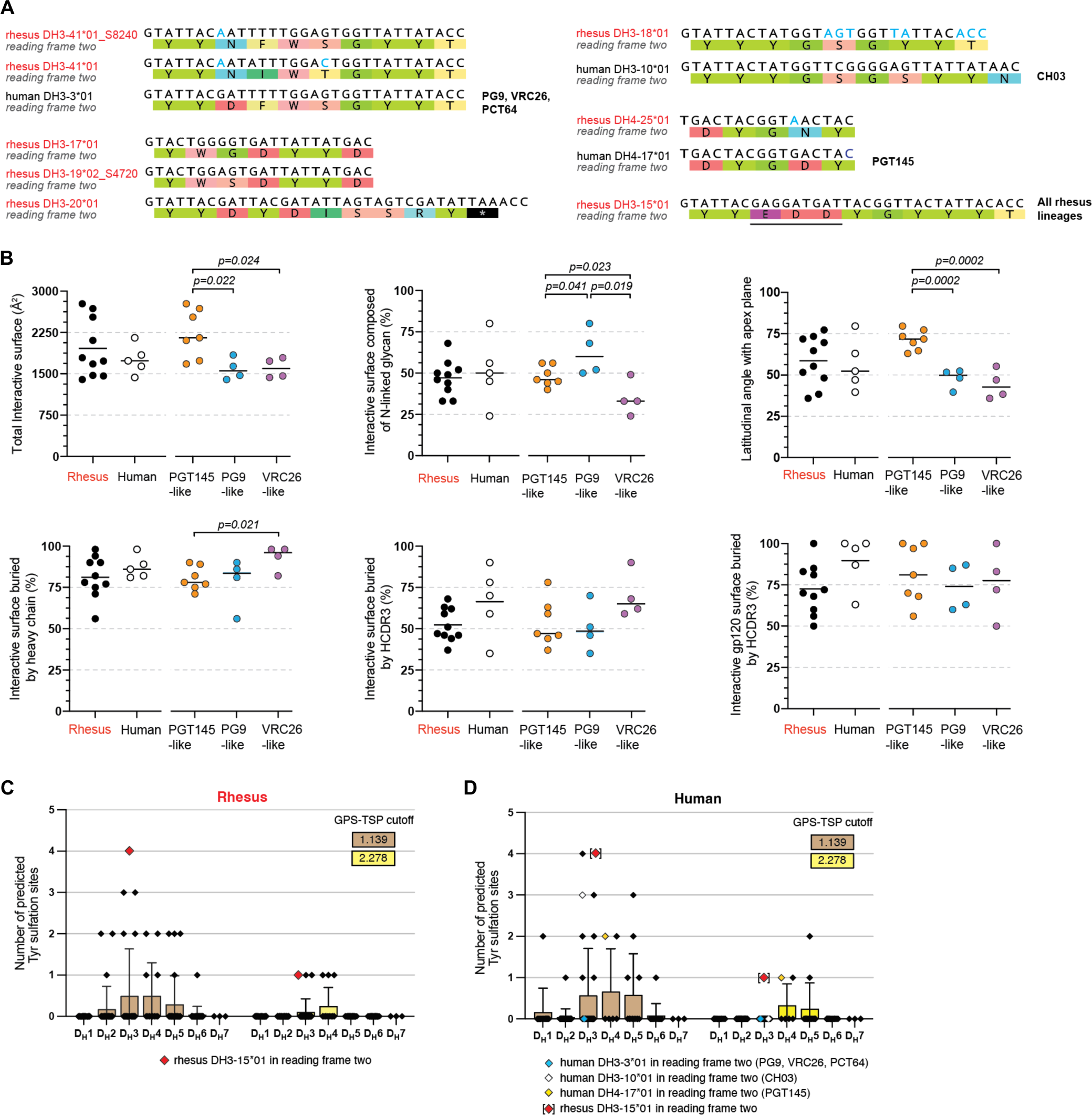
Rhesus macaque and human D gene repertoire analysis and features of V2 apex recognition. (A) Human V2 apex broadly neutralizing lineage D genes are aligned with their rhesus D gene homologs. Nucleotide differences in rhesus homologs are colored in blue. Non-homolog rhesus D genes encoding the “YYD” motif in human DH3-3*01 are included below that alignment. Rhesus DH3-15*01 is included for reference with the unique three-residue anionic motif (EDD) underlined. All rhesus sequence are labeled in red. (B) Epitope and paratope characteristics of V2 apex lineages. Each plot is a different structural feature that groups lineages by host species on the left and by mode of recognition on the right. P values are only listed for statistically significant differences (p<0.05, unpaired t-test) between two groups on the same side of the plot. (C,D) Predicted number of tyrosine sulfation sites using GPS-TSP41 for each rhesus and human D gene expressed in one of the three forward reading frames and modeled at the tip of a 15-residue HCDR3 loop. Results are recorded for the standard threshold/cutoff (1.139) and a high threshold/cutoff (2.278). Values are partitioned by D gene family and D genes used by broadly neutralizing lineages are marked according to the legends.

**Figure S7.**
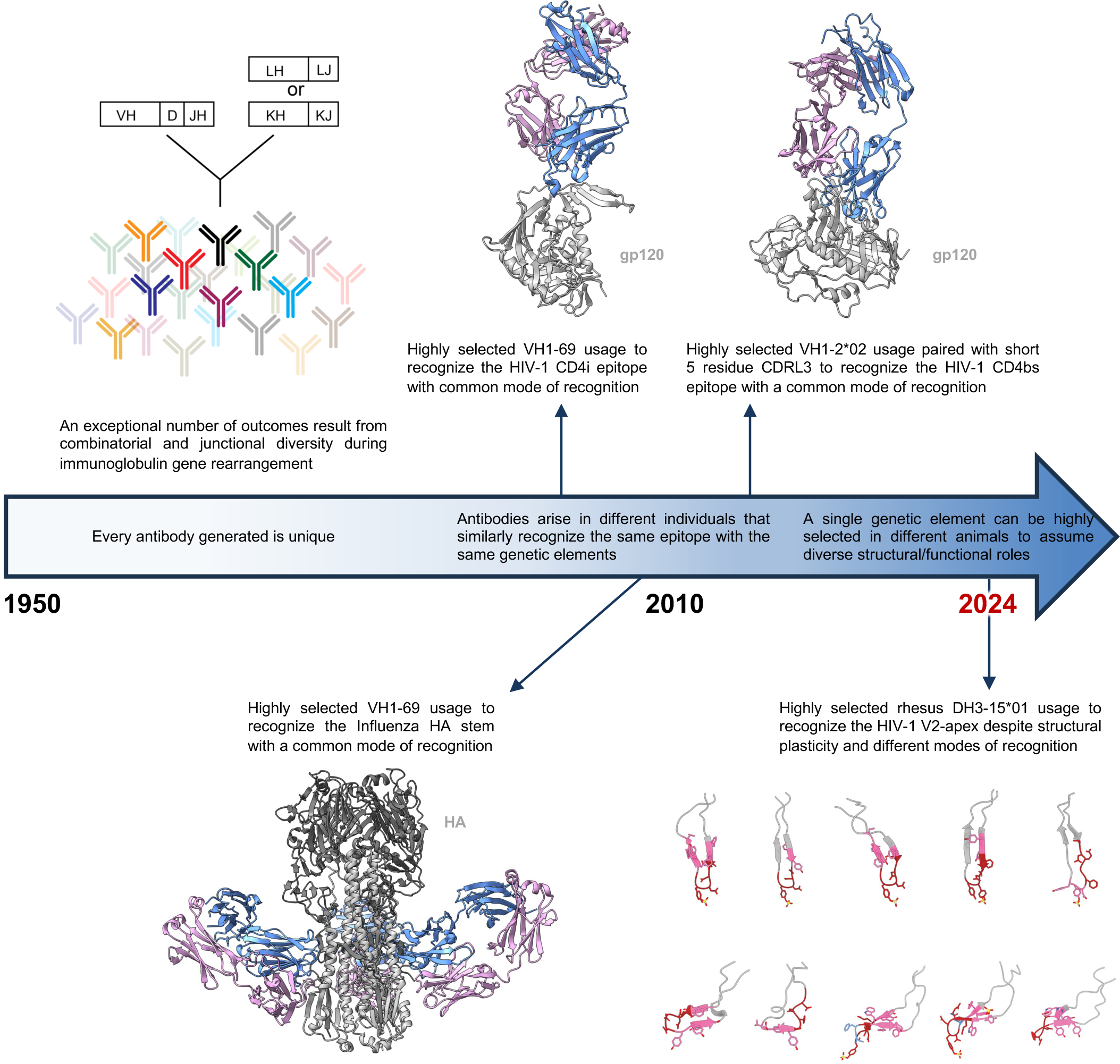
Highly selected immunogenetic elements can adopt diverse structural and functional roles in epitope recognition. A timeline of the field’s evolving understanding of antibody development and specificity. It was initially thought that the immense magnitude of possible outcomes from V(D)J+VJ recombination meant that every generated antibody would be unique. From 2004-2013, several studies established that reproducible multidonor antibody classes arise in the population due to highly selected genetic elements that are used to recognize the same HIV-1 gp12043-45,55 or Influenza HA52-54 epitopes through highly-similar, often identical, structural modes of recognition. Thus, not all antibodies are unique. Now, we have discovered that it is possible for highly selected genetic elements to exhibit structural and functional plasticity.

### Tables

**Table S1.**
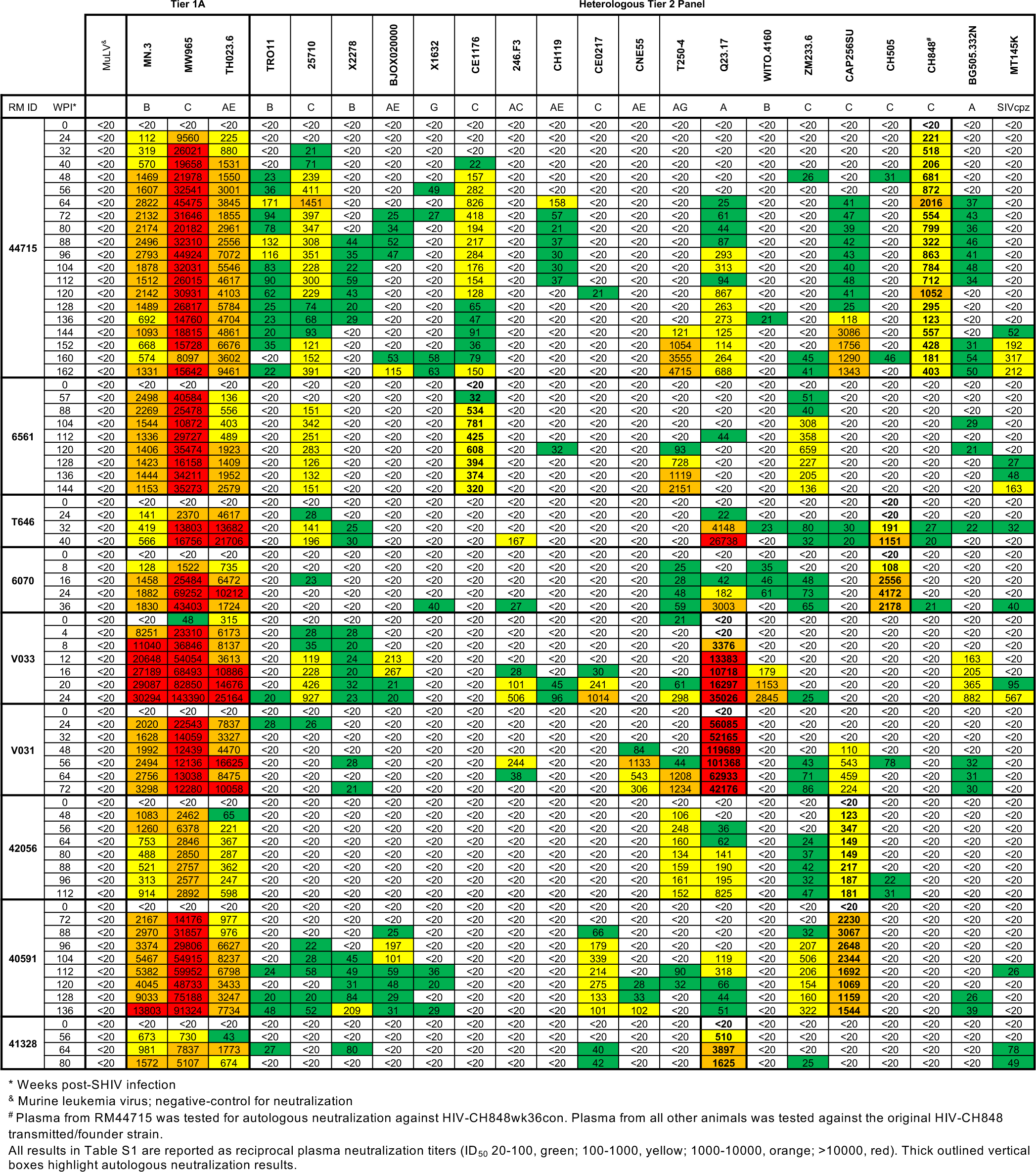
Autologous and heterologous neutralization by plasma from SHIV-infected rhesus macaques with V2 apex-targeted responses.

**Table S2.**
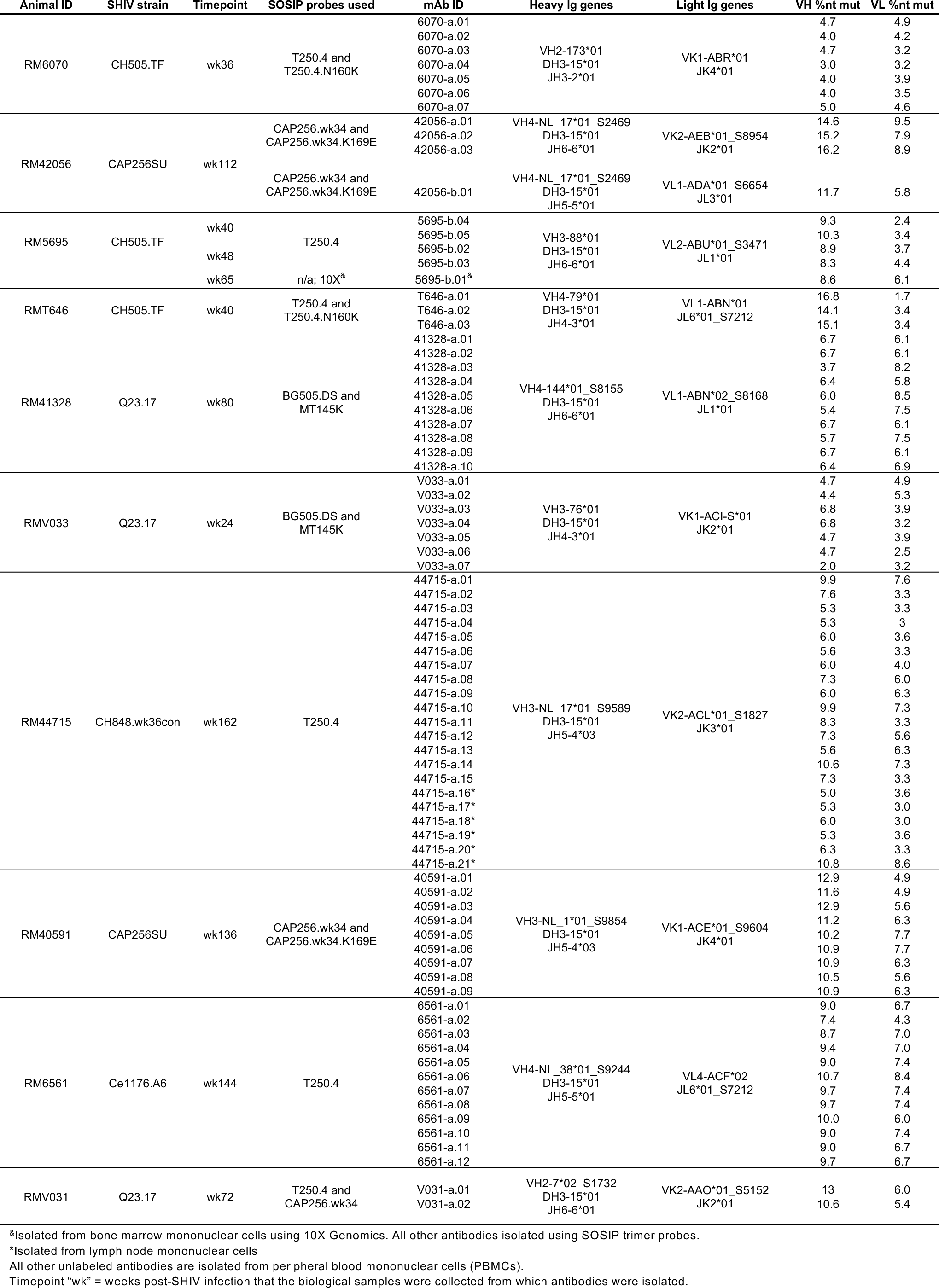
Animal information, isolation strategy, and immunogenetics of rhesus V2 apex lineages.

**Table S3.**
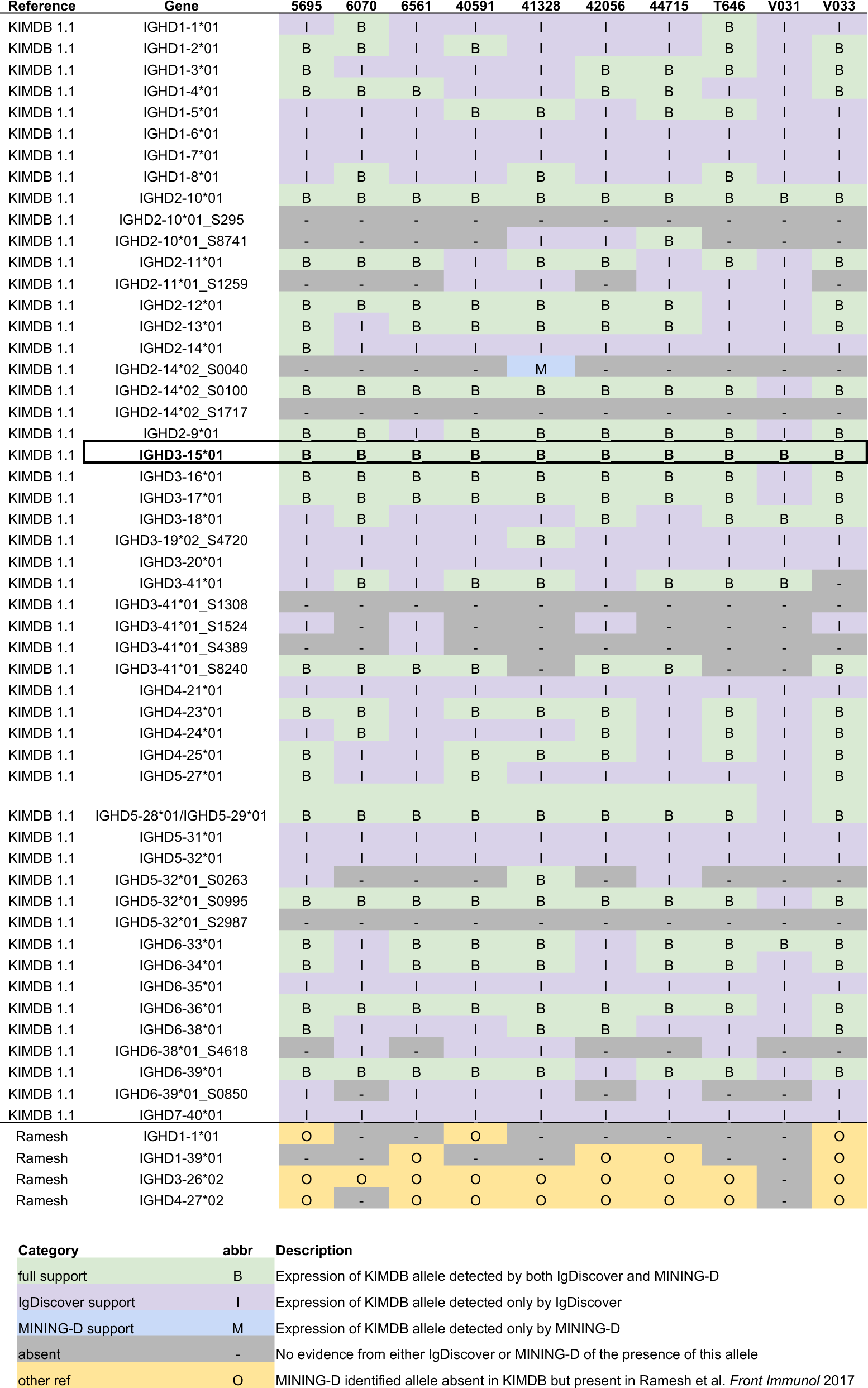
Personalized heavy and light chain immunoglobulin gene repertoires for rhesus macaques from which V2 apex lineages were isolated.

**Table S4.**
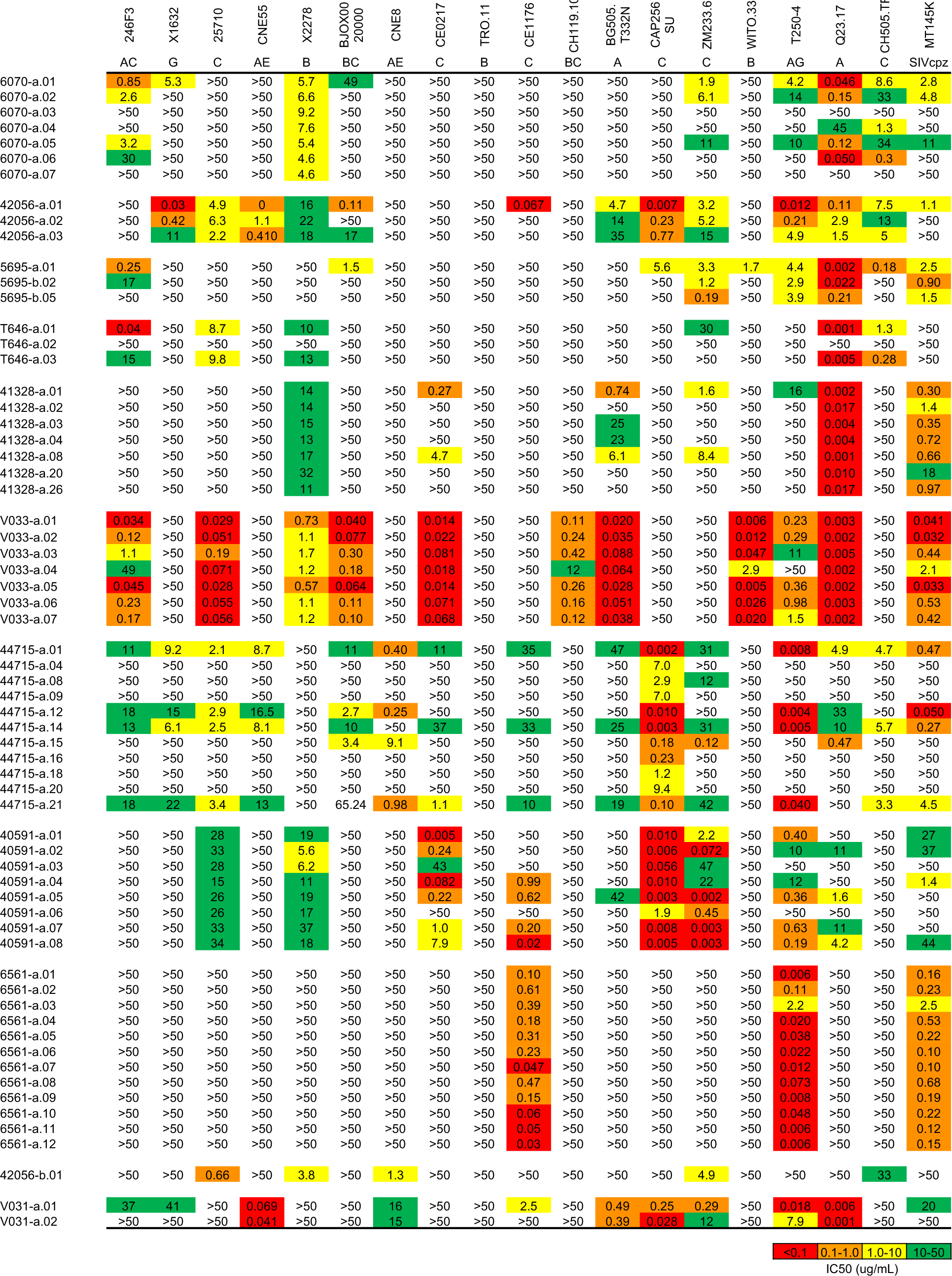
Neutralization titers against tier-2 heterologous viral strains by rhesus V2 apex lineages.

**Table S5.**
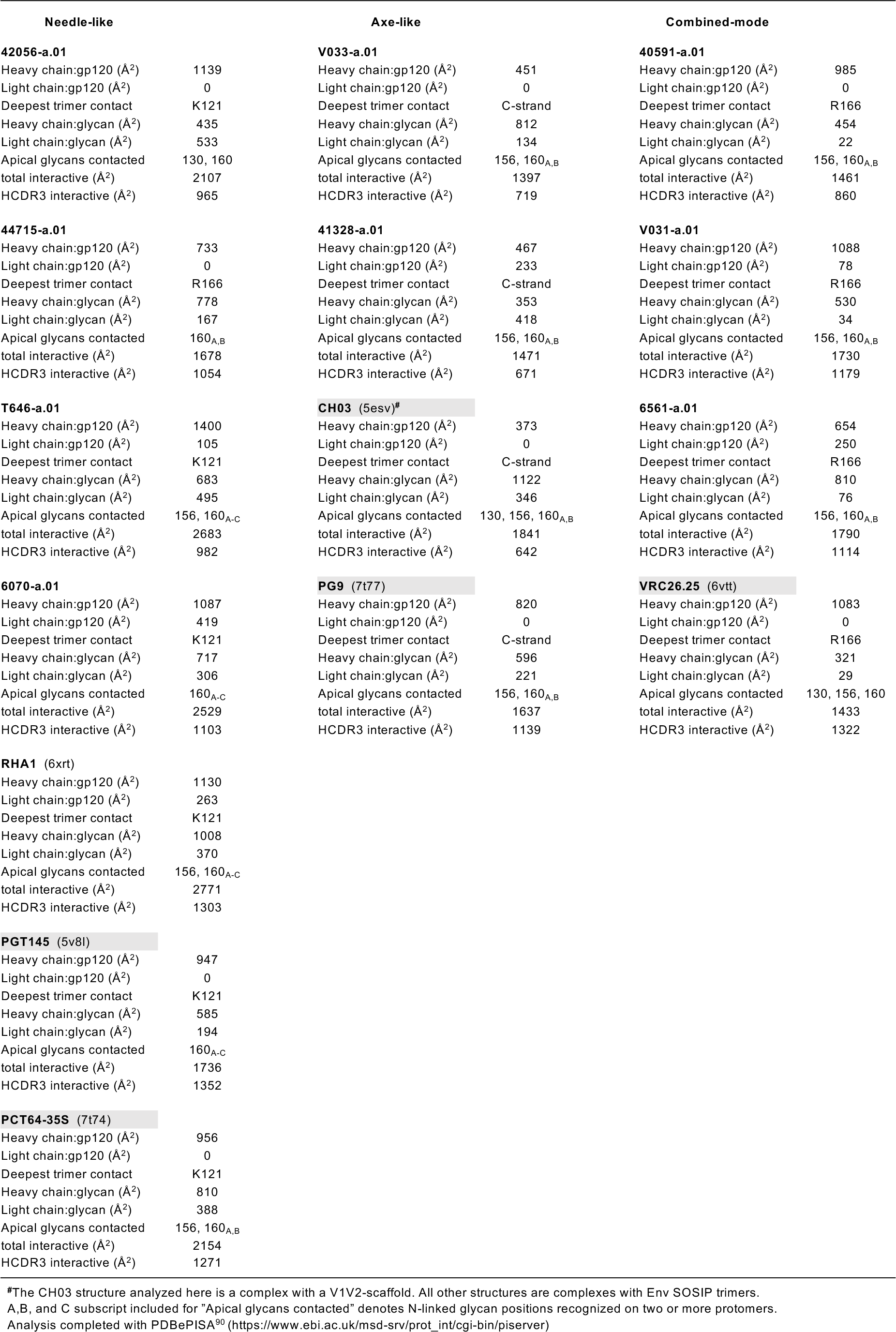
Select structural features of rhesus and human V2 apex broadly neutralizing antibody interactive surfaces.

**Table S6.** Cryo-EM data collection and refinement statistics.

|  | 6070-a.01<br>in complex with<br>Q23.17 MD39 SOSIP | T646-a.01<br>in complex with<br>Q23.17 MD39 SOSIP | 42056-a.01<br>in complex with<br>CAP256 wk34 c80<br>RnS2 SOSIP | 44715-a.01<br>in complex with<br>BG505 DS-SOSIP | 41328-a.01<br>in complex with<br>BG505 DS-SOSIP | V033-a.01<br>in complex with<br>BG505 DS-SOSIP | V031-a.01<br>in complex with<br>BG505 DS-SOSIP | 6561-a.01<br>in complex with<br>Ce1176 RnS2 SOSIP | 40591-a.01<br>in complex with<br>T250.4 RnS2 SOSIP |
| --- | --- | --- | --- | --- | --- | --- | --- | --- | --- |
| PDB | 9BNL | 9BTV | 9BTH | 9BNM | 9BTL | 9BNP | 9BNK | 9BTJ | 9BTI |
| EMDB | 44729 | 44897 | 44890 | 44730 | 44893 | 44733 | 44728 | 44892 | 44891 |
| **Data collection & processing** |  |  |  |  |  |  |  |  |  |
| Microscope | FEI Titan Krios | FEI Titan Krios | FEI Titan Krios | FEI Titan Krios | FEI Titan Krios | FEI Titan Krios | FEI Titan Krios | FEI Titan Krios | FEI Titan Krios |
| Camera | Gatan K3 | Gatan K3 | Gatan K3 | Gatan K3 | Gatan K3 | Gatan K3 | Gatan K3 | Gatan K2 Summit | Gatan K3 |
| Magnification | 105,000x | 105,000x | 64,000x | 105,000x | 105,000x | 105,000x | 105,000x | 22,500x | 64,000x |
| Voltage (kV) | 300 | 300 | 300 | 300 | 300 | 300 | 300 | 300 | 300 |
| Electron dose (e<sup>-</sup>/Å<sup>2</sup>) | 58 | 58 | 56 | 58 | 58 | 58 | 58 | 69 | 56 |
| Defocus range (µm) | 0.8 - 2.0 | 0.8 - 2.0 | 1.0 - 2.5 | 0.8 - 2.0 | 0.8 - 2.0 | 0.8 - 2.0 | 0.8 - 2.0 | 1.0 - 2.5 | 1.0 - 2.5 |
| Pixel size (Å) | 0.83 | 0.83 | 1.08 | 0.83 | 0.83 | 0.83 | 0.83 | 1.073 | 1.08 |
| Micrographs collected | 2,406 | 5,358 | 2,385 | 4,736 | 5,074 | 4,522 | 5,467 | 1,357 | 3,666 |
| Software | cryoSPARC v3.2 | cryoSPARC v3.2 | cryoSPARC v4.1 | cryoSparc v4.1 | cryoSPARC v3.2 | cryoSPARC v3.2 | cryoSparc v4.1 | cryoSPARC v3.0 | cryoSPARC v4.1 |
| Micrographs used | 2,187 | 5,225 | 2,212 | 3,748 | 4,817 | 4,505 | 5,074 | 1,357 | 3,318 |
| Total extracted particles | 648,715 | 1,601,119 | 553,329 | 757,694 | 2,403,591 | 2,214,775 | 1,256,494 | 521,302 | 748,766 |
| Refined particles | 53,426 | 39,411 | 69,264 | 48,848 | 176,070 | 123,355 | 353,856 | 150,641 | 66,103 |
| Symmetry imposed | C1 | C1 | C1 | C1 | C1 | C1 | C1 | C1 | C1 |
| Map Resolution (Å) | 3.66 | 3.48 | 4.20 | 3.97 | 2.96 | 3.17 | 3.10 | 4.22 | 4.14 |
| FSC threshold | 0.143 | 0.143 | 0.143 | 0.143 | 0.143 | 0.143 | 0.143 | 0.143 | 0.143 |
| **Refinement & validation** |  |  |  |  |  |  |  |  |  |
| Initial model used | 7LLK | 7LLK | 6VTT | 6XRT | 4ZMJ | 4ZMJ | 6XRT | 6VTT | 6VTT |
| Software | Phenix 1.20 | Phenix 1.20 | Phenix 1.21 | Phenix 1.21 | Phenix 1.20 | Phenix 1.20 | Phenix 1.21 | Phenix 1.21 | Phenix 1.21 |
| Number of residues |  |  |  |  |  |  |  |  |  |
| Protein | 1,911 | 1,915 | 1,914 | 1,917 | 1,930 | 1,920 | 1,966 | 2,010 | 1,919 |
| Ligand | 120 | 126 | 111 | 101 | 106 | 107 | 93 | 159 | 92 |
| Map CC | 0.84 | 0.80 | 0.78 | 0.74 | 0.86 | 0.82 | 0.86 | 0.79 | 0.73 |
| R.m.s. deviations |  |  |  |  |  |  |  |  |  |
| Bond lengths (Å) | 0.005 | 0.005 | 0.003 | 0.007 | 0.005 | 0.006 | 0.005 | 0.003 | 0.003 |
| Bond angles (°) | 1.14 | 0.96 | 0.725 | 1.18 | 1.21 | 1.19 | 0.96 | 0.67 | 0.652 |
| EMRinger score | 2.43 | 2.21 | 1.36 | 1.75 | 3.77 | 3.10 | 2.41 | 1.58 | 1.37 |
| MolProbity score | 1.55 | 1.60 | 1.69 | 1.56 | 1.55 | 1.44 | 1.59 | 1.59 | 1.6 |
| Clashscore | 4.21 | 4.61 | 4.16 | 4.13 | 4.33 | 3.65 | 4.75 | 4.86 | 4.11 |
| Rotamer outliers (%) | 0 | 0 | 0.29 | 0 | 0 | 0 | 0.97 | 0 | 0.18 |
| Ramachandran plot |  |  |  |  |  |  |  |  |  |
| Favored (%) | 95.0 | 94.7 | 90.64 | 94.6 | 95.1 | 95.9 | 95.1 | 95.14 | 93.95 |
| Allowed (%) | 5.0 | 5.3 | 8.88 | 5.4 | 4.9 | 4.1 | 4.9 | 4.66 | 5.68 |
| Outliers (%) | 0 | 0 | 0.48 | 0 | 0 | 0 | 0 | 0.2 | 0.37 |

### Data Files

Data S1. Cryo-EM data validations for HIV trimer – Fab complexes.

**Dataset S1 Figure 1.**
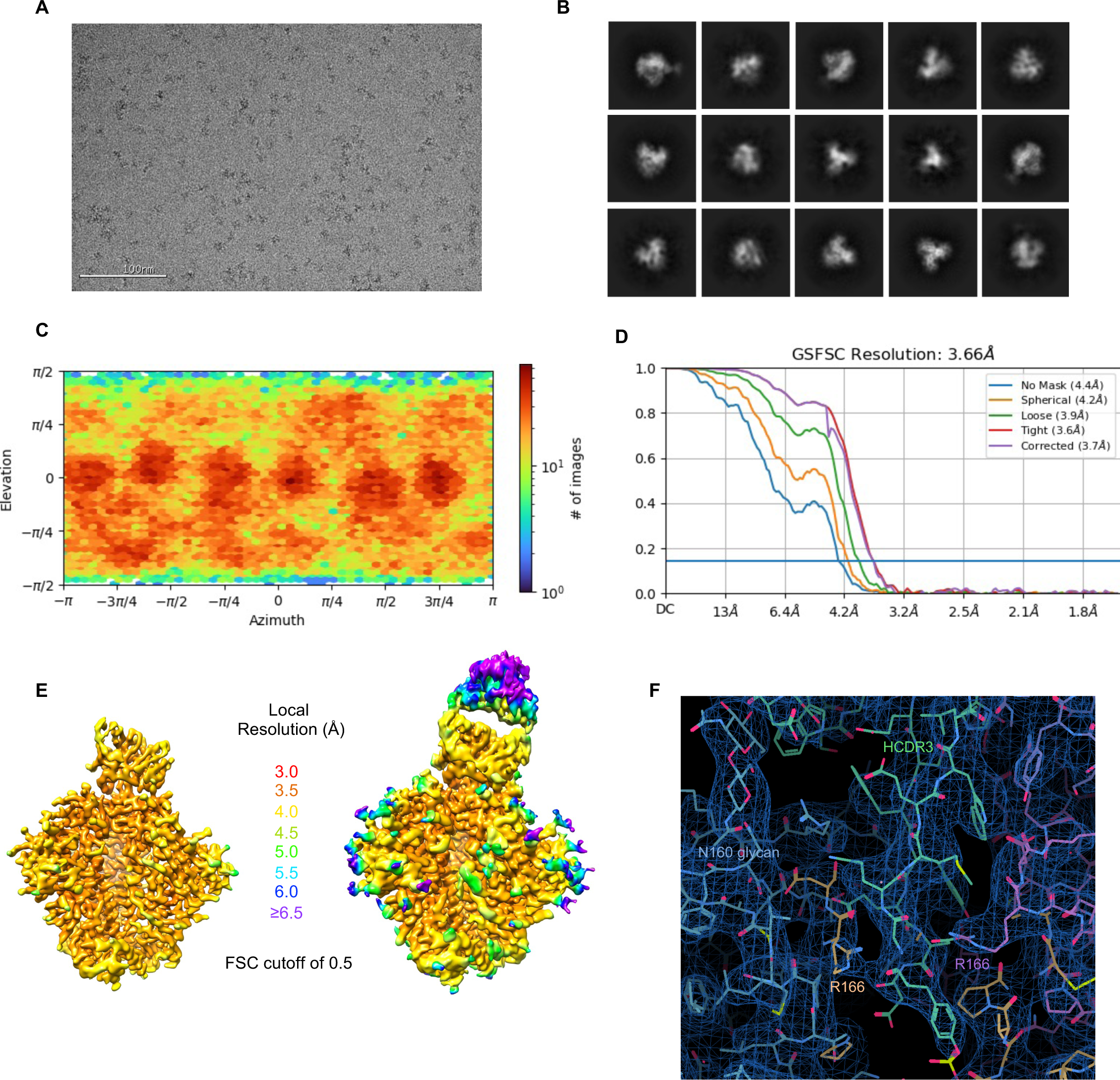
Cryo-EM details of 6070-a.01 in complex with Q23.17 MD39 SOSIP. (A) Representative raw micrograph is shown. (B) Representative 2D class averages of pick particles are shown. (C) The orientations of all particles used in the final refinement are shown as a heatmap. (D) The gold-standard fourier shell correlation (FSC) at threshold of 0.143 resulted in a resolution of 3.66 Å using non-uniform refinement with C1 symmetry. (E) The local resolution of the full map is shown as generated through cryoSPARC using an FSC cutoff of 0.5. Two volume contour levels are shown. (F) Cryo-EM 3D reconstruction density to highlight the Fab-trimer interactive surface.

**Dataset S1 Figure 2.**
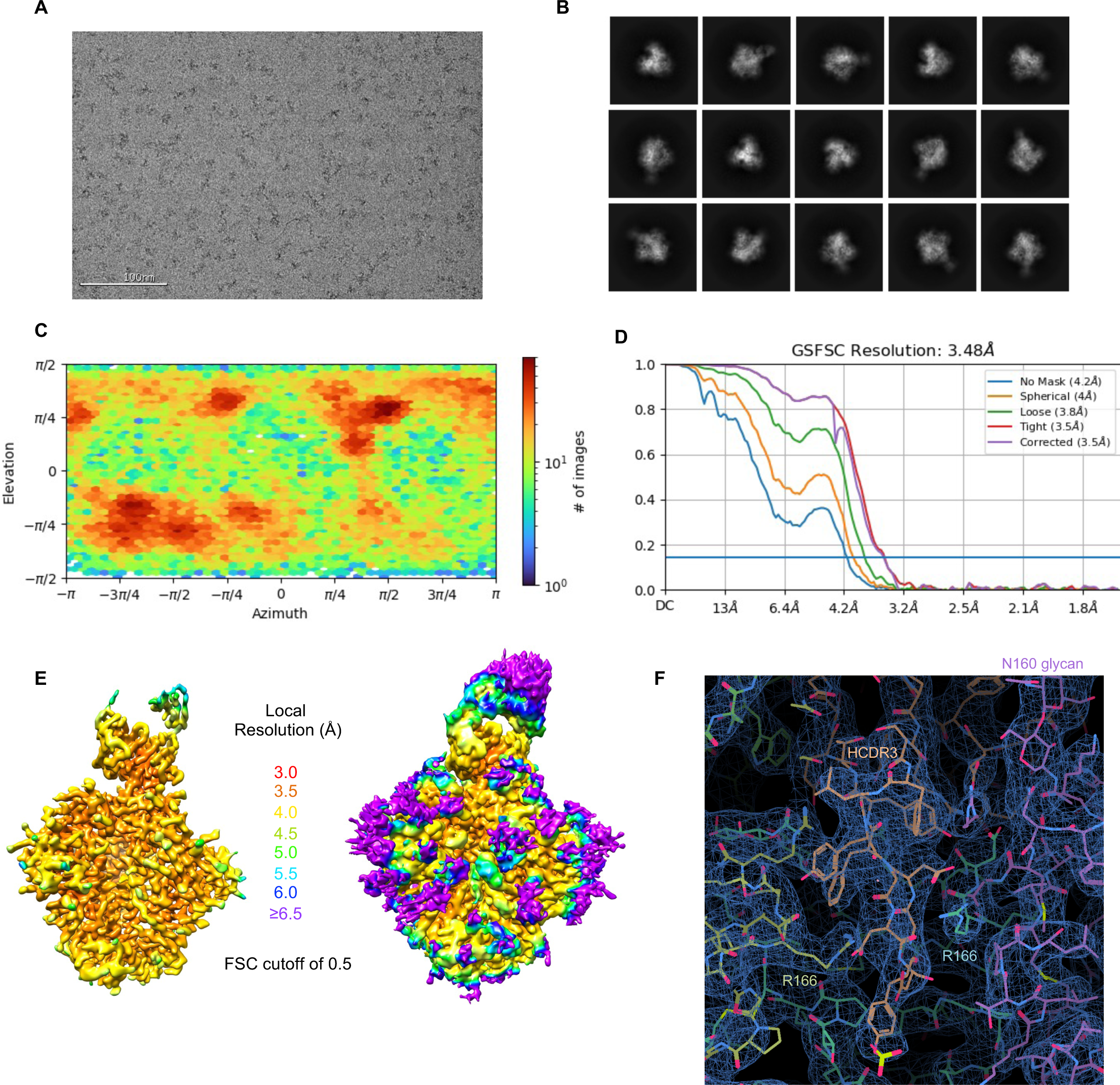
Cryo-EM details of T646-a.01 in complex with Q23.17 MD39 SOSIP. (A) Representative raw micrograph is shown. (B) Representative 2D class averages of pick particles are shown. (C) The orientations of all particles used in the final refinement are shown as a heatmap. (D) The gold-standard fourier shell correlation (FSC) at threshold of 0.143 resulted in a resolution of 3.48 Å using non-uniform refinement with C1 symmetry. (E) The local resolution of the full map is shown generated through cryoSPARC using an FSC cutoff of 0.5. Two volume contour levels are shown. (F) Cryo-EM 3D reconstruction density to highlight the Fab-trimer interactive surface.

**Dataset S1 Figure 3.**
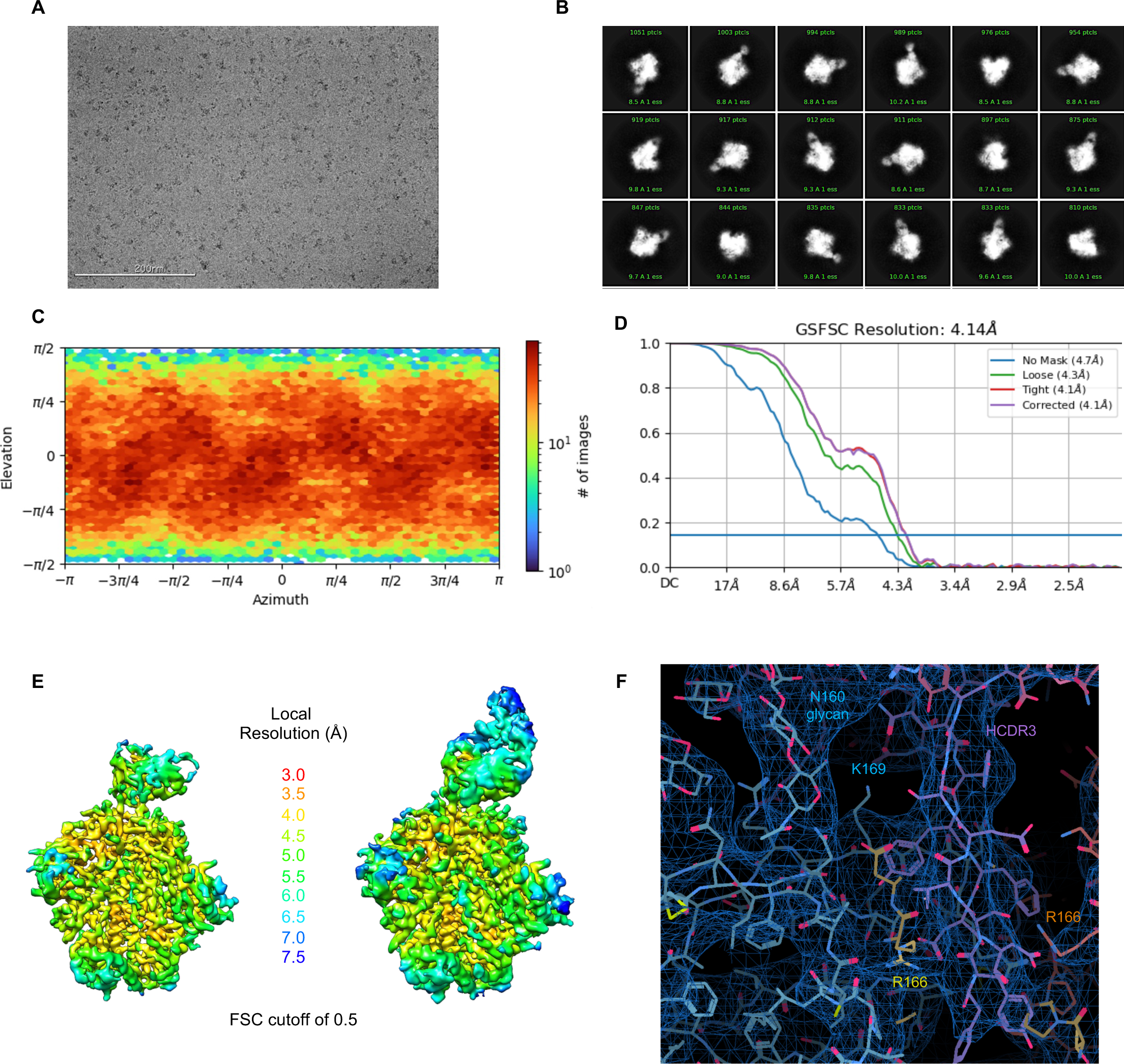
Cryo-EM details of 42056-a.01 in complex with CAP256.wk34.c80 RnS2 SOSIP. (A) Representative raw micrograph is shown. (B) Representative 2D class averages of pick particles are shown. (C) The orientations of all particles used in the final refinement are shown as a heatmap. (D) The gold-standard fourier shell correlation (FSC) at threshold of 0.143 resulted in a resolution of 4.20 Å using non-uniform refinement with C1 symmetry. (E) The local resolution of the full map is shown generated through cryoSPARC using an FSC cutoff of 0.5. Two volume contour levels are shown. (F) Cryo-EM 3D reconstruction density to highlight the Fab-trimer interactive surface.

**Dataset S1 Figure 4.**
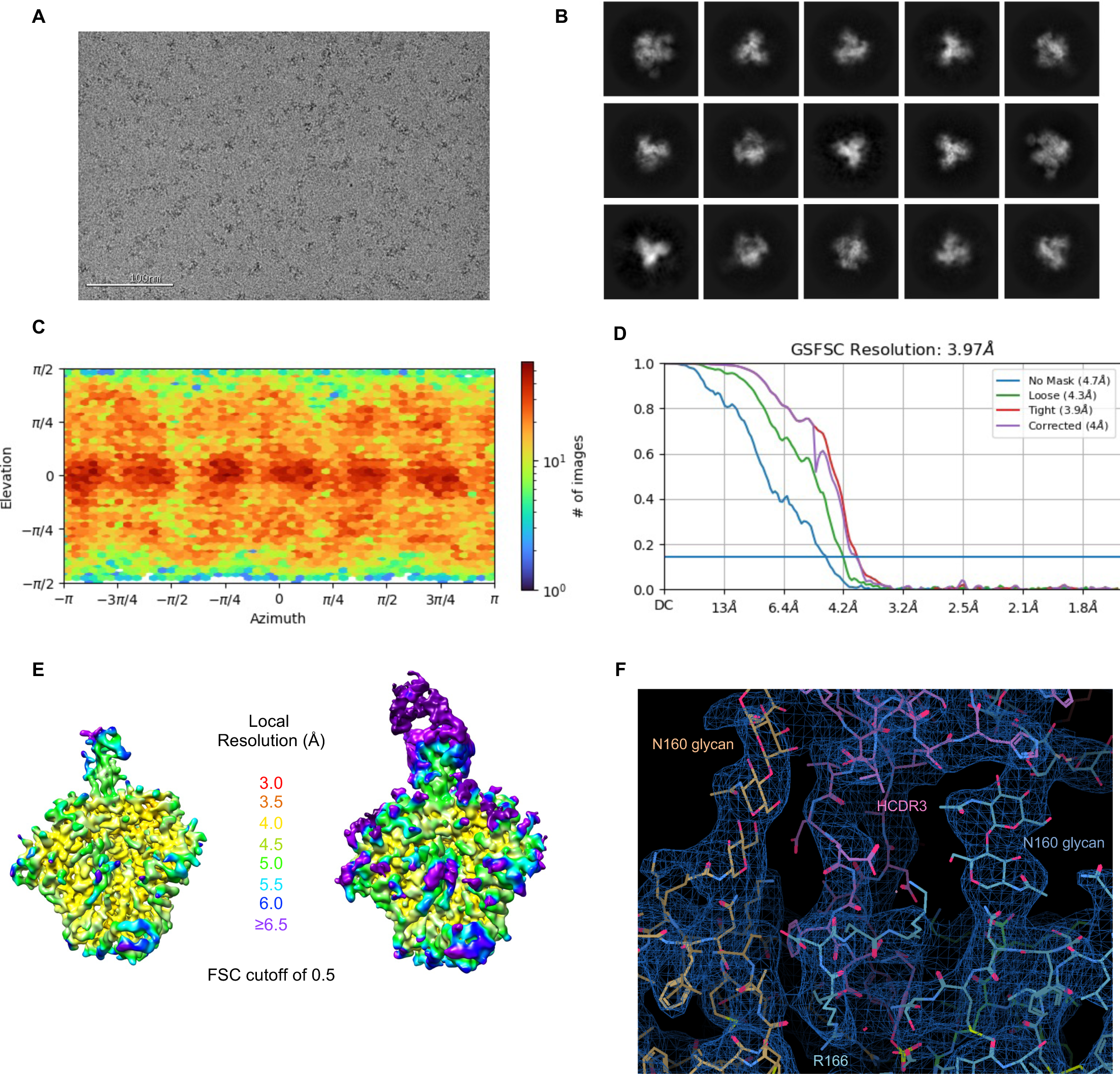
Cryo-EM details of 44715-a.01 in complex with BG505 DS-SOSIP. (A) Representative raw micrograph is shown. (B) Representative 2D class averages of pick particles are shown. (C) The orientations of all particles used in the final refinement are shown as a heatmap. (D) The gold-standard fourier shell correlation (FSC) at threshold of 0.143 resulted in a resolution of 3.97 Å using non-uniform refinement with C1 symmetry. (E) The local resolution of the full map is shown generated through cryoSPARC using an FSC cutoff of 0.5. Two volume contour levels are shown. (F) Cryo-EM 3D reconstruction density to highlight the Fab-trimer interactive surface.

**Dataset S1 Figure 5.**
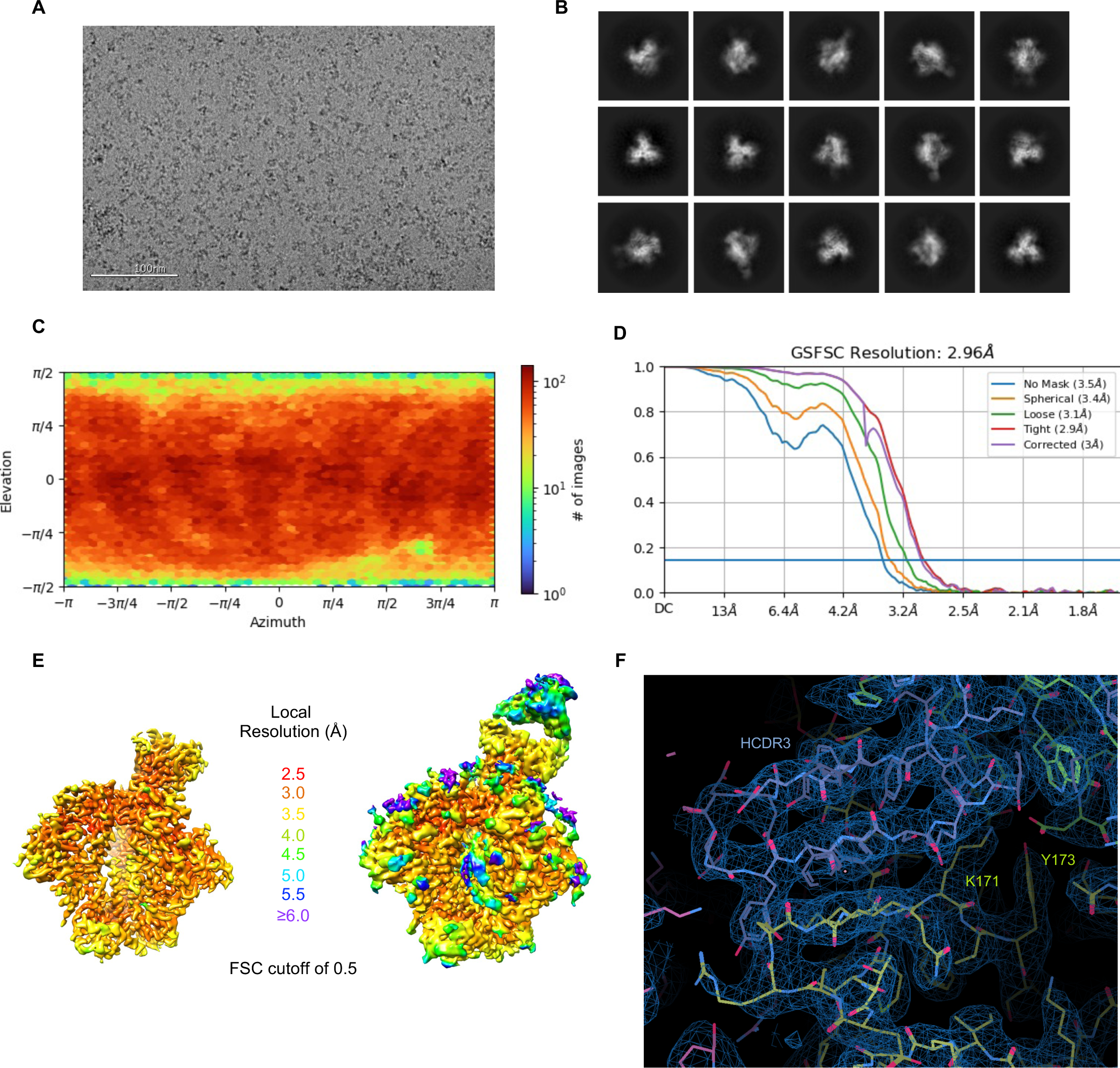
Cryo-EM details of 41328-a.01 in complex with BG505 DS-SOSIP. (A) Representative raw micrograph is shown. (B) Representative 2D class averages of pick particles are shown. (C) The orientations of all particles used in the final refinement are shown as a heatmap. (D) The gold-standard fourier shell correlation (FSC) at threshold of 0.143 resulted in a resolution of 2.96 Å using non-uniform refinement with C1 symmetry. (E) The local resolution of the full map is shown generated through cryoSPARC using an FSC cutoff of 0.5. Two volume contour levels are shown. (F) Cryo-EM 3D reconstruction density to highlight the Fab-trimer interactive surface.

**Dataset S1 Figure 6.**
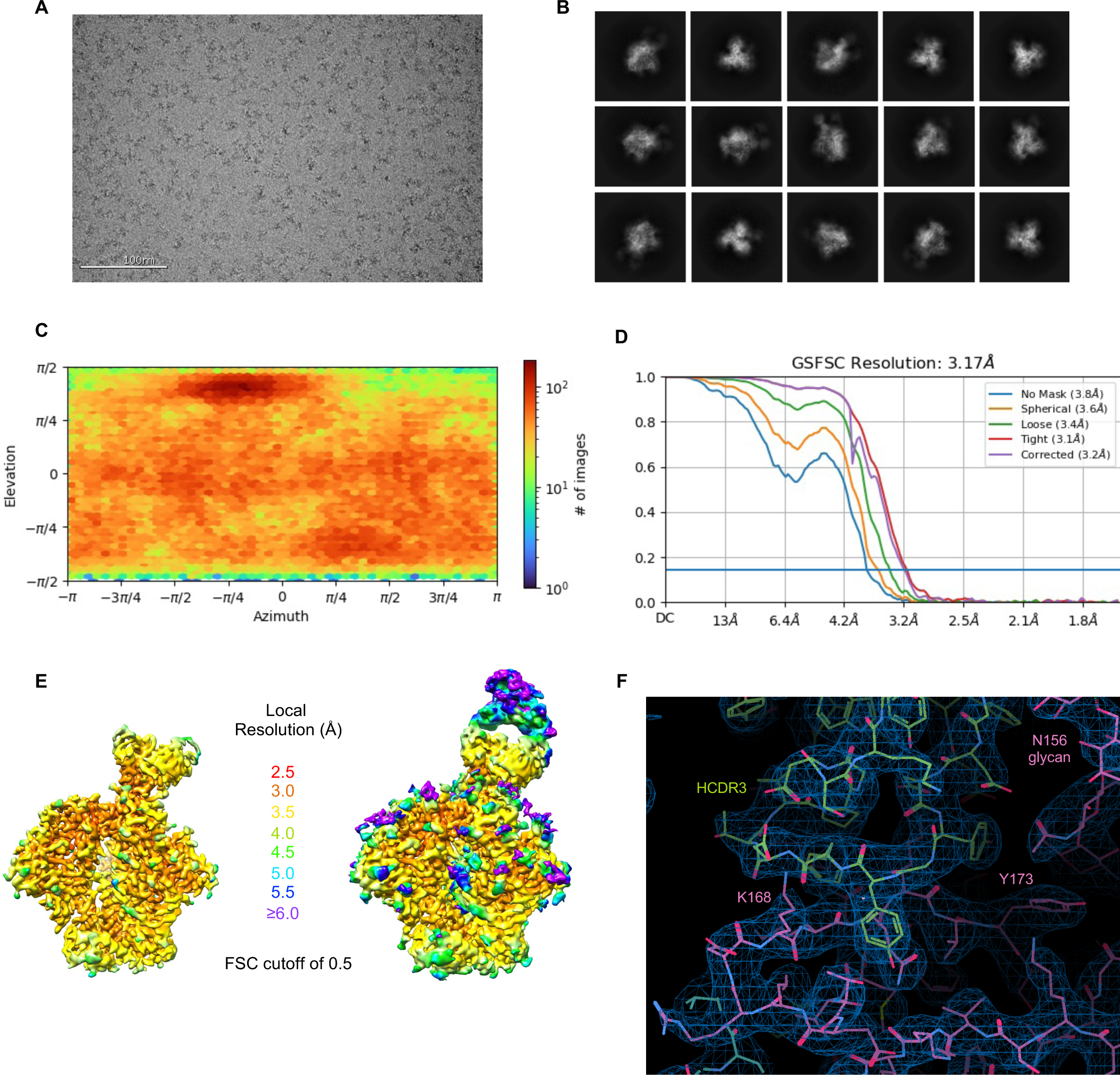
Cryo-EM details of V033-a.01 in complex with BG505 DS-SOSIP. (A) Representative raw micrograph is shown. (B) Representative 2D class averages of pick particles are shown. (C) The orientations of all particles used in the final refinement are shown as a heatmap. (D) The gold-standard fourier shell correlation (FSC) at threshold of 0.143 resulted in a resolution of 4.22 Å using non-uniform refinement with C1 symmetry. (E) The local resolution of the full map is shown generated through cryoSPARC using an FSC cutoff of 0.5. Two volume contour levels are shown. (F) Cryo-EM 3D reconstruction density to highlight the Fab-trimer interactive surface.

**Dataset S1 Figure 7.**
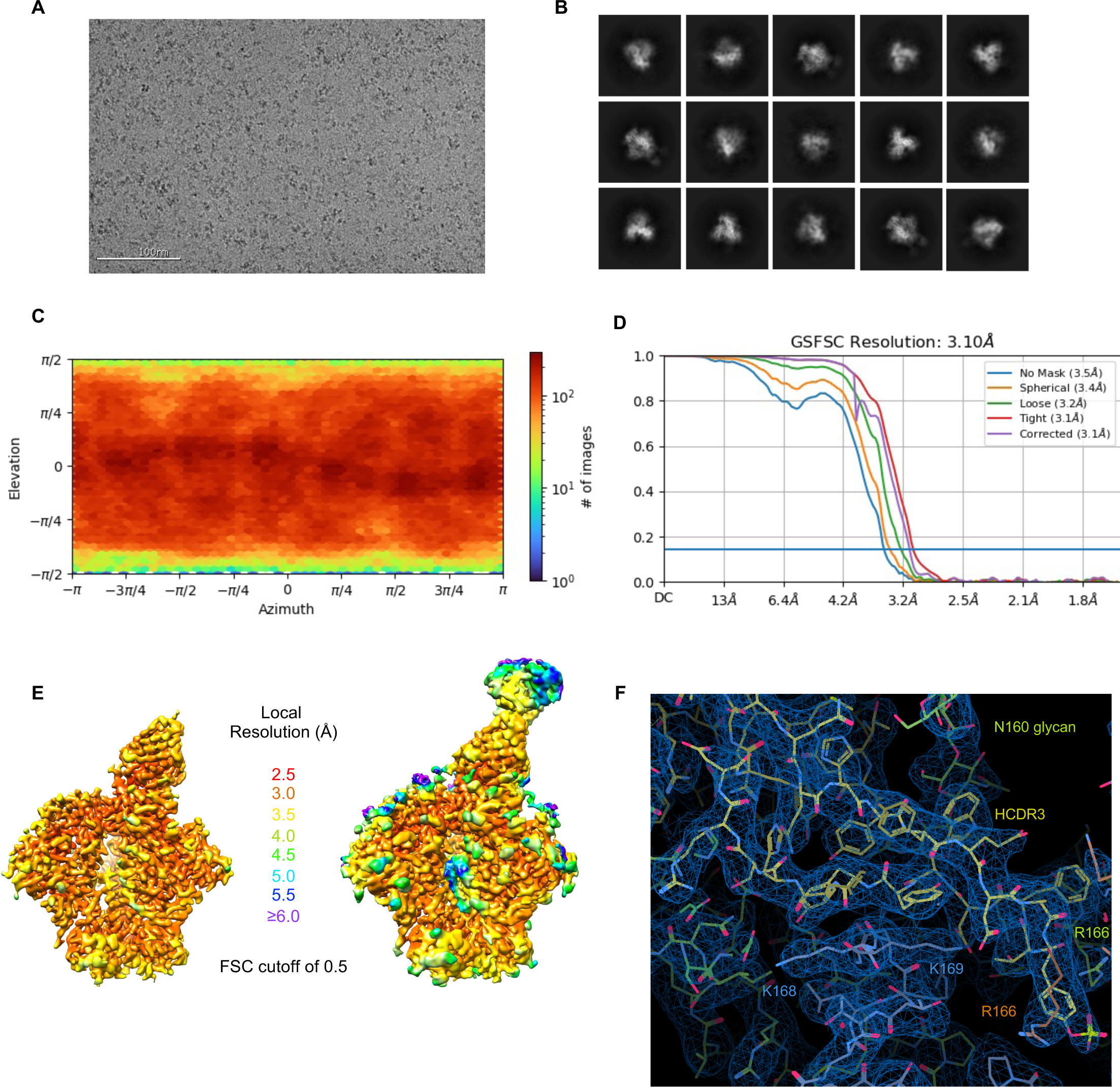
Cryo-EM details of V031-a.01 in complex with BG505 DS-SOSIP. (A) Representative raw micrograph is shown. (B) Representative 2D class averages of pick particles are shown. (C) The orientations of all particles used in the final refinement are shown as a heatmap. (D) The gold-standard fourier shell correlation (FSC) at threshold of 0.143 resulted in a resolution of 4.22 Å using non-uniform refinement with C1 symmetry. (E) The local resolution of the full map is shown generated through cryoSPARC using an FSC cutoff of 0.5. Two volume contour levels are shown. (F) Cryo-EM 3D reconstruction density to highlight the Fab-trimer interactive surface.

**Dataset S1 Figure 8.**
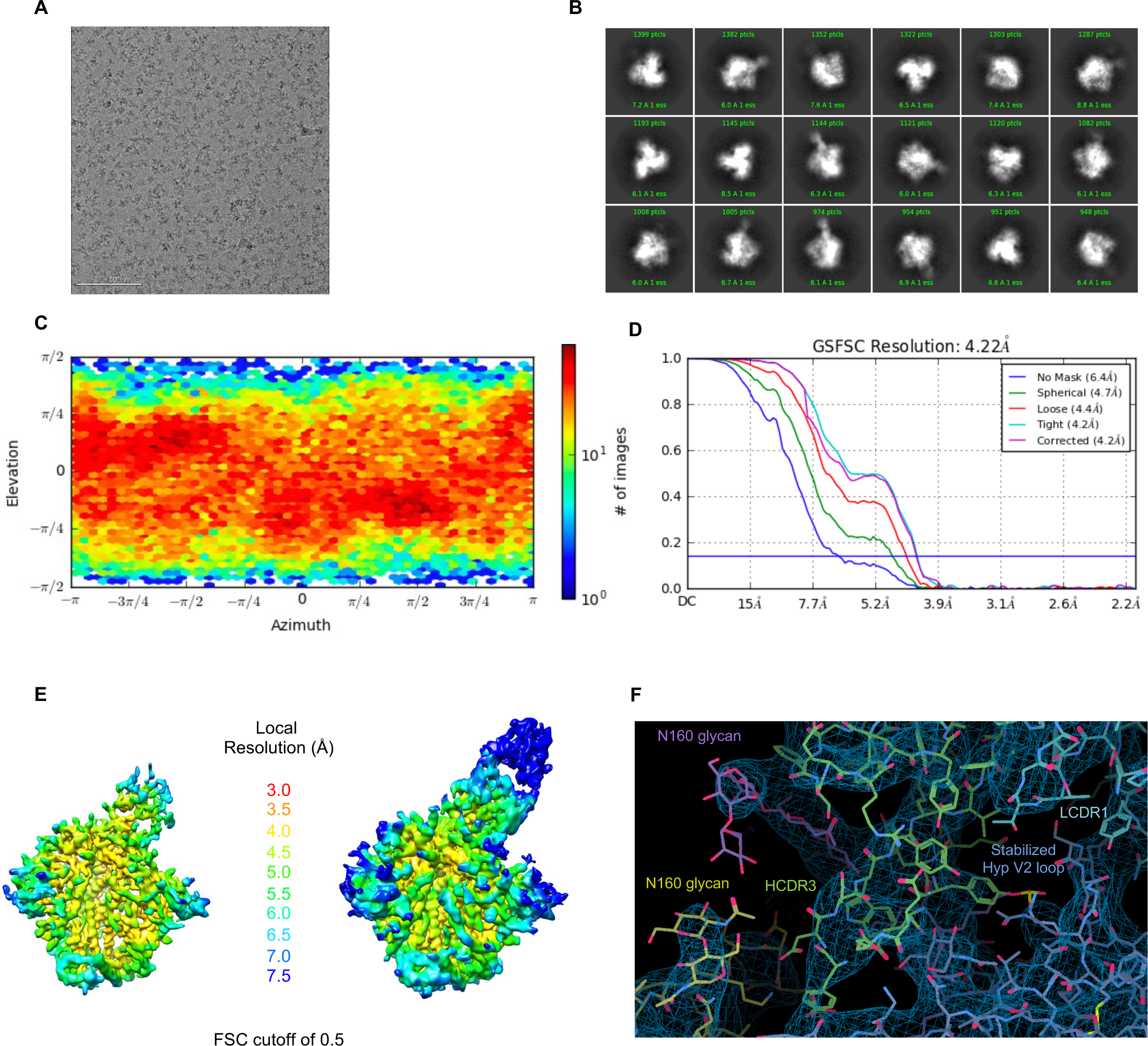
Cryo-EM details of 6561-a.01 in complex with Ce1176 RnS2 SOSIP. (A) Representative raw micrograph is shown. (B) Representative 2D class averages of pick particles are shown. (C) The orientations of all particles used in the final refinement are shown as a heatmap. (D) The gold-standard fourier shell correlation (FSC) at threshold of 0.143 resulted in a resolution of 4.22 Å using non-uniform refinement with C1 symmetry. (E) The local resolution of the full map is shown generated through cryoSPARC using an FSC cutoff of 0.5. Two volume contour levels are shown. (F) Cryo-EM 3D reconstruction density to highlight the Fab-trimer interactive surface.

**Dataset S1 Figure 9.**
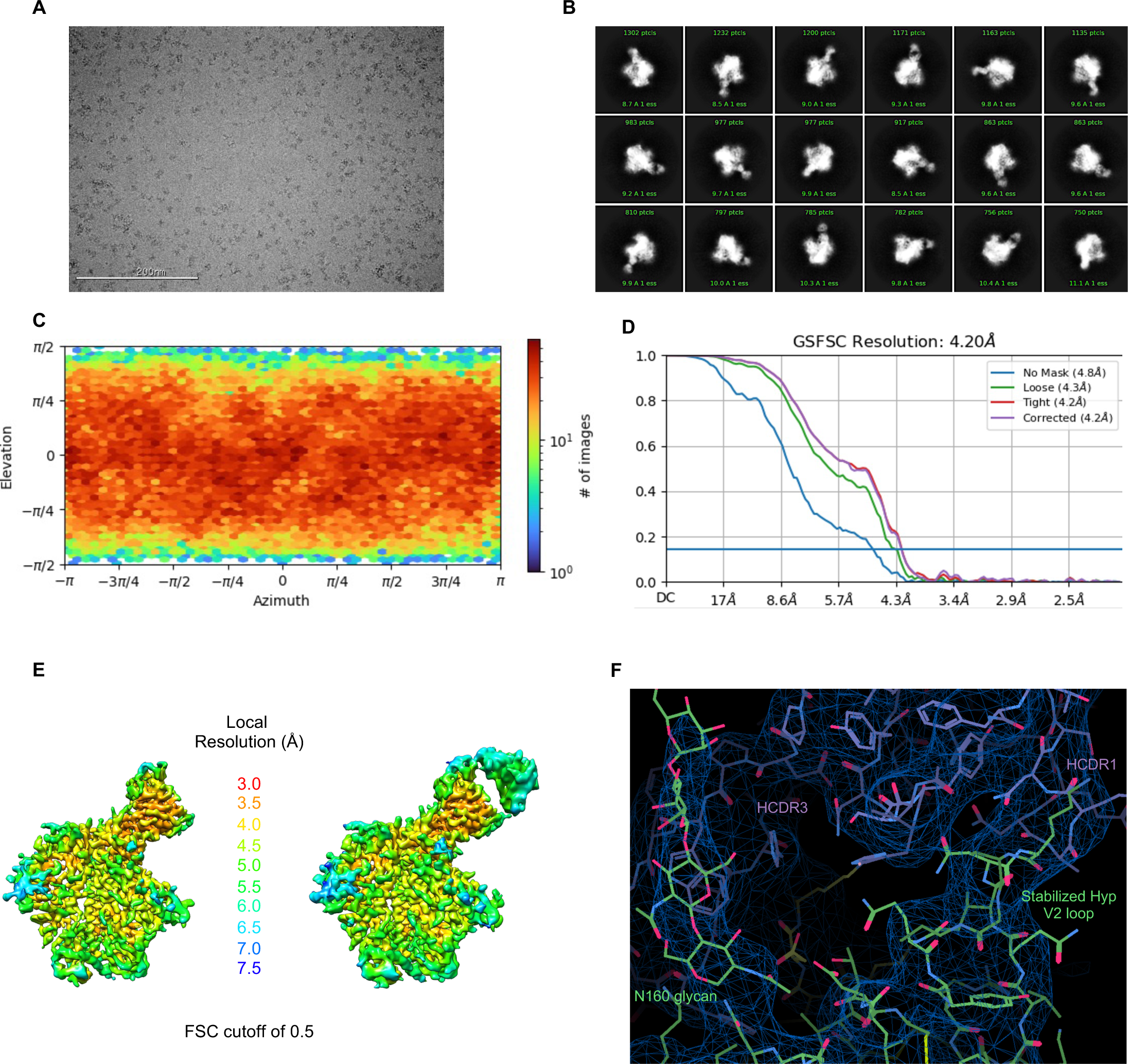
Cryo-EM details of 40591-a.01 in complex with T250.4 RnS2 SOSIP. (A) Representative raw micrograph is shown. (B) Representative 2D class averages of pick particles are shown. (C) The orientations of all particles used in the final refinement are shown as a heatmap. (D) The gold-standard fourier shell correlation (FSC) at threshold of 0.143 resulted in a resolution of 4.14 Å using non-uniform refinement with C1 symmetry. (E) The local resolution of the full map is shown generated through cryoSPARC using an FSC cutoff of 0.5. Two volume contour levels are shown. (F) Cryo-EM 3D reconstruction density to highlight the Fab-trimer interactive surface.

## REFERENCES

1. Chuang, G.Y., Zhou, J., Acharya, P., Rawi, R., Shen, C.H., Sheng, Z., Zhang, B., Zhou, T., Bailer, R.T., Dandey, V.P., et al. (2019). Structural Survey of Broadly Neutralizing Antibodies Targeting the HIV-1 Env Trimer Delineates Epitope Categories and Characteristics of Recognition. Structure 27, 196–206 e196. 10.1016/j.str.2018.10.007.

2. Ditse, Z., Muenchhoff, M., Adland, E., Jooste, P., Goulder, P., Moore, P.L., and Morris, L. (2018). HIV-1 Subtype C-Infected Children with Exceptional Neutralization Breadth Exhibit Polyclonal Responses Targeting Known Epitopes. J Virol 92. 10.1128/JVI.00878-18.

3. Landais, E., Huang, X., Havenar-Daughton, C., Murrell, B., Price, M.A., Wickramasinghe, L., Ramos, A., Bian, C.B., Simek, M., Allen, S., et al. (2016). Broadly Neutralizing Antibody Responses in a Large Longitudinal Sub-Saharan HIV Primary Infection Cohort. PLoS Pathog 12, e1005369. 10.1371/journal.ppat.1005369.

4. McLellan, J.S., Pancera, M., Carrico, C., Gorman, J., Julien, J.P., Khayat, R., Louder, R., Pejchal, R., Sastry, M., Dai, K., et al. (2011). Structure of HIV-1 gp120 V1/V2 domain with broadly neutralizing antibody PG9. Nature 480, 336–343. 10.1038/nature10696.

5. Walker, L.M., Simek, M.D., Priddy, F., Gach, J.S., Wagner, D., Zwick, M.B., Phogat, S.K., Poignard, P., and Burton, D.R. (2010). A limited number of antibody specificities mediate broad and potent serum neutralization in selected HIV-1 infected individuals. PLoS Pathog 6, e1001028. 10.1371/journal.ppat.1001028.

6. Andrabi, R., Voss, J.E., Liang, C.H., Briney, B., McCoy, L.E., Wu, C.Y., Wong, C.H., Poignard, P., and Burton, D.R. (2015). Identification of Common Features in Prototype Broadly Neutralizing Antibodies to HIV Envelope V2 Apex to Facilitate Vaccine Design. Immunity 43, 959–973. 10.1016/j.immuni.2015.10.014.

7. Gorman, J., Soto, C., Yang, M.M., Davenport, T.M., Guttman, M., Bailer, R.T., Chambers, M., Chuang, G.Y., DeKosky, B.J., Doria-Rose, N.A., et al. (2016). Structures of HIV-1 Env V1V2 with broadly neutralizing antibodies reveal commonalities that enable vaccine design. Nat Struct Mol Biol 23, 81–90. 10.1038/nsmb.3144.

8. Doria-Rose, N.A., Schramm, C.A., Gorman, J., Moore, P.L., Bhiman, J.N., DeKosky, B.J., Ernandes, M.J., Georgiev, I.S., Kim, H.J., Pancera, M., et al. (2014). Developmental pathway for potent V1V2-directed HIV-neutralizing antibodies. Nature 509, 55–62. 10.1038/nature13036.

9. Landais, E., Murrell, B., Briney, B., Murrell, S., Rantalainen, K., Berndsen, Z.T., Ramos, A., Wickramasinghe, L., Smith, M.L., Eren, K., et al. (2017). HIV Envelope Glycoform Heterogeneity and Localized Diversity Govern the Initiation and Maturation of a V2 Apex Broadly Neutralizing Antibody Lineage. Immunity 47, 990–1003 e1009. 10.1016/j.immuni.2017.11.002.

10. Willis, J.R., Finn, J.A., Briney, B., Sapparapu, G., Singh, V., King, H., LaBranche, C.C., Montefiori, D.C., Meiler, J., and Crowe, J.E., Jr. (2016). Long antibody HCDR3s from HIV-naive donors presented on a PG9 neutralizing antibody background mediate HIV neutralization. Proc Natl Acad Sci U S A 113, 4446–4451. 10.1073/pnas.1518405113.

11. Briney, B.S., Willis, J.R., and Crowe, J.E., Jr. (2012). Human peripheral blood antibodies with long HCDR3s are established primarily at original recombination using a limited subset of germline genes. PLoS One 7, e36750. 10.1371/journal.pone.0036750.

12. Willis, J.R., Berndsen, Z.T., Ma, K.M., Steichen, J.M., Schiffner, T., Landais, E., Liguori, A., Kalyuzhniy, O., Allen, J.D., Baboo, S., et al. (2022). Human immunoglobulin repertoire analysis guides design of vaccine priming immunogens targeting HIV V2-apex broadly neutralizing antibody precursors. Immunity 55, 2149–2167 e2149. 10.1016/j.immuni.2022.09.001.

13. Morgan, C., Marthas, M., Miller, C., Duerr, A., Cheng-Mayer, C., Desrosiers, R., Flores, J., Haigwood, N., Hu, S.L., Johnson, R.P., et al. (2008). The use of nonhuman primate models in HIV vaccine development. PLoS Med 5, e173. 10.1371/journal.pmed.0050173.

14. Hu, J.K., Crampton, J.C., Cupo, A., Ketas, T., van Gils, M.J., Sliepen, K., de Taeye, S.W., Sok, D., Ozorowski, G., Deresa, I., et al. (2015). Murine Antibody Responses to Cleaved Soluble HIV-1 Envelope Trimers Are Highly Restricted in Specificity. J Virol 89, 10383–10398. 10.1128/JVI.01653-15.

15. Francica, J.R., Sheng, Z., Zhang, Z., Nishimura, Y., Shingai, M., Ramesh, A., Keele, B.F., Schmidt, S.D., Flynn, B.J., Darko, S., et al. (2015). Analysis of immunoglobulin transcripts and hypermutation following SHIV(AD8) infection and protein-plus-adjuvant immunization. Nat Commun 6, 6565. 10.1038/ncomms7565.

16. Sundling, C., Li, Y., Huynh, N., Poulsen, C., Wilson, R., O’Dell, S., Feng, Y., Mascola, J.R., Wyatt, R.T., and Karlsson Hedestam, G.B. (2012). High-resolution definition of vaccine-elicited B cell responses against the HIV primary receptor binding site. Sci Transl Med 4, 142ra196. 10.1126/scitranslmed.3003752.

17. Roark, R.S., Li, H., Williams, W.B., Chug, H., Mason, R.D., Gorman, J., Wang, S., Lee, F.H., Rando, J., Bonsignori, M., et al. (2021). Recapitulation of HIV-1 Env-antibody coevolution in macaques leading to neutralization breadth. Science 371. 10.1126/science.abd2638.

18. Lee, J.H., Andrabi, R., Su, C.Y., Yasmeen, A., Julien, J.P., Kong, L., Wu, N.C., McBride, R., Sok, D., Pauthner, M., et al. (2017). A Broadly Neutralizing Antibody Targets the Dynamic HIV Envelope Trimer Apex via a Long, Rigidified, and Anionic beta-Hairpin Structure. Immunity 46, 690–702. 10.1016/j.immuni.2017.03.017.

19. Mason, R.D., Welles, H.C., Adams, C., Chakrabarti, B.K., Gorman, J., Zhou, T., Nguyen, R., O’Dell, S., Lusvarghi, S., Bewley, C.A., et al. (2016). Targeted Isolation of Antibodies Directed against Major Sites of SIV Env Vulnerability. PLoS Pathog 12, e1005537. 10.1371/journal.ppat.1005537.

20. Wiehe, K., Easterhoff, D., Luo, K., Nicely, N.I., Bradley, T., Jaeger, F.H., Dennison, S.M., Zhang, R., Lloyd, K.E., Stolarchuk, C., et al. (2014). Antibody light-chain-restricted recognition of the site of immune pressure in the RV144 HIV-1 vaccine trial is phylogenetically conserved. Immunity 41, 909–918. 10.1016/j.immuni.2014.11.014.

21. Moore, P.L., Gorman, J., Doria-Rose, N.A., and Morris, L. (2017). Ontogeny-based immunogens for the induction of V2-directed HIV broadly neutralizing antibodies. Immunol Rev 275, 217–229. 10.1111/imr.12501.

22. Corcoran, M.M., Phad, G.E., Vazquez Bernat, N., Stahl-Hennig, C., Sumida, N., Persson, M.A., Martin, M., and Karlsson Hedestam, G.B. (2016). Production of individualized V gene databases reveals high levels of immunoglobulin genetic diversity. Nat Commun 7, 13642. 10.1038/ncomms13642.

23. Walker, L.M., Phogat, S.K., Chan-Hui, P.Y., Wagner, D., Phung, P., Goss, J.L., Wrin, T., Simek, M.D., Fling, S., Mitcham, J.L., et al. (2009). Broad and potent neutralizing antibodies from an African donor reveal a new HIV-1 vaccine target. Science 326, 285–289. 10.1126/science.1178746.

24. Bonsignori, M., Hwang, K.K., Chen, X., Tsao, C.Y., Morris, L., Gray, E., Marshall, D.J., Crump, J.A., Kapiga, S.H., Sam, N.E., et al. (2011). Analysis of a clonal lineage of HIV-1 envelope V2/V3 conformational epitope-specific broadly neutralizing antibodies and their inferred unmutated common ancestors. J Virol 85, 9998–10009. 10.1128/JVI.05045-11.

25. Walker, L.M., Huber, M., Doores, K.J., Falkowska, E., Pejchal, R., Julien, J.P., Wang, S.K., Ramos, A., Chan-Hui, P.Y., Moyle, M., et al. (2011). Broad neutralization coverage of HIV by multiple highly potent antibodies. Nature 477, 466–470. 10.1038/nature10373.

26. Moore, P.L., and Williamson, C. (2016). Approaches to the induction of HIV broadly neutralizing antibodies. Curr Opin HIV AIDS 11, 569–575. 10.1097/COH.0000000000000317.

27. Burton, D.R., and Hangartner, L. (2016). Broadly Neutralizing Antibodies to HIV and Their Role in Vaccine Design. Annu Rev Immunol 34, 635–659. 10.1146/annurev-immunol-041015-055515.

28. Griffith, S.A., and McCoy, L.E. (2021). To bnAb or Not to bnAb: Defining Broadly Neutralising Antibodies Against HIV-1. Front Immunol 12, 708227. 10.3389/fimmu.2021.708227.

29. Kwong, P.D., and Mascola, J.R. (2018). HIV-1 Vaccines Based on Antibody Identification, B Cell Ontogeny, and Epitope Structure. Immunity 48, 855–871. 10.1016/j.immuni.2018.04.029.

30. Doria-Rose, N.A., Bhiman, J.N., Roark, R.S., Schramm, C.A., Gorman, J., Chuang, G.Y., Pancera, M., Cale, E.M., Ernandes, M.J., Louder, M.K., et al. (2016). New Member of the V1V2-Directed CAP256-VRC26 Lineage That Shows Increased Breadth and Exceptional Potency. J Virol 90, 76–91. 10.1128/JVI.01791-15.

31. Ramesh, A., Darko, S., Hua, A., Overman, G., Ransier, A., Francica, J.R., Trama, A., Tomaras, G.D., Haynes, B.F., Douek, D.C., and Kepler, T.B. (2017). Structure and Diversity of the Rhesus Macaque Immunoglobulin Loci through Multiple De Novo Genome Assemblies. Front Immunol 8, 1407. 10.3389/fimmu.2017.01407.

32. Vazquez Bernat, N., Corcoran, M., Nowak, I., Kaduk, M., Castro Dopico, X., Narang, S., Maisonasse, P., Dereuddre-Bosquet, N., Murrell, B., and Karlsson Hedestam, G.B. (2021). Rhesus and cynomolgus macaque immunoglobulin heavy-chain genotyping yields comprehensive databases of germline VDJ alleles. Immunity 54, 355–366 e354. 10.1016/j.immuni.2020.12.018.

33. Gao, N., Gai, Y., Meng, L., Wang, C., Wang, W., Li, X., Gu, T., Louder, M.K., Doria-Rose, N.A., Wiehe, K., et al. (2022). Development of Neutralization Breadth against Diverse HIV-1 by Increasing Ab-Ag Interface on V2. Adv Sci (Weinh) 9, e2200063. 10.1002/advs.202200063.

34. Rantalainen, K., Berndsen, Z.T., Murrell, S., Cao, L., Omorodion, O., Torres, J.L., Wu, M., Umotoy, J., Copps, J., Poignard, P., et al. (2018). Co-evolution of HIV Envelope and Apex-Targeting Neutralizing Antibody Lineage Provides Benchmarks for Vaccine Design. Cell Rep 23, 3249–3261. 10.1016/j.celrep.2018.05.046.

35. Gorman, J., Chuang, G.Y., Lai, Y.T., Shen, C.H., Boyington, J.C., Druz, A., Geng, H., Louder, M.K., McKee, K., Rawi, R., et al. (2020). Structure of Super-Potent Antibody CAP256-VRC26.25 in Complex with HIV-1 Envelope Reveals a Combined Mode of Trimer-Apex Recognition. Cell Rep 31, 107488. 10.1016/j.celrep.2020.03.052.

36. Huttner, W.B. (1982). Sulphation of tyrosine residues-a widespread modification of proteins. Nature 299, 273–276. 10.1038/299273a0.

37. Moore, K.L. (2003). The biology and enzymology of protein tyrosine O-sulfation. J Biol Chem 278, 24243–24246. 10.1074/jbc.R300008200.

38. Monigatti, F., Gasteiger, E., Bairoch, A., and Jung, E. (2002). The Sulfinator: predicting tyrosine sulfation sites in protein sequences. Bioinformatics 18, 769–770. 10.1093/bioinformatics/18.5.769.

39. Cai, C.X., Doria-Rose, N.A., Schneck, N.A., Ivleva, V.B., Tippett, B., Shadrick, W.R., O’Connell, S., Cooper, J.W., Schneiderman, Z., Zhang, B., et al. (2022). Tyrosine O-sulfation proteoforms affect HIV-1 monoclonal antibody potency. Sci Rep 12, 8433. 10.1038/s41598-022-12423-x.

40. Pejchal, R., Walker, L.M., Stanfield, R.L., Phogat, S.K., Koff, W.C., Poignard, P., Burton, D.R., and Wilson, I.A. (2010). Structure and function of broadly reactive antibody PG16 reveal an H3 subdomain that mediates potent neutralization of HIV-1. Proc Natl Acad Sci U S A 107, 11483–11488. 10.1073/pnas.1004600107.

41. Pan, Z., Liu, Z., Cheng, H., Wang, Y., Gao, T., Ullah, S., Ren, J., and Xue, Y. (2014). Systematic analysis of the in situ crosstalk of tyrosine modifications reveals no additional natural selection on multiply modified residues. Sci Rep 4, 7331. 10.1038/srep07331.

42. Kwong, P.D., and Mascola, J.R. (2012). Human antibodies that neutralize HIV-1: identification, structures, and B cell ontogenies. Immunity 37, 412–425. 10.1016/j.immuni.2012.08.012.

43. Wu, X., Yang, Z.Y., Li, Y., Hogerkorp, C.M., Schief, W.R., Seaman, M.S., Zhou, T., Schmidt, S.D., Wu, L., Xu, L., et al. (2010). Rational design of envelope identifies broadly neutralizing human monoclonal antibodies to HIV-1. Science 329, 856–861. 10.1126/science.1187659.

44. Zhou, T., Zhu, J., Wu, X., Moquin, S., Zhang, B., Acharya, P., Georgiev, I.S., Altae-Tran, H.R., Chuang, G.Y., Joyce, M.G., et al. (2013). Multidonor analysis reveals structural elements, genetic determinants, and maturation pathway for HIV-1 neutralization by VRC01-class antibodies. Immunity 39, 245–258. 10.1016/j.immuni.2013.04.012.

45. Wu, X., Zhou, T., Zhu, J., Zhang, B., Georgiev, I., Wang, C., Chen, X., Longo, N.S., Louder, M., McKee, K., et al. (2011). Focused evolution of HIV-1 neutralizing antibodies revealed by structures and deep sequencing. Science 333, 1593–1602. 10.1126/science.1207532.

46. Huang, D., Abbott, R.K., Havenar-Daughton, C., Skog, P.D., Al-Kolla, R., Groschel, B., Blane, T.R., Menis, S., Tran, J.T., Thinnes, T.C., et al. (2020). B cells expressing authentic naive human VRC01-class BCRs can be recruited to germinal centers and affinity mature in multiple independent mouse models. Proc Natl Acad Sci U S A 117, 22920–22931. 10.1073/pnas.2004489117.

47. Luo, S., Jing, C., Ye, A.Y., Kratochvil, S., Cottrell, C.A., Koo, J.H., Chapdelaine Williams, A., Francisco, L.V., Batra, H., Lamperti, E., et al. (2023). Humanized V(D)J-rearranging and TdT-expressing mouse vaccine models with physiological HIV-1 broadly neutralizing antibody precursors. Proc Natl Acad Sci U S A 120, e2217883120. 10.1073/pnas.2217883120.

48. Tian, M., Cheng, C., Chen, X., Duan, H., Cheng, H.L., Dao, M., Sheng, Z., Kimble, M., Wang, L., Lin, S., et al. (2016). Induction of HIV Neutralizing Antibody Lineages in Mice with Diverse Precursor Repertoires. Cell 166, 1471–1484 e1418. 10.1016/j.cell.2016.07.029.

49. Duan, H., Chen, X., Boyington, J.C., Cheng, C., Zhang, Y., Jafari, A.J., Stephens, T., Tsybovsky, Y., Kalyuzhniy, O., Zhao, P., et al. (2018). Glycan Masking Focuses Immune Responses to the HIV-1 CD4-Binding Site and Enhances Elicitation of VRC01-Class Precursor Antibodies. Immunity 49, 301–311 e305. 10.1016/j.immuni.2018.07.005.

50. Lee, J.H., Nakao, C., Appel, M., Le, A., Landais, E., Kalyuzhniy, O., Hu, X., Liguori, A., Mullen, T.M., Groschel, B., et al. (2022). Highly mutated antibodies capable of neutralizing N276 glycan-deficient HIV after a single immunization with an Env trimer. Cell Rep 38, 110485. 10.1016/j.celrep.2022.110485.

51. Leggat, D.J., Cohen, K.W., Willis, J.R., Fulp, W.J., deCamp, A.C., Kalyuzhniy, O., Cottrell, C.A., Menis, S., Finak, G., Ballweber-Fleming, L., et al. (2022). Vaccination induces HIV broadly neutralizing antibody precursors in humans. Science 378, eadd6502. 10.1126/science.add6502.

52. Ekiert, D.C., Bhabha, G., Elsliger, M.A., Friesen, R.H., Jongeneelen, M., Throsby, M., Goudsmit, J., and Wilson, I.A. (2009). Antibody recognition of a highly conserved influenza virus epitope. Science 324, 246–251. 10.1126/science.1171491.

53. Dreyfus, C., Laursen, N.S., Kwaks, T., Zuijdgeest, D., Khayat, R., Ekiert, D.C., Lee, J.H., Metlagel, Z., Bujny, M.V., Jongeneelen, M., et al. (2012). Highly conserved protective epitopes on influenza B viruses. Science 337, 1343–1348. 10.1126/science.1222908.

54. Sui, J., Hwang, W.C., Perez, S., Wei, G., Aird, D., Chen, L.M., Santelli, E., Stec, B., Cadwell, G., Ali, M., et al. (2009). Structural and functional bases for broad-spectrum neutralization of avian and human influenza A viruses. Nat Struct Mol Biol 16, 265–273. 10.1038/nsmb.1566.

55. Huang, C.C., Venturi, M., Majeed, S., Moore, M.J., Phogat, S., Zhang, M.Y., Dimitrov, D.S., Hendrickson, W.A., Robinson, J., Sodroski, J., et al. (2004). Structural basis of tyrosine sulfation and VH-gene usage in antibodies that recognize the HIV type 1 coreceptor-binding site on gp120. Proc Natl Acad Sci U S A 101, 2706–2711. 10.1073/pnas.0308527100.

56. Huang, C.C., Lam, S.N., Acharya, P., Tang, M., Xiang, S.H., Hussan, S.S., Stanfield, R.L., Robinson, J., Sodroski, J., Wilson, I.A., et al. (2007). Structures of the CCR5 N terminus and of a tyrosine-sulfated antibody with HIV-1 gp120 and CD4. Science 317, 1930–1934. 10.1126/science.1145373.

57. Choe, H., Li, W., Wright, P.L., Vasilieva, N., Venturi, M., Huang, C.C., Grundner, C., Dorfman, T., Zwick, M.B., Wang, L., et al. (2003). Tyrosine sulfation of human antibodies contributes to recognition of the CCR5 binding region of HIV-1 gp120. Cell 114, 161–170. 10.1016/s0092-8674(03)00508-7.

58. West, A.P., Jr., Diskin, R., Nussenzweig, M.C., and Bjorkman, P.J. (2012). Structural basis for germ-line gene usage of a potent class of antibodies targeting the CD4-binding site of HIV-1 gp120. Proc Natl Acad Sci U S A 109, E2083–2090. 10.1073/pnas.1208984109.

59. Gao, N., Wang, W., Wang, C., Gu, T., Guo, R., Yu, B., Kong, W., Qin, C., Giorgi, E.E., Chen, Z., et al. (2018). Development of broad neutralization activity in simian/human immunodeficiency virus-infected rhesus macaques after long-term infection. AIDS 32, 555–563. 10.1097/QAD.0000000000001724.

60. Andrabi, R., Pallesen, J., Allen, J.D., Song, G., Zhang, J., de Val, N., Gegg, G., Porter, K., Su, C.Y., Pauthner, M., et al. (2019). The Chimpanzee SIV Envelope Trimer: Structure and Deployment as an HIV Vaccine Template. Cell Rep 27, 2426–2441 e2426. 10.1016/j.celrep.2019.04.082.

61. Bibollet-Ruche, F., Russell, R.M., Ding, W., Liu, W., Li, Y., Wagh, K., Wrapp, D., Habib, R., Skelly, A.N., Roark, R.S., et al. (2023). A Germline-Targeting Chimpanzee SIV Envelope Glycoprotein Elicits a New Class of V2-Apex Directed Cross-Neutralizing Antibodies. mBio 14, e0337022. 10.1128/mbio.03370-22.

62. Bricault, C.A., Yusim, K., Seaman, M.S., Yoon, H., Theiler, J., Giorgi, E.E., Wagh, K., Theiler, M., Hraber, P., Macke, J.P., et al. (2019). HIV-1 Neutralizing Antibody Signatures and Application to Epitope-Targeted Vaccine Design. Cell Host Microbe 25, 59–72 e58. 10.1016/j.chom.2018.12.001.

63. Jardine, J., Julien, J.P., Menis, S., Ota, T., Kalyuzhniy, O., McGuire, A., Sok, D., Huang, P.S., MacPherson, S., Jones, M., et al. (2013). Rational HIV immunogen design to target specific germline B cell receptors. Science 340, 711–716. 10.1126/science.1234150.

64. Steichen, J.M., Lin, Y.C., Havenar-Daughton, C., Pecetta, S., Ozorowski, G., Willis, J.R., Toy, L., Sok, D., Liguori, A., Kratochvil, S., et al. (2019). A generalized HIV vaccine design strategy for priming of broadly neutralizing antibody responses. Science 366. 10.1126/science.aax4380.

65. Steichen, J.M., Kulp, D.W., Tokatlian, T., Escolano, A., Dosenovic, P., Stanfield, R.L., McCoy, L.E., Ozorowski, G., Hu, X., Kalyuzhniy, O., et al. (2016). HIV Vaccine Design to Target Germline Precursors of Glycan-Dependent Broadly Neutralizing Antibodies. Immunity 45, 483–496. 10.1016/j.immuni.2016.08.016.

66. Jardine, J.G., Kulp, D.W., Havenar-Daughton, C., Sarkar, A., Briney, B., Sok, D., Sesterhenn, F., Ereno-Orbea, J., Kalyuzhniy, O., Deresa, I., et al. (2016). HIV-1 broadly neutralizing antibody precursor B cells revealed by germline-targeting immunogen. Science 351, 1458–1463. 10.1126/science.aad9195.

67. Melzi, E., Willis, J.R., Ma, K.M., Lin, Y.C., Kratochvil, S., Berndsen, Z.T., Landais, E.A., Kalyuzhniy, O., Nair, U., Warner, J., et al. (2022). Membrane-bound mRNA immunogens lower the threshold to activate HIV Env V2 apex-directed broadly neutralizing B cell precursors in humanized mice. Immunity 55, 2168–2186 e2166. 10.1016/j.immuni.2022.09.003.

68. Caniels, T.G., Medina-Ramirez, M., Zhang, J., Sarkar, A., Kumar, S., LaBranche, A., Derking, R., Allen, J.D., Snitselaar, J.L., Capella-Pujol, J., et al. (2023). Germline-targeting HIV-1 Env vaccination induces VRC01-class antibodies with rare insertions. Cell Rep Med 4, 101003. 10.1016/j.xcrm.2023.101003.

69. Haynes, B.F., Wiehe, K., Borrow, P., Saunders, K.O., Korber, B., Wagh, K., McMichael, A.J., Kelsoe, G., Hahn, B.H., Alt, F., and Shaw, G.M. (2023). Strategies for HIV-1 vaccines that induce broadly neutralizing antibodies. Nat Rev Immunol 23, 142–158. 10.1038/s41577-022-00753-w.

70. Wiehe, K., Saunders, K.O., Stalls, V., Cain, D.W., Venkatayogi, S., Martin Beem, J.S., Berry, M., Evangelous, T., Henderson, R., Hora, B., et al. (2024). Mutation-guided vaccine design: A process for developing boosting immunogens for HIV broadly neutralizing antibody induction. Cell Host Microbe. 10.1016/j.chom.2024.04.006.

71. Saunders, K.O., Wiehe, K., Tian, M., Acharya, P., Bradley, T., Alam, S.M., Go, E.P., Scearce, R., Sutherland, L., Henderson, R., et al. (2019). Targeted selection of HIV-specific antibody mutations by engineering B cell maturation. Science 366. 10.1126/science.aay7199.

72. Haynes, B.F., Kelsoe, G., Harrison, S.C., and Kepler, T.B. (2012). B-cell-lineage immunogen design in vaccine development with HIV-1 as a case study. Nat Biotechnol 30, 423–433. 10.1038/nbt.2197.

73. Barbian, H.J., Decker, J.M., Bibollet-Ruche, F., Galimidi, R.P., West, A.P., Jr., Learn, G.H., Parrish, N.F., Iyer, S.S., Li, Y., Pace, C.S., et al. (2015). Neutralization properties of simian immunodeficiency viruses infecting chimpanzees and gorillas. mBio 6. 10.1128/mBio.00296-15.

74. Yoon, H., Macke, J., West, A.P., Jr., Foley, B., Bjorkman, P.J., Korber, B., and Yusim, K. (2015). CATNAP: a tool to compile, analyze and tally neutralizing antibody panels. Nucleic Acids Res 43, W213–219. 10.1093/nar/gkv404.

75. Li, H., Wang, S., Lee, F.H., Roark, R.S., Murphy, A.I., Smith, J., Zhao, C., Rando, J., Chohan, N., Ding, Y., et al. (2021). New SHIVs and Improved Design Strategy for Modeling HIV-1 Transmission, Immunopathogenesis, Prevention and Cure. J Virol 95. 10.1128/JVI.00071-21.

76. Bauer, A., Lindemuth, E., Marino, F.E., Krause, R., Joy, J., Docken, S.S., Mallick, S., McCormick, K., Holt, C., Georgiev, I., et al. (2023). Adaptation of a transmitted/founder simian-human immunodeficiency virus for enhanced replication in rhesus macaques. PLoS Pathog 19, e1011059. 10.1371/journal.ppat.1011059.

77. Williams, W.B., Zhang, J., Jiang, C., Nicely, N.I., Fera, D., Luo, K., Moody, M.A., Liao, H.X., Alam, S.M., Kepler, T.B., et al. (2017). Initiation of HIV neutralizing B cell lineages with sequential envelope immunizations. Nat Commun 8, 1732. 10.1038/s41467-017-01336-3.

78. Han, Q., Bradley, T., Williams, W.B., Cain, D.W., Montefiori, D.C., Saunders, K.O., Parks, R.J., Edwards, R.W., Ferrari, G., Mueller, O., et al. (2020). Neonatal Rhesus Macaques Have Distinct Immune Cell Transcriptional Profiles following HIV Envelope Immunization. Cell Rep 30, 1553–1569 e1556. 10.1016/j.celrep.2019.12.091.

79. Ye, J., Ma, N., Madden, T.L., and Ostell, J.M. (2013). IgBLAST: an immunoglobulin variable domain sequence analysis tool. Nucleic Acids Res 41, W34–40. 10.1093/nar/gkt382.

80. Brochet, X., Lefranc, M.P., and Giudicelli, V. (2008). IMGT/V-QUEST: the highly customized and integrated system for IG and TR standardized V-J and V-D-J sequence analysis. Nucleic Acids Res 36, W503–508. 10.1093/nar/gkn316.

81. Krebs, S.J., Kwon, Y.D., Schramm, C.A., Law, W.H., Donofrio, G., Zhou, K.H., Gift, S., Dussupt, V., Georgiev, I.S., Schatzle, S., et al. (2019). Longitudinal Analysis Reveals Early Development of Three MPER-Directed Neutralizing Antibody Lineages from an HIV-1-Infected Individual. Immunity 50, 677–691 e613. 10.1016/j.immuni.2019.02.008.

82. Schramm, C.A., Sheng, Z., Zhang, Z., Mascola, J.R., Kwong, P.D., and Shapiro, L. (2016). SONAR: A High-Throughput Pipeline for Inferring Antibody Ontogenies from Longitudinal Sequencing of B Cell Transcripts. Front Immunol 7, 372. 10.3389/fimmu.2016.00372.

83. Bhardwaj, V., Franceschetti, M., Rao, R., Pevzner, P.A., and Safonova, Y. (2020). Automated analysis of immunosequencing datasets reveals novel immunoglobulin D genes across diverse species. PLoS Comput Biol 16, e1007837. 10.1371/journal.pcbi.1007837.

84. Sok, D., van Gils, M.J., Pauthner, M., Julien, J.P., Saye-Francisco, K.L., Hsueh, J., Briney, B., Lee, J.H., Le, K.M., Lee, P.S., et al. (2014). Recombinant HIV envelope trimer selects for quaternary-dependent antibodies targeting the trimer apex. Proc Natl Acad Sci U S A 111, 17624–17629. 10.1073/pnas.1415789111.

85. Kwon, Y.D., Pancera, M., Acharya, P., Georgiev, I.S., Crooks, E.T., Gorman, J., Joyce, M.G., Guttman, M., Ma, X., Narpala, S., et al. (2015). Crystal structure, conformational fixation and entry-related interactions of mature ligand-free HIV-1 Env. Nat Struct Mol Biol 22, 522–531. 10.1038/nsmb.3051.

86. Saunders, K.O., Verkoczy, L.K., Jiang, C., Zhang, J., Parks, R., Chen, H., Housman, M., Bouton-Verville, H., Shen, X., Trama, A.M., et al. (2017). Vaccine Induction of Heterologous Tier 2 HIV-1 Neutralizing Antibodies in Animal Models. Cell Rep 21, 3681–3690. 10.1016/j.celrep.2017.12.028.

87. Rawi, R., Rutten, L., Lai, Y.T., Olia, A.S., Blokland, S., Juraszek, J., Shen, C.H., Tsybovsky, Y., Verardi, R., Yang, Y., et al. (2020). Automated Design by Structure-Based Stabilization and Consensus Repair to Achieve Prefusion-Closed Envelope Trimers in a Wide Variety of HIV Strains. Cell Rep 33, 108432. 10.1016/j.celrep.2020.108432.

88. Wrapp, D., Mu, Z., Thakur, B., Janowska, K., Ajayi, O., Barr, M., Parks, R., Mansouri, K., Edwards, R.J., Hahn, B.H., et al. (2023). Structure-Based Stabilization of SOSIP Env Enhances Recombinant Ectodomain Durability and Yield. J Virol 97, e0167322. 10.1128/jvi.01673-22.

89. Suloway, C., Pulokas, J., Fellmann, D., Cheng, A., Guerra, F., Quispe, J., Stagg, S., Potter, C.S., and Carragher, B. (2005). Automated molecular microscopy: the new Leginon system. J Struct Biol 151, 41–60. 10.1016/j.jsb.2005.03.010.

90. Punjani, A., Rubinstein, J.L., Fleet, D.J., and Brubaker, M.A. (2017). cryoSPARC: algorithms for rapid unsupervised cryo-EM structure determination. Nat Methods 14, 290–296. 10.1038/nmeth.4169.

91. Dunbar, J., Krawczyk, K., Leem, J., Marks, C., Nowak, J., Regep, C., Georges, G., Kelm, S., Popovic, B., and Deane, C.M. (2016). SAbPred: a structure-based antibody prediction server. Nucleic Acids Res 44, W474–478. 10.1093/nar/gkw361.

92. Pettersen, E.F., Goddard, T.D., Huang, C.C., Couch, G.S., Greenblatt, D.M., Meng, E.C., and Ferrin, T.E. (2004). UCSF Chimera--a visualization system for exploratory research and analysis. J Comput Chem 25, 1605–1612. 10.1002/jcc.20084.

93. Meng, E.C., Goddard, T.D., Pettersen, E.F., Couch, G.S., Pearson, Z.J., Morris, J.H., and Ferrin, T.E. (2023). UCSF ChimeraX: Tools for structure building and analysis. Protein Sci 32, e4792. 10.1002/pro.4792.

94. Emsley, P., and Cowtan, K. (2004). Coot: model-building tools for molecular graphics. Acta Crystallogr D Biol Crystallogr 60, 2126–2132. 10.1107/S0907444904019158.

95. Liebschner, D., Afonine, P.V., Baker, M.L., Bunkoczi, G., Chen, V.B., Croll, T.I., Hintze, B., Hung, L.W., Jain, S., McCoy, A.J., et al. (2019). Macromolecular structure determination using X-rays, neutrons and electrons: recent developments in Phenix. Acta Crystallogr D Struct Biol 75, 861–877. 10.1107/S2059798319011471.

96. Chen, V.B., Arendall, W.B., 3rd, Headd, J.J., Keedy, D.A., Immormino, R.M., Kapral, G.J., Murray, L.W., Richardson, J.S., and Richardson, D.C. (2010). MolProbity: all-atom structure validation for macromolecular crystallography. Acta Crystallogr D Biol Crystallogr 66, 12–21. 10.1107/S0907444909042073.

97. Barad, B.A., Echols, N., Wang, R.Y., Cheng, Y., DiMaio, F., Adams, P.D., and Fraser, J.S. (2015). EMRinger: side chain-directed model and map validation for 3D cryo-electron microscopy. Nat Methods 12, 943–946. 10.1038/nmeth.3541.

98. Krissinel, E., and Henrick, K. (2007). Inference of macromolecular assemblies from crystalline state. J Mol Biol 372, 774–797. 10.1016/j.jmb.2007.05.022.

99. Sievers, F., Wilm, A., Dineen, D., Gibson, T.J., Karplus, K., Li, W., Lopez, R., McWilliam, H., Remmert, M., Soding, J., et al. (2011). Fast, scalable generation of high-quality protein multiple sequence alignments using Clustal Omega. Mol Syst Biol 7, 539. 10.1038/msb.2011.75.

